# Data-driven model of glycolysis shows that allostery maintains high ATP levels while mass action controls flux and energy of ATP hydrolysis

**DOI:** 10.1101/2022.12.28.522046

**Authors:** Mangyu Choe, Tal Einav, Rob Phillips, Denis V. Titov

**Affiliations:** Department of Nutritional Sciences and Toxicology, University of California; Berkeley, CA 94720, USA; Department of Molecular and Cell Biology, University of California; Berkeley, CA 94720, USA; Center for Computational Biology, University of California; Berkeley, CA 94720, USA; Division of Biology and Biological Engineering, California Institute of Technology; Pasadena, CA 91125, USA; Department of Physics, California Institute of Technology; Pasadena, CA 91125, USA; Basic Sciences Division and Computational Biology Program, Fred Hutchinson Cancer Research Center, Seattle, WA 98109, USA; Center for Vaccine Innovation, La Jolla Institute for Immunology, La Jolla, CA 92037, USA

## Abstract

Glycolysis is a conserved metabolic pathway that produces ATP and biosynthetic precursors. Here, we use mathematical modeling to investigate how the control of mammalian glycolytic enzymes through allostery and mass action accomplishes various tasks of ATP homeostasis, such as controlling the rate of ATP production, maintaining high and stable ATP levels, and ensuring that ATP hydrolysis generates a net excess of energy. Our model uses data-derived enzyme rate equations, recapitulates the key tasks of glycolytic ATP homeostasis, and accurately predicts absolute concentrations of glycolytic intermediates and isotope tracing kinetics in live cells. We find that allosteric regulation of hexokinase (HK) and phosphofructokinase (PFK) by ATP, ADP, inorganic phosphate and glucose-6-phosphate (G6P), the surplus of lower glycolysis enzymes, and a large non-adenine phosphate pool are essential to robustly maintain high ATP levels and to prevent uncontrolled accumulation of phosphorylated intermediates of upper glycolysis. Meanwhile, mass action alone is sufficient to control ATP production rate and maintain high energy of ATP hydrolysis. Our results suggest a revision of the textbook view that the function of allosteric regulation of HK, PFK and PK is to control the net flux through glycolysis in response to variable ATP demand.

## INTRODUCTION

Glycolysis is conserved across all domains of life. This key pathway harnesses the breakdown of glucose to produce energy in the form of ATP and precursors for the biosynthesis of amino acids, nucleotides, carbohydrates, and lipids. Glycolysis produces ATP both directly by a process referred to as fermentation or aerobic glycolysis (***Figure 1A***) and indirectly by producing pyruvate, which is used as a substrate for ATP generation by respiration. The net reaction of mammalian aerobic glycolysis converts extracellular glucose to extracellular lactic acid, while the only net intracellular reaction is the production of ATP from ADP and inorganic phosphate (***Figure 1A***). Aerobic glycolysis is the most active metabolic pathway in proliferating mammalian cells (an observation known as the Warburg Effect (***DeBerardinis and Chandel, 2020***)), able to satisfy all the ATP demand even in the absence of respiration (***King and Attardi, 1989; Titov et al., 2016***). The reliance of proliferating mammalian cells on aerobic glycolysis for ATP production and the lack of intracellular products except for ATP makes glycolysis a convenient self-contained system for studying ATP homeostasis.

**Figure 1.**
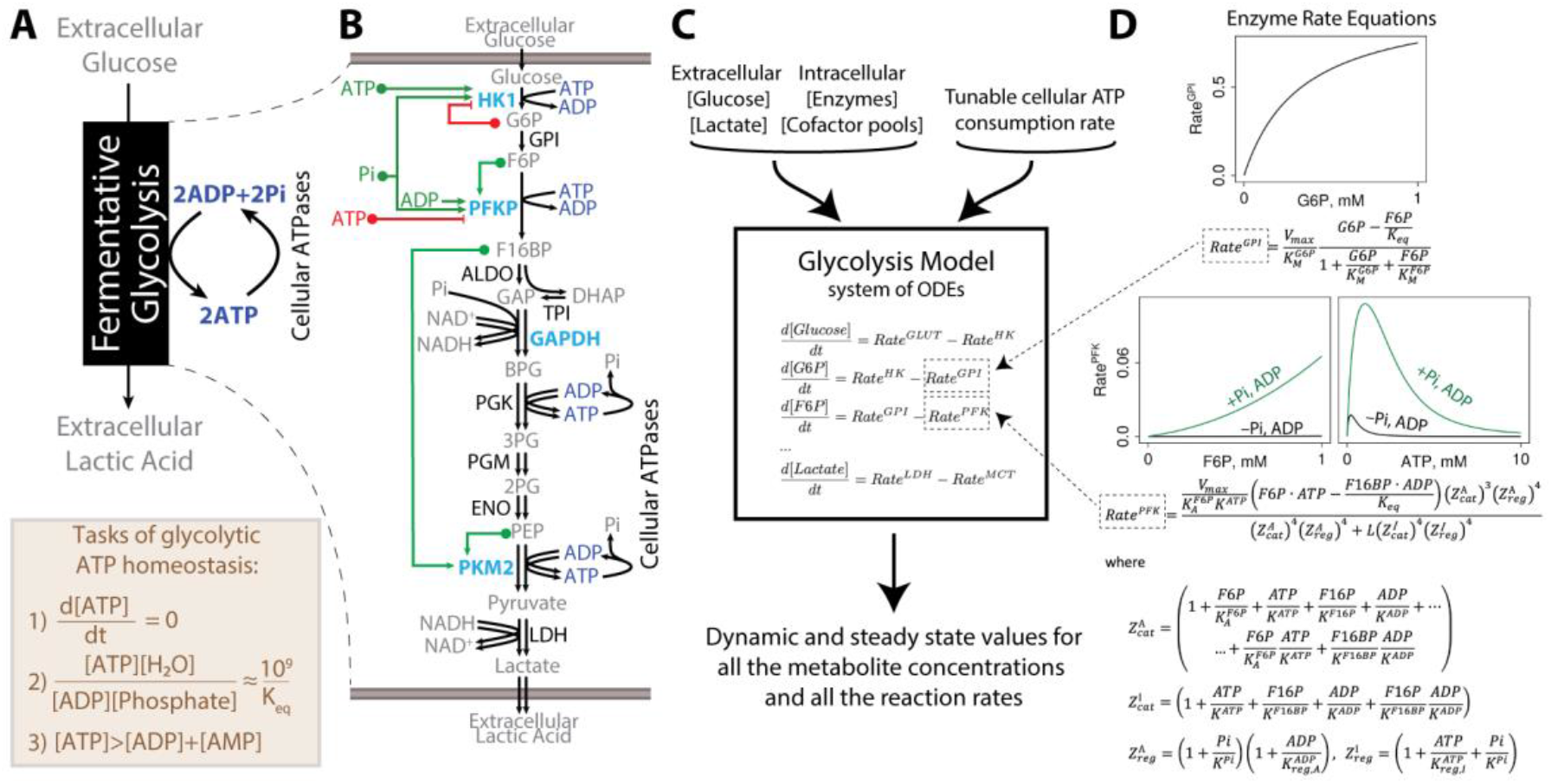
Overview of the mammalian glycolysis model. **(A)** Coarse-grained description of aerobic glycolysis highlighting its main function to transform glucose into ATP and listing the tasks of glycolytic ATP homeostasis. **(B)** Qualitative schematic of glycolysis showing the chain of enzymes (allosterically regulated enzymes in teal and the other enzymes in black) that convert substrates into products (gray). Allosteric activators (green) and inhibitors (red) that are included in the model are highlighted. **(C)** Schematic of the glycolysis model, including inputs and outputs. **(D)** Kinetic rate equations are shown for GPI and PFK, and plots are actual GPI and PFK rates calculated by the respective equations with rates normalized to V_max_. Note the dramatic allosteric activation of the PFK rate in the presence of inorganic phosphate (Pi) and ADP. See also ***Appendix 1*** for details on enzyme rate equation derivation.

To maintain ATP homeostasis, the glycolysis pathway must simultaneously achieve several tasks, such as regulating the rate of ATP production, ensuring that the ATP hydrolysis reaction generates a net excess of energy, maintaining most of the adenine nucleotide pool in the form of ATP, and balancing the activity of glycolysis with other metabolic pathways. These tasks must be achieved at a wide range of ATP demands and be robust to small variations in enzyme concentrations and kinetic parameters that are observed due to the stochasticity of gene expression or physiological changes in temperature or pH. For example, the robust control of glycolysis maintains near-constant ATP levels in response to a >100-fold change in ATP demand (***Hochachka and McClelland, 1997***).

Glycolytic ATP homeostasis is an emergent property of the activities of individual enzymes that are controlled by a combination of mass action and allosteric regulation. Mass action involves the regulation of enzyme rate in response to changes in substrate and product concentrations. Allosteric regulation (***Monod et al., 1965a; Koshland et al., 1966a; Phillips, 2020***) refers to the modulation of enzyme kinetic properties by metabolite binding to a non-catalytic site or posttranslational modifications, allowing for feedback regulation to achieve the desired properties of homeostasis by modifying the mass action trends of biochemical reactions (***Hofmeyr, 1995; Alon, 2019; El-Samad, 2021***). Decades of biochemical studies have uncovered that four glycolytic enzymes–hexokinase (HK), phosphofructokinase (PFK), glyceraldehyde-3-dehydrogenase (GAPDH), pyruvate kinase (PK)–are allosterically regulated by a constellation of metabolites like ATP, ADP, AMP, inorganic phosphate, and other regulators (***Figure 1A-B***). The textbook view (***Berg et al., 2015; Nelson, 2021***) is that the function of allosteric regulation of HK, PFK, and PK is to control ATP production rate or flux through glycolysis in response to ATP demand. For example, allosteric inhibition of PFK by ATP would inhibit flux through glycolysis when ATP level is high and activate it when ATP level is low. Yet, a simple mass action regulation could achieve the same effect as ATP is the final product of the pathway and its consumption should speed up the pathway through mass action alone. In fact, no experiments have assessed the impact of inactivating these allosteric interactions on the entire glycolytic pathway. This is largely due to the practical difficulty of simultaneously deleting or mutating a minimum of 10 genes encoding various human isoforms of HK, PFK, GAPDH, and PK. Therefore, the roles of mass action and allostery in the control of specific tasks of glycolytic ATP homeostasis are unknown.

Owing to the complexity of glycolytic regulation and the challenges associated with experimentally inactivating allostery, mathematical modeling has been employed to gain insights into the control of glycolytic ATP homeostasis. Theoretical analysis of metabolic regulation has been an active area of research for many decades with extensive work done on Metabolic Control Analysis (***Heinrich and Rapoport, 1974; Kacser and Burns, 1973; Reich and Sel’kov, 1974***), which applied derivative-based local sensitivity analysis techniques to studying the control of flux and concentrations by metabolic pathways. Several groups have developed mathematical models of glycolysis (***Sel’kov, 1975; Rapoport et al., 1976; Termonia and Ross, 1981; Joshi and Palsson, 1990; Teusink et al., 1996; Mulquiney and Kuchel, 1999; Teusink et al., 2000; Hynne et al., 2001; Chandra et al., 2011; Eunen et al., 2012; Smallbone et al., 2013; van Heerden et al., 2014; Shestov et al., 2014; Mulukutla et al., 2014***). These efforts ranged from simple idealized models involving a subset of enzymes (***Chandra et al., 2011; Rapoport et al., 1976; Sel’kov, 1975; Termonia and Ross, 1981; Teusink et al., 1996***) to full-scale models containing all the enzymes (***Eunen et al., 2012; Hynne et al., 2001; Joshi and Palsson, 1990; Mulquiney and Kuchel, 1999; Mulukutla et al., 2014; Shestov et al., 2014; Smallbone et al., 2013; Teusink et al., 2000; van Heerden et al., 2014***). Models of glycolysis were used to investigate the control of glycolytic flux and ATP levels (***Chandra et al., 2011; Rapoport et al., 1976; Sel’kov, 1975; Termonia and Ross, 1981; Teusink et al., 1996***), compare model prediction to data from live cells (***Eunen et al., 2012; Hynne et al., 2001; Joshi and Palsson, 1990; Mulquiney and Kuchel, 1999; Mulukutla et al., 2014; Shestov et al., 2014; Smallbone et al., 2013; Teusink et al., 2000; van Heerden et al., 2014***), and investigate the mechanisms leading to the occurrence of glycolytic oscillations (***Chandra et al., 2011; Sel’kov, 1975; Termonia and Ross, 1981; Teusink et al., 1996***). Published mathematical models of glycolysis have not assessed the impact of inactivating allosteric regulators on ATP homeostasis and used at least one simplifying condition–omitting inorganic phosphate, fixing ATP, ADP, or Pi levels, or simulating only one fixed rate of ATP demand–that makes it difficult to investigate the control of ATP homeostasis.

Here, we investigated the role of mass action and allostery in controlling various tasks of glycolytic ATP homeostasis using a combination of theory and experiments. The core of our approach is a mathematical model that uses enzyme rate equations to simulate glycolysis activity using a system of ordinary differential equations (ODEs). The key components of our model are the newly derived rate equations for allosterically regulated enzymes HK, PFK, GAPDH, and PK based on decades of *in vitro* kinetic data. To the best of our knowledge, we compiled the largest dataset of mammalian glycolytic enzyme kinetic data reported to date, comprising about 3000 measurements (***Supplementary File 2***). We used the Monod-Wyman-Changeux model (***Monod et al., 1965a; Koshland et al., 1966a; Marzen et al., 2013; Einav et al., 2016; Phillips, 2020***) in combination with rigorous statistical approaches to identify the relevant rate equations and estimate the kinetic constants using a manually curated dataset of thousands of data points from dozens of publications. We solved a difficult problem of parameter estimation of a large mechanistic model by fully constraining our model with 145 parameters estimated independently based on proteomics, metabolomics, and *in vitro* kinetics data from dozens of datasets. Our model robustly performed the key tasks of glycolytic ATP homeostasis and accurately predicted absolute concentrations of glycolytic intermediates and isotope tracing patterns in live cells. Analysis of our model suggests that allosteric regulation of HK and PFK, an excess of lower glycolysis enzymes, and a large labile pool of phosphate are required to maintain high ATP levels and to prevent uncontrolled accumulation of phosphorylated intermediates of upper glycolysis. Allostery achieves the latter tasks by preventing an imbalance between ATP-producing and ATP-consuming enzymes of glycolysis that, in the absence of allosteric regulation, leads to the disappearance of inorganic phosphate and phosphorylated glycolytic intermediates downstream of the GAPDH reaction and accumulation of phosphorylated glycolytic intermediates upstream of the GAPDH reaction. At the same time, mass action performed other tasks of glycolytic ATP homeostasis previously attributed to allostery, such as controlling the rate of ATP production to match ATP supply and demand, and maintaining ATP, ADP, and inorganic phosphate levels such that ATP hydrolysis reaction generates energy.

## RESULTS

### Tasks of glycolytic ATP homeostasis

Glycolytic ATP homeostasis performs multiple tasks that likely require distinct regulation. In this section, we highlight three tasks of glycolytic ATP homeostasis that we have investigated (***Figure 1A***):

Task #1: matching the rates of ATP production and consumption. Any mismatch between the ATP consumption rate by intracellular processes and glycolytic ATP production rate must only be short-lived due to the ∼1-100 seconds turnover time of intracellular ATP (***Milo and Phillips, 2015***).

Task #2: ensuring that ATP hydrolysis generates energy. The energy of ATP hydrolysis is set by the extent to which ATP hydrolysis reaction is maintained far from equilibrium, which, in turn, is determined by the relative levels of ATP, ADP, and inorganic phosphate. Inside the cell, ATP hydrolysis reaction is maintained 10^9^-10^11^-fold away from equilibrium or 20-25 k_B_T of energy (***Milo and Phillips, 2015***).

Task #3: maintaining most of the adenine nucleotide pool in the form of ATP. Maintenance of the adenine nucleotide pool in the form of ATP under different ATP consumption rates is a well-established task of ATP homeostasis that ensures that ATP levels are high and stable (***Allen et al., 1997; Balaban et al., 1986***). One function of maintaining high and stable ATP levels could be to ensure that ATP-consuming enzymes are not kinetically limited for ATP. For example, the K_M_ values for ATP of kinases (***Knight and Shokat, 2005***), myosin (***Hackney and Clark, 1985***), Na^+^/K^+^-ATPase (***Soltoff and Mandel, 1984***) are in the 1-400 µM range, indicating that these enzymes are mostly saturated with ATP given the 1-10 mM range of intracellular ATP levels. We note that we deliberately did not refer to Task #3 by the frequently used term “energy charge of the adenylate pool” (***Atkinson, 1968***) to highlight that Task #3 is unrelated to the energy of ATP hydrolysis and is distinct from Task #2 as the ATP hydrolysis reaction can generate the same amount of energy at a wide range of ATP/ADP ratios due to the offsetting effect of inorganic phosphate.

In addition, glycolysis performs the tasks above at a wide range of ATP demands, responds quickly to changes in ATP demand, maintains glycolytic intermediate concentrations within a physiological range, and is robust to the physiological variability of enzyme levels, cofactor pool sizes, extracellular glucose, and lactate concentrations, as well as kinetic constants that can change with temperature or pH. Such robustness is an important property of the homeostasis (***Barkai and Leibler, 1997***).

### Kinetic model of glycolytic ATP homeostasis

To investigate the regulation of ATP homeostasis, we have developed a comprehensive mathematical model of mammalian glycolysis. In this section, we provide an overview of the model development while most of the technical details are described in ***Materials and Methods*** as well as comprehensive ***Appendix 1*** that contain all the enzyme rate derivations. Our model includes key allosteric regulators based on a thorough literature search. Most human glycolytic enzymes are encoded by several homologous genes that produce enzyme isoforms with tissue-specific expression and distinct kinetic and allosteric properties. To facilitate experimental testing, we focused on glycolytic enzyme isoforms that are most abundant in proliferating cell lines (i.e., HK1, PFKP, and PKM2 isoforms of HK, PFK, and PK) based on proteomics data (***Geiger et al., 2012a***) (see ***Supplementary File 1*** for enzyme concentrations). We sought to develop a core model of mammalian glycolysis that supports ATP homeostasis in the absence of input from other pathways, and hence we only included allosteric regulators that are products or substrates of glycolytic enzymes. Our model converts a qualitative schematic of allosteric regulation (***Figure 1B***) into a precise mathematical language. As input, the model uses *i*) the extracellular concentrations of glucose and lactate, *ii*) cellular concentrations of cofactor pools and glycolytic enzymes, and *iii*) the rate of cellular ATP consumption (***Figure 1C***). We highlight these inputs separately from the model as these inputs are not controlled by glycolysis but depend on the cell type or experimental conditions and thus cannot be predicted by a glycolysis model. The model takes these inputs and uses enzyme rate equations assembled into a system of coupled ODEs to calculate the concentration of every glycolytic intermediate and the rate of every reaction. ***Materials and Methods*** contain a detailed description of the model, including ODE equations, stoichiometric matrix, ODE solvers, conserved moieties, simulation setting, etc. Our model can calculate both steady-state behavior and dynamical responses to perturbations. In effect, our model uses *in vitro* enzyme kinetics to predict any measurable property of glycolysis and many properties that cannot currently be measured.

In total, our model contains 172 parameters, including kinetic and thermodynamic constants and estimates of enzyme and cofactor pool concentrations, yet all these parameters are tightly constrained by experimental data. Kinetic constants were estimated from *in vitro* enzyme kinetics data (***Supplementary File 2***) as described briefly below and in full detail in ***Appendix 1***. Thermodynamic constants are taken from the eQuilibrator database (***Beber et al., 2022***). Enzyme and cofactor pool concentrations are estimated from proteomics and metabolomics data, respectively, as described in ***Materials and Methods*** and reported in ***Supplementary File 1***. We numerically simulate the model using DifferentialEquations.jl library (***Rackauckas and Nie, 2017***) written in the Julia Programming Language (***Bezanson et al., 2015***). Our code is heavily optimized so that it takes ∼10 milliseconds to calculate the results of the model under given conditions using a single core of a modern computer processor. Optimized code allows us to run the model under millions of different conditions to systematically explore the regulation of glycolytic ATP homeostasis.

The defining feature of our model is the use of newly derived kinetic rate equations to describe the activity of four allosterically regulated enzymes (i.e., HK1, PFKP, GAPDH, PKM2) using the Monod-Wyman-Changeux (MWC) model. MWC is a powerful model for describing the activity of allosterically regulated enzymes (***Monod et al., 1965a; Koshland et al., 1966a; Marzen et al., 2013; Einav et al., 2016; Phillips, 2020***), which assumes that allosteric enzymes exist in two or more conformations with different kinetic properties. Both the binding of substrates and allosteric regulators can modify the kinetic properties of an MWC enzyme by stabilizing one conformation over another. We use statistical learning approaches–regularization and cross-validation–to identify the simplest MWC kinetic rate equation that adequately describes the available *in vitro* kinetic data, allowing us to avoid overfitting and parameter identifiability issues common for fitting complex equations to finite data (***Transtrum et al., 2015***). The MWC equation describing the rate of PFKP in the presence of substrates and regulators is shown in ***Figure 1D***. We described non-allosterically regulated glycolytic enzymes (black in ***Figure 1B***) using standard kinetic rate equations derived from quasi-steady-state or rapid equilibrium approximations. For the eight enzymes and transporters with one substrate and one product and for aldolase (ALDO) with two products, we used reversible Michaelis-Menten equations (see equation describing the rate of glucose-6-phosphate isomerase (GPI) in ***Figure 1D***) and sequential release equation for ALDO, respectively, and estimated their kinetic constants by averaging the values from at least three publications per constant to verify their consistency and accuracy. For the two remaining enzymes with more than one substrate or product–phosphoglycerate kinase (PGK) and lactate dehydrogenase (LDH)–we fitted their more complex rate equations to manually curated *in vitro* kinetics datasets containing 350-700 data points per enzyme (***Supplementary File 2***). We chose to fit data from PGK, and LDH instead of averaging over published kinetic constants because many publications describing PGK, and LDH activity used different rate equations, and hence the published kinetic constants are not directly comparable. A comprehensive description of the derivations and fitting of all the enzyme rate equations is reported in ***Appendix 1*** and *in vitro* kinetic data is compiled in ***Supplementary File 2***, and we hope this compendium of information will serve as a great resource for future investigations of these enzymes.

Our model uses several assumptions that must be considered when interpreting its predictions. First, the model assumes that the activity of enzymes in living cells is accurately described using *in vitro* activity of purified enzymes. Second, the model assumes that enzymes and metabolites are well-mixed in the cytosol of the cell. Third, the model only describes the effect of regulators that are included in the model. Fourth, the model assumes that other pathways (e.g., pentose phosphate pathway, mitochondrial pyruvate consumption) do not affect the concentration of glycolytic intermediates or glycolytic reaction rates. Deviations between the model and experimental observations could be due to any number of these assumptions not being valid under the conditions of a particular experiment. On the other hand, if the model accurately predicts experimental data, it suggests that these assumptions are valid under given experimental conditions.

### Glycolysis model robustly performs the key tasks of glycolytic ATP homeostasis

After constructing the model, we first performed simulations to determine whether the glycolysis model can perform the key tasks of ATP homeostasis described above (***Figure 1A***). To model ATP consumption by intracellular processes, we added an ATPase enzyme to the model. Glycolysis cannot proceed faster than the maximal rate of its slowest enzyme, so we report all ATPase rates in percent of the maximal activity of HK1–the slowest enzyme of glycolysis based on the product of intracellular enzyme concentration and *V*_*max*_. First, we found that the glycolysis model produces as much ATP as is consumed and rapidly adjusts to stepwise increase or decrease of ATP consumption, as is necessary under physiological conditions (***Figure 2A***). Second, we showed that > 90% of the adenine nucleotide pool was maintained as ATP and ATP concentrations varied by < 10% when we introduced physiologically relevant 2-fold stepwise changes in ATP consumption rate (***Figure 2B***). Thus, as has been observed *in vivo* (***Allen et al., 1997; Balaban et al., 1986***), our model maintains nearly constant ATP levels in response to large changes in ATP turnover. Finally, we showed that the glycolysis model maintains ATP, ADP and inorganic phosphate concentrations such that the ATP hydrolysis reaction is ∼10^9^-fold away from equilibrium, equivalent to ∼20-25 k_B_T (***Figure 2C***), which is similar to what has been measured in cells (***Milo and Phillips, 2015***).

**Figure 2.**
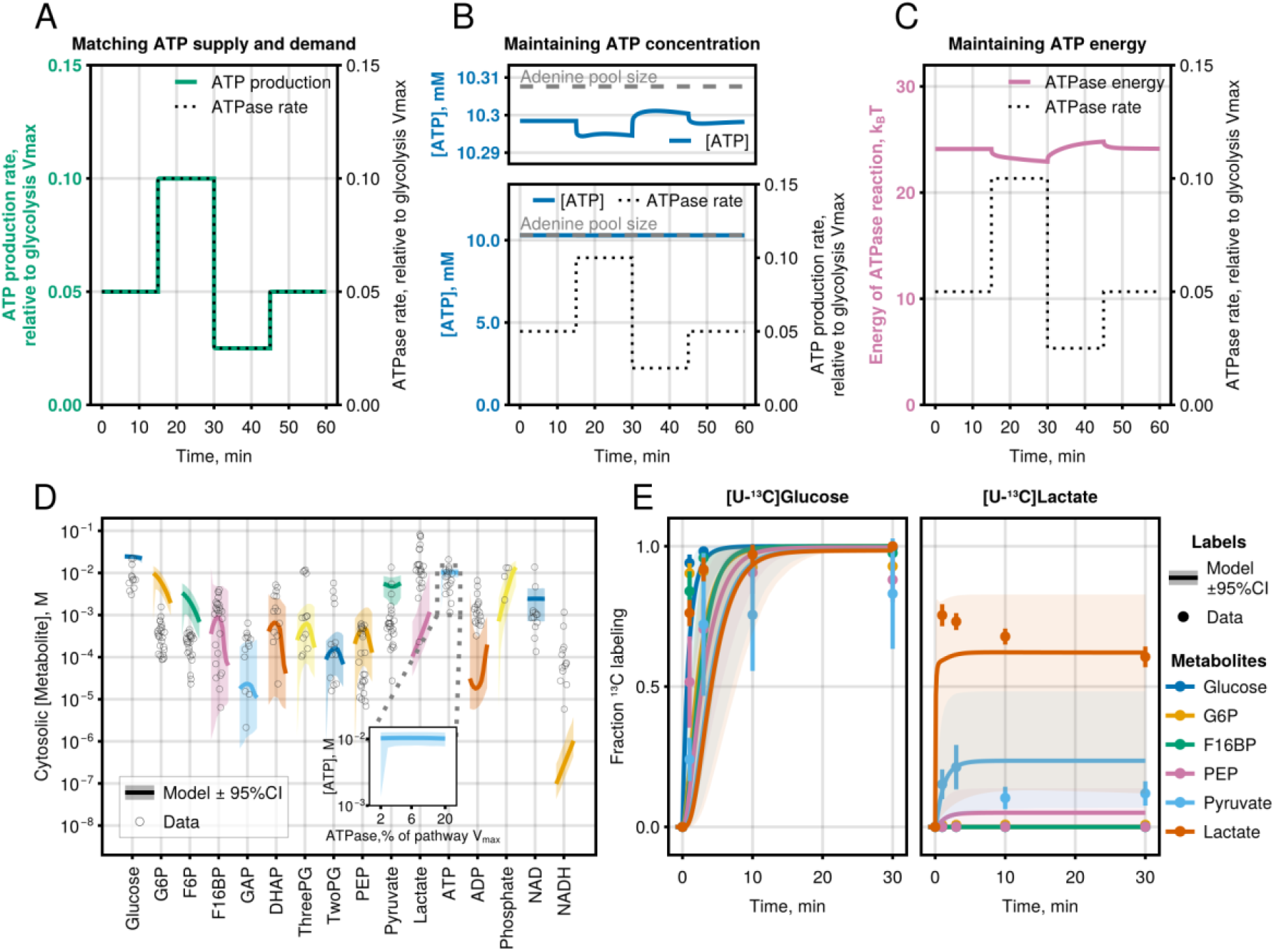
Glycolysis model recapitulates glycolysis activity observed in live cells. **(A-C)** Model simulations showing changes in (A) ATP production rate, (B) ATP concentration (total adenine pool size is labeled with dashed grey line), and (C) energy released during ATP hydrolysis in response to a 2-fold stepwise increase or decrease of ATPase rate. Starting ATPase rate = 5% of glycolysis *V*_*max*_ was used for (A-C). ATPase energy is calculated as a natural logarithm of the disequilibrium ratio for the ATPase reaction (i.e., mass-action ratio divided by the equilibrium constant) for the ATPase reaction. **(D)** Comparison of model predictions with measured glycolytic intermediate concentrations from multiple cell lines and experimental conditions. Black empty circles indicate the experimentally determined metabolite concentrations. Colored lines with ribbon indicate the median and 95% CI of model predictions. The shift from left to right for each metabolite level prediction corresponds to increasing rates of ATP consumption from 2% to 20% of glycolysis *V*_*max*_ (see inset for ATP) to highlight the effect of ATP consumption on metabolite concentrations. Note that data points are displayed with pseudorandom jitter and do not have shifts corresponding to ATP consumption rate. Model predictions represent the concentration of bound + free metabolites in the whole cell analogous to whole cell metabolite measurements, while in all other figures, model output is the free concentration of metabolites in the cytosol. **(E)** Comparison of model prediction with [U-^13^C_6_]Glucose and [U-^13^C_3_]Lactate tracing data averaged for C2C12 and HeLa cells. Points are data (error bars are SD), and lines are model predictions (ribbons are 95% CI). ATPase rate = 15% of glycolysis Vmax was chosen for model predictions. Model predictions in (D) and (E) are from simulations where bootstrapped model parameter combinations could match ATP supply and demand, which were > 97% and >95% of simulations for (D) and (E), respectively. See also ***Figure 2–figure supplement 1***. Julia code to reproduce this figure is available at https://github.com/DenisTitovLab/Glycolysis.jl.

We next tested whether the ability of the model to perform the relevant tasks of ATP homeostasis is robust to changes in glycolytic enzyme levels that are variable from cell to cell and to uncertainty in our estimates of biochemical constants, cofactor pool concentrations, and initial concentrations of intermediates. To demonstrate robustness, we reran our model 10,000 times using random values of enzyme levels spanning a 9-fold range around the model estimates based on proteomics data (***Figure 2–figure supplement 1A-C***) as well as random values of model parameters (***Figure 2–figure supplement 1D-F***) and initial conditions (***Figure 2–figure supplement 1G-K***) based on their respective estimated mean and standard error (see ***Supplementary File 1*** for model estimates on enzyme levels and initial conditions and ***Appendix 1*** for kinetic parameter mean and standard error). Our simulations showed that 95% confidence intervals of model prediction are largely independent of glycolytic enzyme levels, model parameters, and initial conditions, as would be expected for the model that captures the robust endogenous regulatory circuit of glycolytic ATP homeostasis.

### Glycolysis model accurately predicts metabolite levels and fluxes in living cells

Given that our model could reproduce the wealth of measurements of each glycolytic enzyme *in vitro* and perform the key tasks of ATP homeostasis, we set out to determine whether the model could predict the results of experiments in live cells.

We first compared the model predictions of absolute intracellular metabolite concentrations with corresponding measurements. To test if our model can recapitulate the vast range of glycolytic activities seen in prior experiments, we not only compiled data from nine publications analyzing whole-cell absolute intracellular glycolytic intermediate concentrations using LC-MS, but also supplemented these data with our own measurements in four cell lines (see ***Supplementary File 1***). Our model predicts free metabolite concentration in the cytosol while LC-MS measures the sum of free and enzyme-bound metabolite concentrations in whole cells. To directly compare model predictions to data from bulk cell measurements, we adjusted the model predictions to report the sum of enzyme-bound and free metabolite concentration and adjusted the whole-cell measurements to report cytosolic metabolite concentrations (see ***Materials and Methods*** for additional details of the adjustments). In addition, we reported model predictions at a range of cellular ATP demands as the ATP demand of the cell is not known for the LC-MS measurements. We observed that 95% confidence intervals of model predictions of most of the 18 glycolytic intermediate concentrations overlapped with experimental measurement (***Figure 2D***). The latter result is notable given that our model contains no direct information about intracellular metabolite levels and is free to predict concentrations from 0 to +∞ and even below zero if the model is implemented incorrectly. The confidence intervals of model predictions for BPG and NADH do not overlap with experimental values. The prediction of lower levels of NADH might be attributed to the fact that most of NADH is in the mitochondria (***Williamson et al., 1967***)–a compartment not included in our model. The prediction of lower levels of BPG might be attributed to the fact that most of the BPG in our model predictions is bound to GAPDH, but whole-cell lysis techniques cannot measure this BPG pool as it forms a covalent complex with GAPDH (***Furfine and Velick, 1965***).

Next, we compared [U-^13^C_6_]Glucose and [U-^13^C_3_]Lactate labeling kinetics predicted by the model to cellular measurements. For these measurements, we exchanged normal media for media containing [U-^13^C_6_]Glucose or [U-^13^C_3_]Lactate and then lysed cells at different time intervals to estimate the rate and fraction at which ^13^C from [U-^13^C_6_]Glucose or [U-^13^C_3_]Lactate is incorporated into glycolytic intermediates. We observed that the 95% confidence intervals of glycolysis model prediction overlapped with the measurements of the ^13^C labeling fraction of intermediates after switching to [U-^13^C_6_]Glucose or [U-^13^C_3_]Lactate-containing media (***Figure 2E***). It is particularly noteworthy that the model can recapitulate labeling from [U-^13^C_3_]Lactate where only intracellular lactate and pyruvate are labeled, as this indicates that our model can accurately capture the extent of *reversibility* of glycolysis reactions.

Our model can recapitulate measurements of glycolytic intermediate levels and fluxes observed in living cells. Additional purposefully designed experiments will be able to pinpoint if quantitative differences between model predictions and experimental estimates that are observed are due to inaccuracies of the model, systematic experimental measurement error, or due to the fact that model predictions and experiments are reporting different values (e.g., whole cell vs cytosolic levels of metabolites).

### Allosteric regulation is required to robustly maintain high ATP levels while mass action performs other tasks of glycolytic ATP homeostasis

Having established that the model recapitulates the key tasks of ATP homeostasis and accurately predicts measurements in live cells, we next used the model to investigate the function of allosteric regulation and mass action in maintaining ATP homeostasis.

We first removed all allosteric regulators to see if the resulting simple pathway remained functional. Specifically, we made the HK1, PFKP, GAPDH, and PKM2 enzymes behave as Michaelis-Menten-like enzymes with kinetic parameters corresponding to their active MWC conformation. We also removed inhibition of the HK catalytic site by G6P, which proceeds through standard non-allosteric competitive inhibition. Surprisingly, the model without any regulation could match ATP supply and demand and maintain the high energy of ATP hydrolysis with only minor quantitative differences compared to the model with allostery (***Figure 3A, C, D, F***). However, we observed a complete breakdown of high ATP level maintenance without allosteric regulation, where ATP levels were >100-fold lower, and a small 2-fold increase and decrease in ATPase rate led to an almost 10-fold change in ATP concentration compared to <10% change in ATP concentration for the complete model (***Figure 3B, E***). Systematic investigation of steady-state model behavior at different ATPase rates showed that the model without regulation is not capable of maintaining most of the adenine nucleotide pool in the form of ATP at any ATPase rate, which is contrary to the constant ATP concentrations observed both *in vivo* (***Allen et al., 1997; Balaban et al., 1986***) and in the complete model (***Figure 3E***, vertical drop-off indicates where the ATP production by the model cannot match ATPase rate anymore). Finally, we note that the full model fails to maintain most of the adenine pool in the form of ATP at high ATPase rate–an expected consequence of ATPase rate being higher than the activity of glycolytic enzymes–and low ATPase rate–an unexpected observation that we will explore below as it connects to the function of allosteric regulation of glycolysis (***Figure 3E***, vertical drop-offs at high and low ATPase rates).

**Figure 3.**
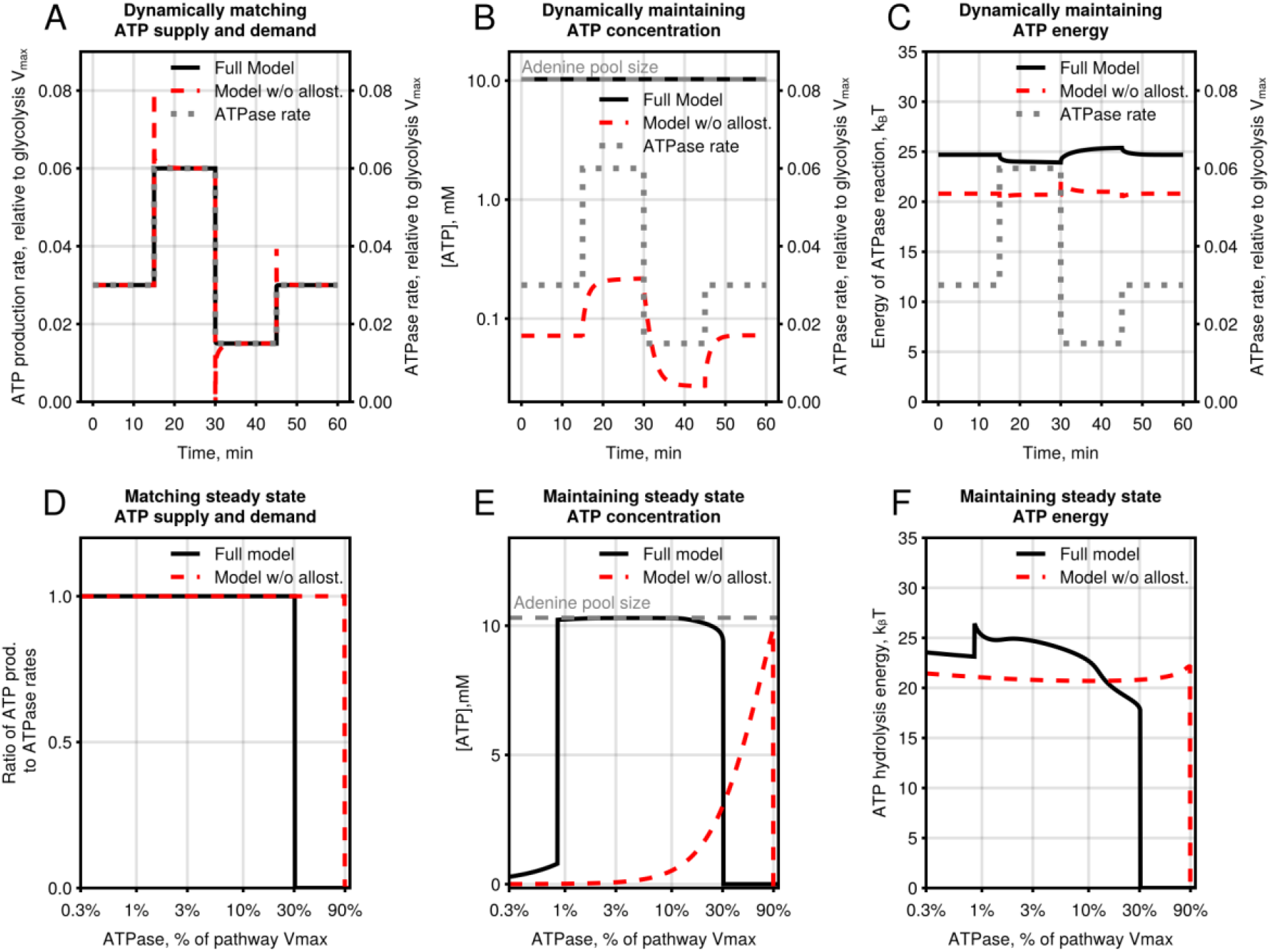
Allosteric regulation is required for robust maintenance of high and stable ATP levels. **(A-C)** Model simulations showing the effect of removal of allosteric regulation of HK1, PFKP, GAPDH, and PKM2 on dynamic changes in (A) ATP production rate, (B) ATP concentration, and (C) energy released during ATP hydrolysis [Left axes] in response to a 2-fold stepwise increase or decrease of ATPase rate [Right axes]. Starting ATPase rate = 3% of glycolysis *V*_*max*_ was used for (A-C). Energy is calculated as a natural logarithm of the disequilibrium ratio (i.e., mass-action ratio divided by the equilibrium constant) for the ATPase reaction. **(D-F)** Model simulations showing the effect of removal of allosteric regulation of HK1, PFKP, GAPDH, and PKM2 on steady-state (D) ATP production rate, (E) ATP concentration, and (F) energy released during ATP hydrolysis at a 100-fold range of ATPase rates. Dashed grey lines in B and E indicate the total adenine pool size (i.e., ATP + ADP + AMP). See also ***Figure 3–figure supplement 1***. Julia code to reproduce this figure is available at https://github.com/DenisTitovLab/Glycolysis.jl.

### Comparison of our model to published glycolysis models

Several kinetics models of glycolysis have been reported over the past several decades (***Sel’kov, 1975; Rapoport et al., 1976; Termonia and Ross, 1981; Joshi and Palsson, 1990; Teusink et al., 1996; Mulquiney and Kuchel, 1999; Teusink et al., 2000; Hynne et al., 2001; Chandra et al., 2011; Eunen et al., 2012; Smallbone et al., 2013; van Heerden et al., 2014; Shestov et al., 2014; Mulukutla et al., 2014***), yet none of these reports investigated whether those models can recapitulate the three tasks of ATP homeostasis that we identified. Most of the published models (***Sel’kov, 1975; Rapoport et al., 1976; Termonia and Ross, 1981; Joshi and Palsson, 1990; Teusink et al., 1996, 2000; Hynne et al., 2001; Chandra et al., 2011; Eunen et al., 2012; Smallbone et al., 2013; Mulukutla et al., 2014***) either did not include ATP, ADP, or Pi or kept them constant, making it difficult to investigate ATP homeostasis using those models. We performed head-to-head comparisons of our model with three mechanistic models of glycolysis that we will refer to as the Mulquiney model (***Mulquiney and Kuchel, 1999***), the van Heerden model (***van Heerden et al., 2014***), and the Shestov model (***Shestov et al., 2014***). To the best of our knowledge, the latter are the only three mechanistic models of glycolysis that use *in vitro* enzyme kinetics and include variable ATP, ADP, and Pi. Mulquiney and Shestov models simulate mammalian glycolysis and van Heerden model–*S. cerevisiae* glycolysis. All three models incorporate allosteric regulation of HK, PFK, and PK enzymes, but each model only contains a subset of allosteric regulators included in our model, and none of the models were previously used to investigate the function of allosteric regulation of glycolysis. We repeated the simulations in ***Figure 3D-F*** using the kinetic rate equations for glycolytic enzymes from the Mulquiney, Shestov, and van Heerden models (***Figure 3–figure supplement 1***). All three models could match ATP supply and demand and maintain the high energy of ATP hydrolysis (***Figure 3–figure supplement 1A, D, G, C, F, I***). Note that the Mulquiney, van Heerden, and our model could match ATP supply and demand at similar ATPase rates up to about 10-30% of the respective pathway *V*_*max*_, while the Shestov model could only support much lower ATPase rates up to 0.003% of its *V*_*max*_. The Mulquiney and Shestov models could maintain the majority of the adenine pool in the form of ATP at a 2.8- and 2.2-fold range of ATPase values compared to a 38-fold range for our model, while van Heerden model could not maintain high ATP under any ATPase rates (***Figure 3–figure supplement 1B, E, H***). The ability of our model to maintain high ATP levels at a wider range of ATPase rates compared to published models is likely the result of incorporating more allosteric regulators and fitting MWC rate equation to kinetic data from multiple publications. Importantly, as for our model, the ability of the Mulquiney and Shestov models to maintain high ATP levels depended on the presence of allosteric regulation (***Figure 3– figure supplement 1B, E, H***). Comparison to previously published models further confirms our observations that the maintenance of high ATP levels requires allosteric regulation. In contrast, mass action is sufficient for the ability of glycolysis to match ATP supply and demand and maintain the high energy of ATP hydrolysis.

### Redundant allosteric regulators of HK1 and PFKP maintain high and stable ATP levels

To identify the role of specific allosteric regulators in maintaining high and stable ATP levels, we computationally dissected the pathway by removing the allosteric regulation of enzymes one by one (***Figure 4***). Specifically, we removed a particular allosteric regulator from the relevant kinetic rate equations by setting its binding constant for both active and inactive MWC conformation to ∞ and setting the constant *L* that determines the ratio of inactive to active MWC conformations to zero. We found that the allosteric regulation of HK1 and PFKP is responsible for maintaining high and stable ATP levels, whereas removing the allosteric regulation of GAPDH and PKM2 had no discernable effect (***Figure 4A-D***). Digging further, we removed each of the allosteric activators and inhibitors of HK1 and PFKP and found that these regulators work together to ensure the robust maintenance of ATP levels, since removing any single regulator only led to a partial loss of ATP maintenance capacity. In general, removing inhibitors–G6P for HK1 and ATP for PFKP–led to worse ATP maintenance at low ATPase rates (***Figure 4E-H***), while removing activators–phosphate for both HK1 and PFKP and ADP for PFKP–led to poorer ATP maintenance at high ATPase rates (***Figure 4K-N***).

**Figure 4.**
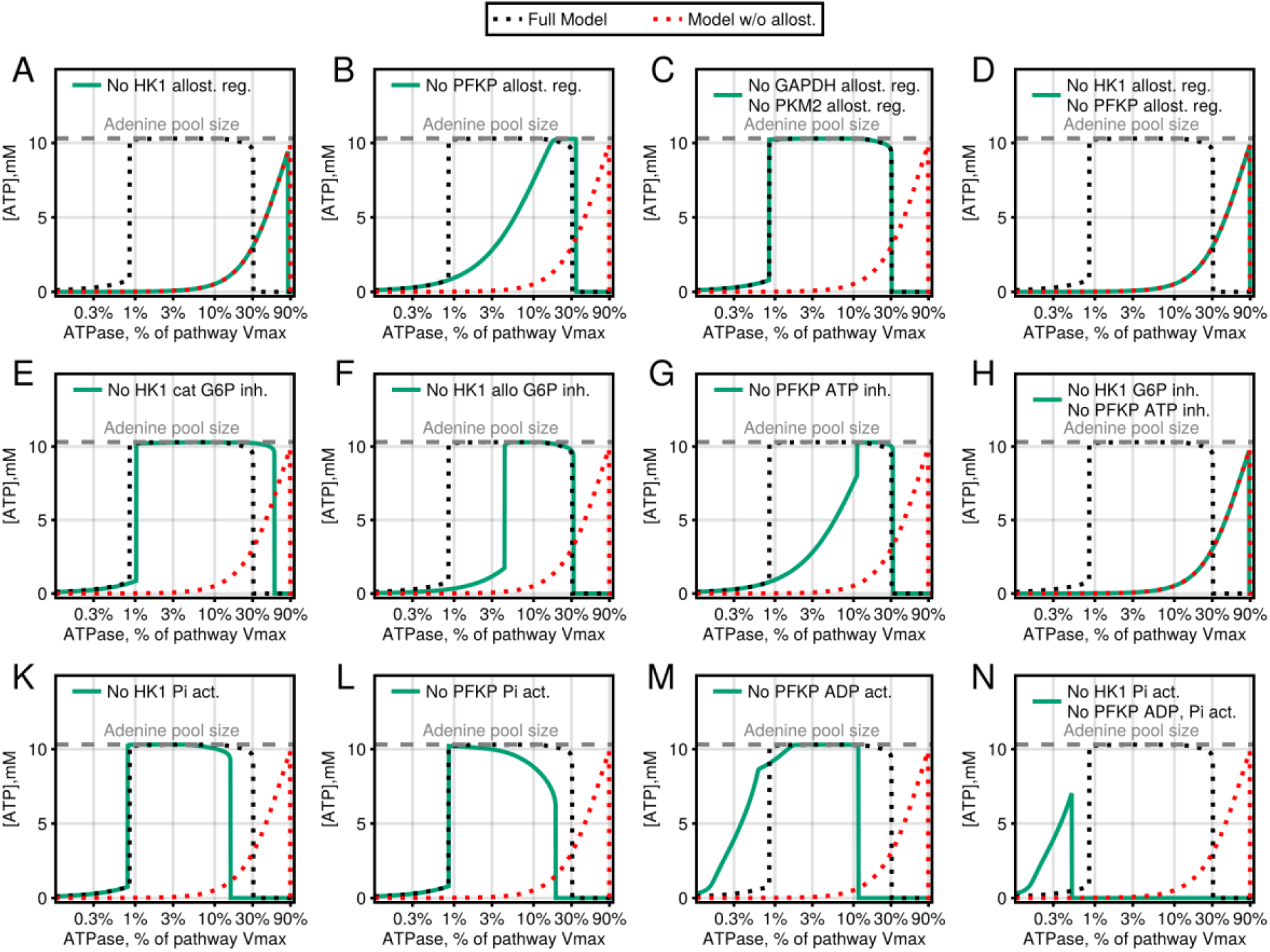
Redundant allosteric regulators of HK1 and PFKP are required to maintain high and stable ATP levels. **(A-N)** Steady-state ATP concentrations at a range of ATPase rates of the glycolysis model with and without (A-D) allosteric regulation of enzymes, (E-H) allosteric inhibitors, and (K-N) allosteric activators. Dashed grey line indicates the total adenine pool size (i.e., ATP + ADP + AMP). See also ***Figure 4–figure supplement 1***. Julia code to reproduce this figure is available at https://github.com/DenisTitovLab/Glycolysis.jl.

### Global sensitivity analysis of the glycolysis model

We next performed a global sensitivity analysis of our model to systematically explore the role of all model parameters in matching ATP supply and demand, maintaining high ATP levels, and maintaining high energy of ATP hydrolysis. The goal of global sensitivity analysis is to estimate the contribution of each model parameter and parameter interactions to the variance of a specific model output (***Saltelli, 2008***). We used the average ratio of ATP production to ATP consumption, average ratio of ATP concentration to adenine pool size, and average energy of ATP hydrolysis at a log range of ATPase values as proxies for the model’s ability to match ATP supply and demand, maintain high ATP levels and maintain high energy of ATP hydrolysis, respectively. We first estimated the variance of the proxies by randomly varying each parameter of the model independently from a log-uniform distribution spanning a 9-fold range from 3 times lower to 3 times higher than its value in the model. The coefficient of variation (CV) for all three proxies was in the narrow range of 0.15-0.34 in response to random changes in parameter values spanning 9-fold, indicating that these are robust properties of the model (***Figure 4–figure supplement 1A-C***). The coefficient of variation for maintaining high ATP levels was about 2-fold higher than for maintaining high energy of ATP hydrolysis or for matching ATP supply and demand, suggesting that the former is more sensitive to changes in specific enzyme kinetic parameters as would be expected for a task that depends on allosteric regulation and not just mass action. Two metrics for each model parameter are typically reported for variance-based global sensitivity analysis. The first-order effect sensitivity index *S*_*1*_ reports the fraction of variance of the model output that will be removed if the corresponding parameter is fixed. The total-order sensitivity index *S*_*T*_ reports the variance that will be left if values of all but the corresponding parameter are fixed. Larger values of *S*_*1*_ and *S*_*T*_ indicate that a given parameter is important. Global sensitivity analysis showed that kinetic parameters for HK1, PFKP, and GAPDH had the highest *S*_*1*_ and *S*_*T*_ for all three model outputs. We subdivided kinetic parameters for HK1 and PFK into allosteric and non-allosteric parameters and showed that non-allosteric parameters had highest *S*_*1*_ and *S*_*T*_ for matching ATP supply in demand (***Figure 4–figure supplement 1D***), as would be expected given that HK1 and PFK are the slowest enzymes in the pathway, while allosteric parameters had highest *S*_*1*_ and *S*_*T*_ for maintain high and stable ATP levels (***Figure 4–figure supplement 1E***). Thus, global sensitivity analysis confirmed the importance of allosteric regulation in maintaining high and stable ATP levels, as shown in ***Figure 3***.

### Allosteric regulation maintains high ATP levels and prevents uncontrolled accumulation of phosphorylated intermediated by dynamically restricting the maximal rate of HK and PFK to half of the ATPase rate

We next investigated the mechanism that allows allosteric regulators of HK1 and PFKP to maintain high and stable ATP levels such that most of the adenine pool is in the form of ATP. Here, we first provide an intuitive explanation of the mechanism and then show simulations supporting it. ATP levels are bounded by the size of the adenine pool from the top and the ability of glycolysis to convert most of the ADP into ATP from the bottom. The latter leads to near-constant ATP levels under any conditions where glycolysis can convert most of the ADP into ATP. Glycolysis has an unusual organization where upper glycolysis enzymes–from HK1 to ALDO–consume ATP, while lower glycolysis enzymes–from GAPDH to LDH–consume inorganic phosphate and produce ATP. Such an organization can lead to the trapping of inorganic phosphate such that GAPDH and PGK enzymes of lower glycolysis convert inorganic phosphate to ATP that is then used by HK1 and PFKP enzymes of upper glycolysis to produce phosphorylated glycolytic intermediates upstream of GAPDH–G6P, F6P, F16BP, DHAP, and GAP (***Figure 5A***). The net reaction of phosphate trapping is:

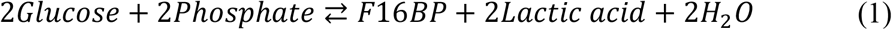

This net reaction is extremely thermodynamically favorable with an estimated (***Beber et al., 2022***) *K*_eq_ ≈ 3 ·10^30^ and Δ_*r*_*G*^′°^ ≈ −170 *kJ*/*mol*, resulting in the sequestration of inorganic phosphate in phosphorylated intermediates of upper glycolysis. In the presence of exogenous phosphate uptake (e.g., uptake from the extracellular media), the consequence of the phosphate trap is the uncontrolled accumulation of phosphorylated intermediates of upper glycolysis to concentrations that will explode cells. In the absence of exogenous phosphate uptake, the consequence of the phosphate trap is low levels of free inorganic phosphate that inhibit the GAPDH reaction and lead to a lower concentration of substrates of ATP-producing enzymes of lower glycolysis–BPG for PGK and PEP for PKM2–requiring higher levels of ADP to drive PGK and PKM2 forward through mass action. Thus, the presence of a thermodynamically favorable phosphate trap makes it either impossible for glycolysis to maintain a high ATP/ADP ratio or explodes the cells due to uncontrolled accumulation of upper glycolysis intermediates. We propose that the function of allosteric regulation of HK1 and PFKP is to restrict their rates to half of the ATPase rate, making the phosphate trap impossible due to the lack of excess HK1 and PFKP capacity necessary to catalyze the trapping reactions (***Figure 5A***).

**Figure 5.**
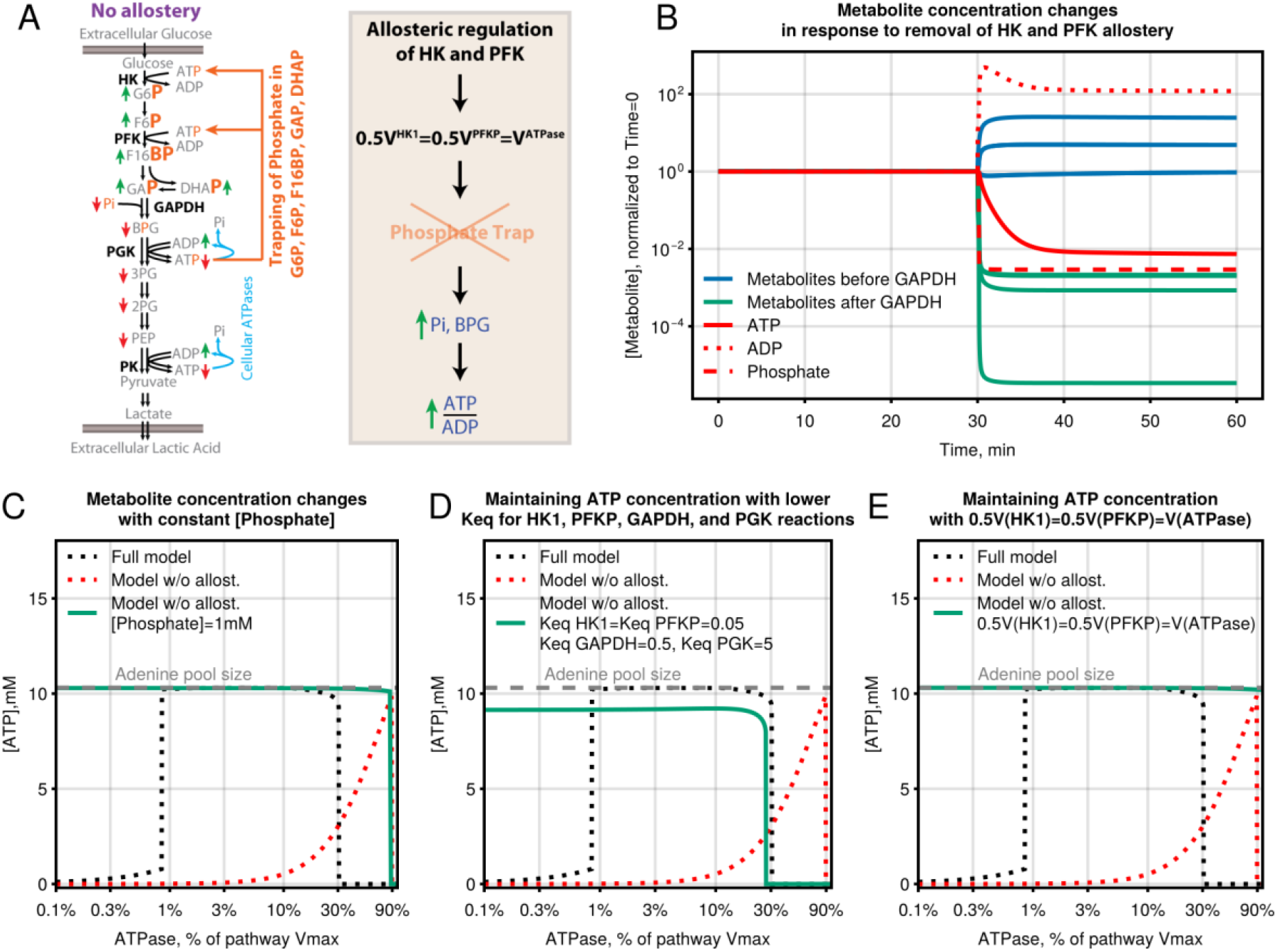
Allosteric regulation maintains ATP levels by preventing a phosphate trap due to an imbalance between lower and upper glycolysis. **A)** Schematic of the mechanism of ATP maintenance by allosteric regulation of HK1 and PFKP. **B)** Phosphorylated metabolite level changes upon an instantaneous removal of allostery at 30 min. ATPase rate = 3% of glycolysis Vmax was used for model predictions. **C-E)** Steady-state ATP concentrations at a range of ATPase rate of the glycolysis model with (C) constant [Phosphate] = 1mM, (D) *K*_*eq*_^*HK1*^=0.05 (model value 2,700), *K*_*eq*_^*PFKP*^=0.05 (model value 760), *K*_*eq*_^*GAPDH*^=0.5 (model value 16), and *K*_*eq*_^*PGK*^=5 (model value 2000), (E) Rate equations for HK1 and PFKP substituted for one-half of ATPase rate equations. See also ***Figure 5–figure supplement 1***. Julia code to reproduce this figure is available at https://github.com/DenisTitovLab/Glycolysis.jl.

We performed simulations to support the role of allostery in preventing the phosphate trap mechanism shown in ***Figure 5A***. Removal of allosteric regulation of HK1 and PFKP leads to upregulation of phosphorylated metabolites upstream of the GAPDH reaction except for phosphate and downregulation of phosphorylated metabolites downstream of the GAPDH reaction, suggesting that low phosphate blocks the GAPDH step in the absence of allosteric control of HK1 and PFKP (***Figure 5B***). We next showed that keeping constant levels of inorganic phosphate at 1 mM abrogated the requirement for allostery to maintain ATP levels, showing that allostery works by maintaining high phosphate levels (***Figure 5C***). Similarly, a large decrease in equilibrium constants of HK1, PFKP, GAPDH, and PGK that makes the phosphate trap less thermodynamically favorable also removed the requirement for allostery (***Figure 5D***). Finally, substituting HK1 and PFKP rate equations for one-half of the ATPase rate abolished the requirement for allostery, suggesting that the latter works by tying HK1 and PFKP rates to the ATPase rate, which makes the phosphate trap impossible. Importantly, maintaining constant phosphate levels (e.g., through uptake across plasma membrane) cannot substitute for allosteric regulation as the former will lead to the accumulation of phosphorylated upper glycolysis intermediates to >100,000 M, which is well above the 55M concentration of water in water and will presumably explode cells (***Figure 5–figure supplement 1***).

We note that most of the published models of glycolysis assumed constant phosphate levels, which made it impossible to identify the role of allosteric regulation.

### Excess activity of enzymes relative to HK and PFK is required to maintain high ATP levels

We next explored whether attributes of the glycolysis pathway other than allosteric regulation of HK1 and PFKP might be required for maintaining most of the adenine pool in the form of ATP. During model construction, we noted that proteomics data shows that most mammalian glycolytic enzymes are expressed in large excess compared to HK1, PFKP, glucose transporter, and lactate transporter (***Figure 6A***). After accounting for specific activity, the maximal cellular activity of most enzymes is at 10-100-fold higher levels than is required for the maximal activity of the glycolysis pathway that is limited by HK1 (***Figure 6B***). Allosteric inhibition of HK1 and PFKP by G6P and ATP, respectfully, further increases the differential between the activity of HK1 and PFKP and the rest of the enzymes. We used our model to understand the rationale for this seemingly wasteful enzyme expression pattern, given that the production of excess glycolytic enzymes represents a significant investment of resources by the cell, with enzymes like GAPDH and PKM2 approaching 1% of the proteome. Increasing the concentrations of enzymes from TPI to LDH by 10-fold led to no changes in ATP maintenance (***Figure 6C***). Decreasing the concentrations of these enzymes by 10- and 50-fold led to a small decrease and complete collapse of ATP level maintenance, respectively, without affecting the ability of the model to produce ATP (***Figure 6C***). Low expression of HK1 and PFKP contributes to limiting the occurrence of the phosphate trap, as the latter requires the excess activity of HK1 and PFKP. We propose that the high expression of glycolytic enzymes in relation to HK1 and PFKP works together with allosteric regulation to maintain high and stable ATP levels. Our results provide a mechanistic explanation for the seemingly wasteful large excess expression of enzymes of glycolysis relative to HK and PFK.

**Figure 6.**
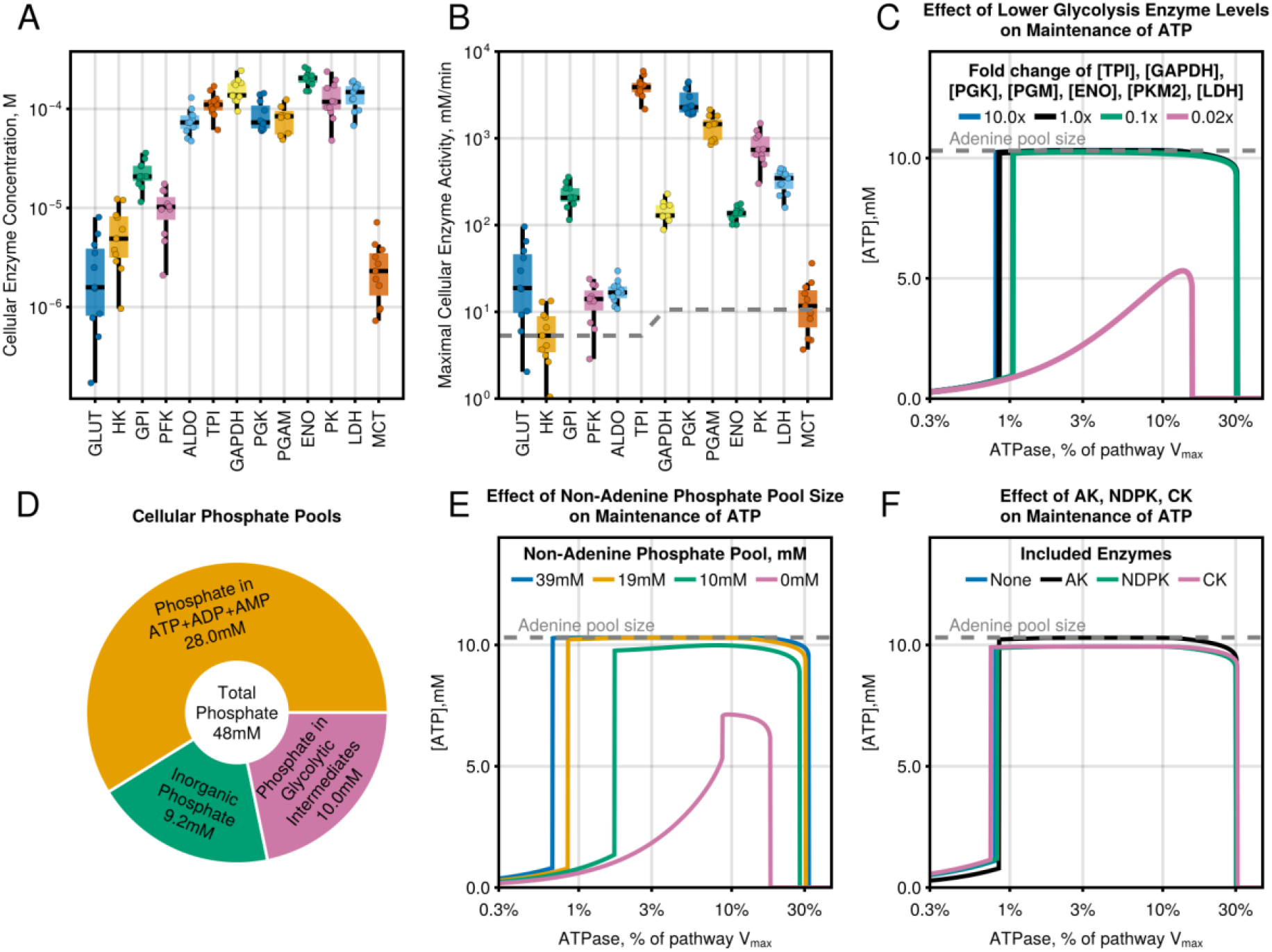
Excess activity of glycolytic enzymes relative to HK and PFK and large non-adenine phosphate pool improve the maintenance of high and stable ATP levels. **(A)** Cytosolic enzyme concentrations in the model estimated from proteomics data. **(B)** Maximal cytosolic enzyme activities in the model. Activities of all isoforms for each enzyme were summed together. The dashed line indicates the activity for each enzyme that matches the maximal activity of the slowest enzyme HK1 corrected for the stoichiometry of reactions. **(C)** Steady-state ATP concentrations at a range of ATPase rates of the glycolysis model with scaled concentrations of lower glycolysis enzymes TPI, GAPDH, PGK, PGAM, ENO, PK, and LDH. **(D)** Phosphate content of various cellular metabolite pools in proliferating mammalian cell lines based on LC-MS measurement in ***Supplementary File 1***. Metabolite levels were corrected for the number of phosphate atoms and assumed to localize to the cytosol. **(E)** Steady-state ATP concentrations at a range of ATPase rates of the glycolysis model with different levels of non-adenine phosphate pool size. **(F)** Steady-state ATP concentrations in the presence or absence of Adenylate Kinase (AK), NDP Kinase (NDPK), and Creatine Kinase (CK). The dashed grey line indicates the total adenine pool size (i.e., ATP + ADP + AMP). See also ***Figure 6–figure supplement 1***. Julia code to reproduce this figure is available at https://github.com/DenisTitovLab/Glycolysis.jl.

### Large non-adenine phosphate pool size improves the maintenance of high ATP levels

We next investigated the role of cofactor pool sizes in maintaining ATP levels. The three cofactor pools that are included in our model are the adenine nucleotide pool (i.e., ATP+ADP+AMP), NAD(H) pool (i.e., NAD^+^ + NADH), and the non-adenine phosphate pool (i.e., the sum of all phosphate atoms except for ATP, ADP, and AMP). We considered the phosphate content of the adenine and non-adenine pools separately to distinguish the effects of phosphate and adenine nucleotide pool sizes. Experimental estimates compiled in ***Supplementary File 1*** suggest that metabolically accessible inorganic phosphate in cells (i.e., phosphate not fixed in nucleic acids or lipids) is evenly split between adenine and non-adenine pools, with the latter evenly split between free inorganic phosphate and phosphorylated glycolytic intermediates (***Figure 6D***). We performed a global sensitivity analysis by varying cofactor pool sizes over a 9-fold range centered around the values of each pool size estimated from the literature and our LC-MS measurements (see ***Supplementary File 1***). Based on *S*_*1*_ and *S*_*T*_ sensitivity indexes, the non-adenine phosphate pool size had the largest effect on the model’s ability to maintain high ATP levels, while adenine pool size had a smaller effect and NAD(H) pool size had little to no effect (***Figure 6–figure supplement 1A-B***).

To confirm the effect of the non-adenine phosphate pool on maintaining ATP levels, we simulated an increase or decrease of the level of all phosphorylated metabolites except ATP, ADP, and AMP. The simulations confirmed that changes in the non-adenine phosphate pool led to a large change in the model’s ability to maintain high and stable ATP levels with the near-complete collapse of ATP maintenance in the absence of the non-adenine phosphate pool (***Figure 6E***). It is noteworthy that the experimentally estimated non-adenine phosphate pool size used in our model is close to optimal as its decrease led to the deterioration of the model’s ability to maintain high and stable ATP levels at a range of ATPase rates and decrease had a small effect. The effect of non-adenine phosphate pool size is driven by the concentration of inorganic phosphate, which, if kept constant, abrogates the effect of a changing non-adenine phosphate pool (***Figure 6–figure supplement 1C***). Cells have complex regulatory machinery to regulate intracellular phosphate content, and it is likely that this machinery–which is not part of our model–works together with allosteric regulation and relative expression of glycolytic enzymes to maintain high and stable ATP levels.

Finally, we confirmed the effect of adenine pool size on ATP level maintenance using simulations where we increased or decreased the combined level of ATP, ADP, and AMP (***Figure 6–figure supplement 1D***). As for the non-adenine phosphate pool, the size of the adenine nucleotide pool that we estimated from metabolomics data and used in our model was optimal for the ability of the model to support high and stable ATP levels. The observations that both non-adenine phosphate pool and adenine nucleotide pool sizes estimated from metabolomics data are optimal for the ability of our model to support high and stable ATP levels provide additional evidence of the capacity of our model to describe the physiologically relevant behavior of glycolysis.

### Effect of Creatine Kinase on ATP homeostasis is driven by the non-adenine phosphate pool size

We next tested the roles of Creatine Kinase (CK), Adenylate Kinase (AK), and Nucleotide Diphosphate Kinase (NDPK) in ATP homeostasis. Our model contains AK, and for this analysis, we included NDPK and CK. We first tested the ability of our model to maintain the majority of the adenine pool in the form of ATP in the presence or absence of AK, NDPK, and CK and observed no benefit of including either of these enzymes at steady state (***Figure 6F***) or in response to dynamic changes in ATPase rate. In the absence of the non-adenine phosphate pool, CK rescued the maintenance of ATP in the presence of creatine pool size comparable to the 20mM estimate for the non-adenine phosphate pool, while AK and NDPK had only a minor effect (***Figure 6–figure supplement 1E-G***). The creatine pool size in muscle cells, ∼20 mM (***Hochachka and McClelland, 1997***), is comparable to the values that our model predicts are beneficial for ATP homeostasis. These results suggest that the role of Creatine and CK in ATP homeostasis could be driven by the control of the non-adenine phosphate pool instead of direct buffering of ATP levels, while our model does not support an important role for AK and NDPK in ATP homeostasis. We note that our model does not describe the spatial heterogeneity of ATP production and consumption that might lead to gradients of ATP inside cells, where local buffering of ATP levels by CK, AK, or NDPK might play an important role.

### Coarse-grained model of glycolysis recapitulates key tasks of ATP homeostasis

To gain a better understanding of the tasks of ATP homeostasis that can be achieved by mass action alone and which might require allostery, we used a simplified model of glycolysis, referred to as the two-enzyme model, containing ATP-consuming Enzyme 1, ATP-producing Enzyme 2 and ATPase, which represents upper glycolysis from HK to ALDO, lower glycolysis from GAPDH to LDH, and cellular ATPase activity, respectively (***Figure 7A***):

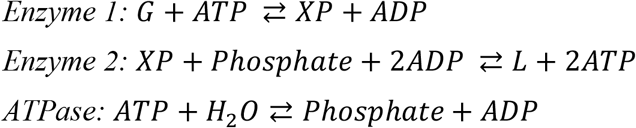

Note that metabolite XP, which represents all intracellular glycolysis intermediates, is phosphorylated to conserve phosphate in each step.

**Figure 7.**
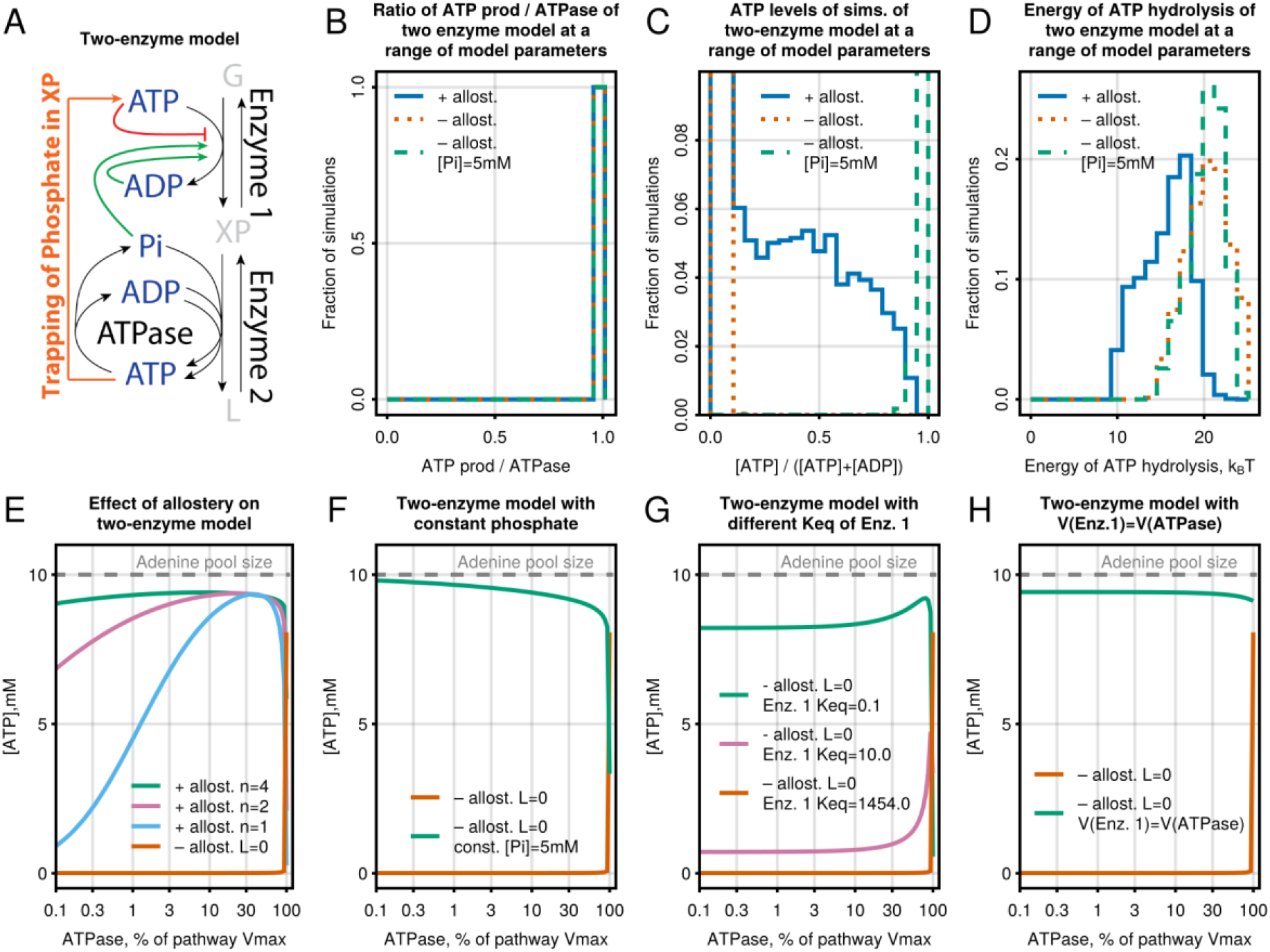
Coarse-grained model of glycolysis recapitulates key tasks of ATP homeostasis. **(A)** Schematic of a simplified glycolysis-like pathway containing two enzymes with feedback on Enzyme 1. **(B-D)** Simulation of the two-enzyme model at random values of the 6 parameters controlling this pathway showing the ability to (B) match ATP supply and demand, (C) maintain ATP levels in relation to ATP+ADP, and (D) generate energy from ATP hydrolysis. Random values for parameters were taken from log-uniform distributions in the interval of [10^−5^, 10^−1^] for *L*, [10^−3^, 10^−1^] for *Vmax*, and [10^1^, 10^4^] for *Keq*. Parameters controlling ATPase were not varied, and *n* was randomly chosen from a set [1, 2, 3, 4]. **(E-H)** Simulation of the two-enzyme with best-performing parameters from (C) showing maintenance of steady-state ATP levels at a 100x range of ATPase rates with (E) presence and absence of allostery, (D) constant inorganic phosphate levels, (G) different K_eq_ for Enzyme 1 and (H) rate of Enzyme 1 equal to rate of ATPase. Julia code to reproduce this figure is available at https://github.com/DenisTitovLab/Glycolysis.jl.

To simplify the analysis, we described the kinetics of Enzyme 2 and ATPase using modified reversible Michaelis-Menten equations with the assumption that Enzyme 2 and ATPase are saturated with substrates (***Noor et al., 2013***). For Enzyme 1, we used the MWC model to add allosteric activation by ADP and phosphate and allosteric inhibition by ATP. We assumed that ADP and Phosphate only bind to active conformation, ATP only binds to inactive conformation, inactive MWC conformation of Enzyme 1 is catalytically inactive, and regulatory and catalytic sites are saturated with regulators and substrates. The resulting two-enzyme model is described using the following rate equations (see ***Materials and Methods*** for additional detail on the derivation of the two-enzyme model rate equations):

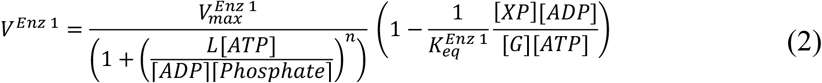

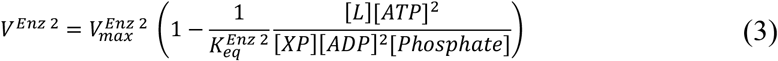

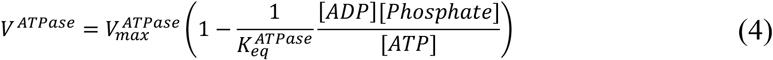

Two-enzyme model reduces the number of parameters from >150 for the full model to only 8 for the two-enzyme model, including only two parameters–*L* and *n–*responsible for allosteric regulation. *L* represents the ratio of active to inactive conformation of Enzyme 1 in absence of regulators and *n*–oligomeric state of Enzyme 1.

We searched for values of the parameters of the two-enzyme model that would support the three tasks of ATP homeostasis. We performed 10,000 simulations using random values of two-enzyme model parameters with or without allostery to see which combinations of parameters can support the three tasks at a range of ATPase values (***Figure 7B-D***). All the parameter combinations could match ATP supply and demand and maintain the energy of ATP hydrolysis >9 k_B_T in the presence or absence of allostery, confirming that mass action alone is fully capable of supporting these two tasks (***Figure 7B, D***). By contrast, only a small fraction of simulations in the presence of allostery and none of the simulation in absence of allostery could maintain ATP > ADP (***Figure 7C***). As for the full model (***Figure 5C***), keeping the phosphate level constant abrogated the requirement for allostery, confirming that maintenance of ATP > ADP is a non-trivial task that requires allostery to prevent the phosphate trap (***Figure 7C, F***).

We next performed simulations of the two-enzyme model with the parameters that supported the highest level of ATP at a range of ATPase values. We found that removing allostery by setting *L=0* completely abrogated while decreasing *n* from 4 to 1 attenuated ATP level maintenance (***Figure 7E***). The effect of *n* suggests an important physiological role for the oligomeric structure of allosteric enzymes like PFKP. As for the full model (***Figure 5D***), decreasing the equilibrium constant of Enzyme 1 abrogated the requirement for allostery as the trapping of phosphate in XP is less thermodynamically favorable under these conditions. It is noteworthy that the two-enzyme model without allostery could maintain ATP>ADP under one exact condition when the 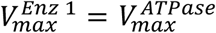 (red line in ***Figure 7E*** at 100% ATPase rate)–a condition when the trapping of phosphate would be impossible as all of the Enzyme 1 capacity is used to satisfy ATP demand with no spare capacity to catalyze the trapping reactions. As for the full model (***Figure 5E***), making 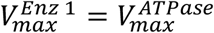 abrogated the requirement for allostery (***Figure 7H***). The latter again highlights that allostery prevents the trapping of phosphate by adjusting 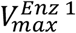 such that 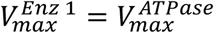 at a large range of 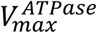.

Overall, our analysis of the two-enzyme model confirms the results of the full model that allosteric regulation is required to maintain high and stable ATP levels, while mass action alone is sufficient to match ATP supply and demand and to ensure that ATP hydrolysis generates energy.

## DISCUSSION

Metabolic homeostasis is an emergent property of the activities of many individual enzymes. Regulation of metabolic pathways has historically been studied by rigorously characterizing enzymes in isolation. While this has greatly advanced our understanding of the key regulatory enzymes, many emergent functions of metabolic networks are difficult to understand by focusing on one enzyme at a time. Approaches that combine theory and experiments can explore how the activities of individual enzymes give rise to metabolic homeostasis. Here, we developed a data-driven model of glycolysis that uses enzyme rate equations to describe the activity of the whole glycolysis pathway and applied it to identify the regulatory mechanisms that allow glycolysis to perform the three key tasks of ATP homeostasis (***Figure 1A***). The resulting model can quantitatively predict the output of glycolysis in response to perturbations of metabolite levels and kinetic properties of enzymes, such as changes in media conditions, enzyme expression, drug treatment, and expression of enzyme mutants.

One of the remarkable properties of ATP homeostasis is that it can maintain stable ATP levels even when a cell’s demand increases by up to 100-fold (***Hochachka and McClelland, 1997***). Given the near constancy of ATP, ADP, and phosphate levels under various conditions (***Allen et al., 1997; Balaban et al., 1986***), it has been debated whether ATP homeostasis is achieved through feedback sensing of changes in ATP, ADP, and phosphate levels that result from ATP consumption or whether additional mechanisms such as feedforward activation of ATP-producing enzymes by Ca^2+^ are required (***Balaban, 1990; Beard and Kushmerick, 2009***). The ability of our model to maintain stable ATP levels at >10-fold changes in the ATP demand suggests that known *in vitro* enzyme kinetic properties of glycolytic enzymes are already sufficient to maintain ATP homeostasis in response to changes in ATP, ADP, and phosphate concentrations due to ATP hydrolysis.

Glycolysis is regulated by a combination of mass action and allostery, but it is not fully understood which properties of ATP homeostasis are controlled by mass action versus allostery. We showed that mass action alone is sufficient to match ATP supply and demand, as well as to maintain ATP, ADP, and inorganic phosphate concentrations so that ATP hydrolysis generates energy (***Figure 3***, ***Figure 7***). The key role of mass action can be seen in the stoichiometry of the net reaction of glycolysis:

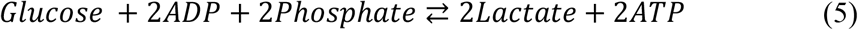

An increase in ATP consumption rate would push this reaction forward by mass action due to a decrease in the product (ATP) and an increase in substrate levels (ADP and phosphate) resulting from ATP hydrolysis. Likewise, the ATP hydrolysis reaction will be maintained far from equilibrium as long as the net reaction of glycolysis is thermodynamically favorable and can proceed faster than ATP hydrolysis. Thus, the ability of glycolysis to match ATP supply and demand and maintain ATP hydrolysis far from equilibrium is built into the glycolytic pathway, and it is largely independent of the kinetic properties of individual enzymes.

If mass action controls the rate of ATP production, what then is the function of allosteric regulation? We showed that the allosteric regulation of HK1 and PFKP by G6P, ATP, ADP, and Pi is required to maintain most of the adenine nucleotide pool in the form of ATP and prevent the uncontrolled accumulation of phosphorylated intermediates in upper glycolysis (***Figure 3***, ***Figure 5, Figure 7, Figure 5–figure supplement 1***). The latter is an especially difficult task due to the structure of the glycolysis pathway where an imbalance between the ATP-producing enzyme of lower glycolysis PGK and ATP-consuming enzymes of upper glycolysis HK1 and PFKP can trap phosphate in the upper glycolysis intermediates G6P, F6P, F16BP, GAP and DHAP (***Figure 5A***). This phosphate trap presents the cell with an impossible choice. If the cell stops the uptake of extracellular phosphate, then the concentration of phosphate and phosphorylated intermediates of lower glycolysis drop to low levels, necessitating high levels of ADP to maintain flux and making it impossible to maintain high ATP > ADP + AMP.

Conversely, if the cell continues to uptake extracellular phosphate, then G6P, F6P, and F16BP accumulate to toxic levels of > 100 M. Neither option is likely to be compatible with a living cell. We showed that allosteric regulation of HK1 and PFKP prevents phosphate trapping by tethering the rates of HK1 and PFKP to one-half of the ATPase rate such that there is no spare catalytic activity of HK1 and PFKP to catalyze the phosphate trapping reactions. These findings predict the specific outcomes of inactivation of allosteric regulation of HK and PFK that can be experimentally tested.

The phosphate trapping mechanism helps explain several previously puzzling observations. It has long been known that *S. cerevisiae* mutant Δtps1 lacking trehalose-6-phosphate (T6P) synthase is defective for growth on glucose (***Van Aelst et al., 1993***). In the presence of glucose, Δtps1 stops growing and accumulates large amounts of phosphorylated glycolytic intermediates upstream of GAPDH while having depleted levels of ATP, inorganic phosphate and phosphorylated glycolytic intermediates downstream of GAPDH (***Van Aelst et al., 1993***), as we predicted for a phosphate trap mechanism in the absence of allostery (***Figure 5B***). It turns out that *S. cerevisiae* hexokinase is feedback inhibited by T6P (***Blázquez et al., 1993***) instead of G6P and, thus, the Δtps1 mutant is likely defective in hexokinase feedback inhibition. Our results show that the lack of HK feedback inhibition is sufficient to break the maintenance of ATP levels and lead to trapping of inorganic phosphate, which would explain the unusual phenotype of Δtps1 mutant.

Several organisms have modified versions of HK1 and PFKP reactions with a phosphate donor other than ATP. For example, many bacteria use PEP to phosphorylate glucose into G6P using the phosphotransferase system (***Deutscher et al., 2006***), some archaea use ADP-dependent HK and PFK (***Guixé and Merino, 2009***), and select members of all three domains of life use pyrophosphate (PPi) to phosphorylate F6P using PPi-dependent PFK (***Reeves et al., 1974; Siebers et al., 1998***). Our results suggest that the function of these reactions might be to bypass the requirement for allosteric regulation by making phosphate trapping impossible, as free inorganic phosphate is transferred to ATP through sequential GAPDH and PGK reactions so that trapping can only occur if ATP is used by HK and PFK (***Figure 5A***). Thus, PEP-, PPi-, or ADP-dependent HK and PFK cannot participate in phosphate trapping since they do not use ATP as a phosphate donor. Observations that ADP-dependent HKs and PFKs (***Guixé and Merino, 2009***) and PPi-dependent PFKs (***Siebers et al., 1998***) are not allosterically regulated provide additional evidence that using phosphate donors other than ATP obviates the need for allosteric regulation.

Glycolytic enzymes are expressed at unusual levels, with GLUT, HK1, PFKP, and MCT having 10-100x less activity than most of the other enzymes (***Figure 6A, B***). Such an expression pattern is puzzling given that glycolysis cannot proceed faster than its slowest enzymes, so this 10-100x higher expression seems wasteful. Our results provide a mechanistic explanation for this unusual expression pattern by showing that it is required to maintain high and stable ATP levels and to prevent phosphate trapping (***Figure 6C***).

Beyond the applications described in this report, our model of glycolysis can be used for a variety of applications. The model can be used to predict the effect of enzyme mutants or simulate the effects of glycolysis-targeting drugs by changing the corresponding kinetic parameters. Global sensitivity analysis (***Figure 4–figure supplement 1***) can be used to identify key kinetic parameters that control the output of glycolysis, such as the tasks of ATP homeostasis or a concentration of a specific metabolite. The rate equations for glycolytic enzymes can be used to simulate reaction-diffusion systems and investigate the effect of the 3D distribution of glycolytic enzymes. The existence of novel regulators not included in the current model can be inferred by investigating the quantitative differences between model prediction and data.

Our framework to combine theory and experiments can be readily applied to other metabolic pathways. Enzyme rate equations can be derived from the vast literature of *in vitro* kinetics data, threading together the individual enzyme behaviors to generate the system’s behavior. Our approaches for simulations, global sensitivity analysis, and error propagation of systems of ODEs can be directly applied to other pathways without modification. As with glycolysis, we anticipate that such work will lead to a better understanding of the functions of mass action and allostery in the regulation of metabolic homeostasis.

## Supporting information

Supplementary File 1. Levels of enzymes, metabolites and isotope tracing

Supplementary File 2. Kinetic rate data for glycolytic enzymes

## FIGURE SUPPLEMENTS

**Figure 2–figure supplement 1.**
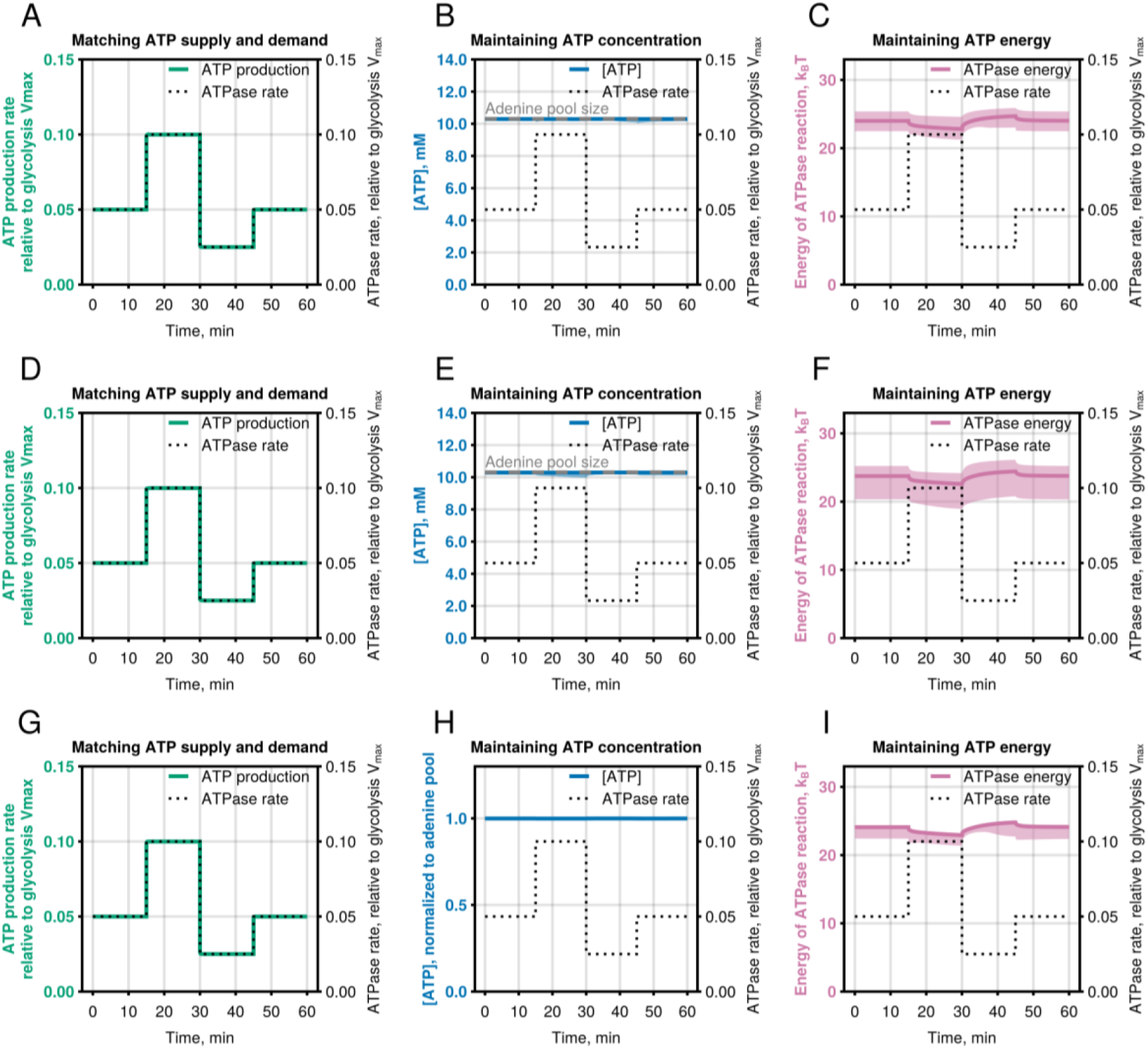
Related to Figure 2. Sensitivity of model predictions to enzyme levels and experimental error in biochemical constants, protein and cofactor pool concentrations, and initial concentrations of intermediates. **(A-I)** Model simulations showing median changes and 95% confidence interval of (A, D, G) ATP production rate, (B, E, H) ATP concentration (total adenine pool size is labeled with dashed grey line), and (C, F, I) energy released during ATP hydrolysis in response to a 2-fold stepwise increase or decrease of ATPase rate from 10,000 bootstrapped models with (A-C) enzyme concentrations drawn from a log-uniform distribution spanning 9-fold range around the model value, (D-F) parameters drawn from a normal distribution with mean and SD being equal to experimental mean and SEM for each parameter of the model, (G-I) initial concentration of metabolites taken from a normal distribution with mean and SD being equal to experimental mean and SEM for each metabolite level. Blue line shows median value at each ATPase rate and blue ribbon is 95% confidence interval. ATPase energy is calculated as a natural logarithm of the disequilibrium ratio for the ATPase reaction (i.e., mass-action ratio divided by the equilibrium constant) for the ATPase reaction. Julia code to reproduce this figure is available at https://github.com/DenisTitovLab/Glycolysis.jl.

**Figure 3–figure supplement 1.**
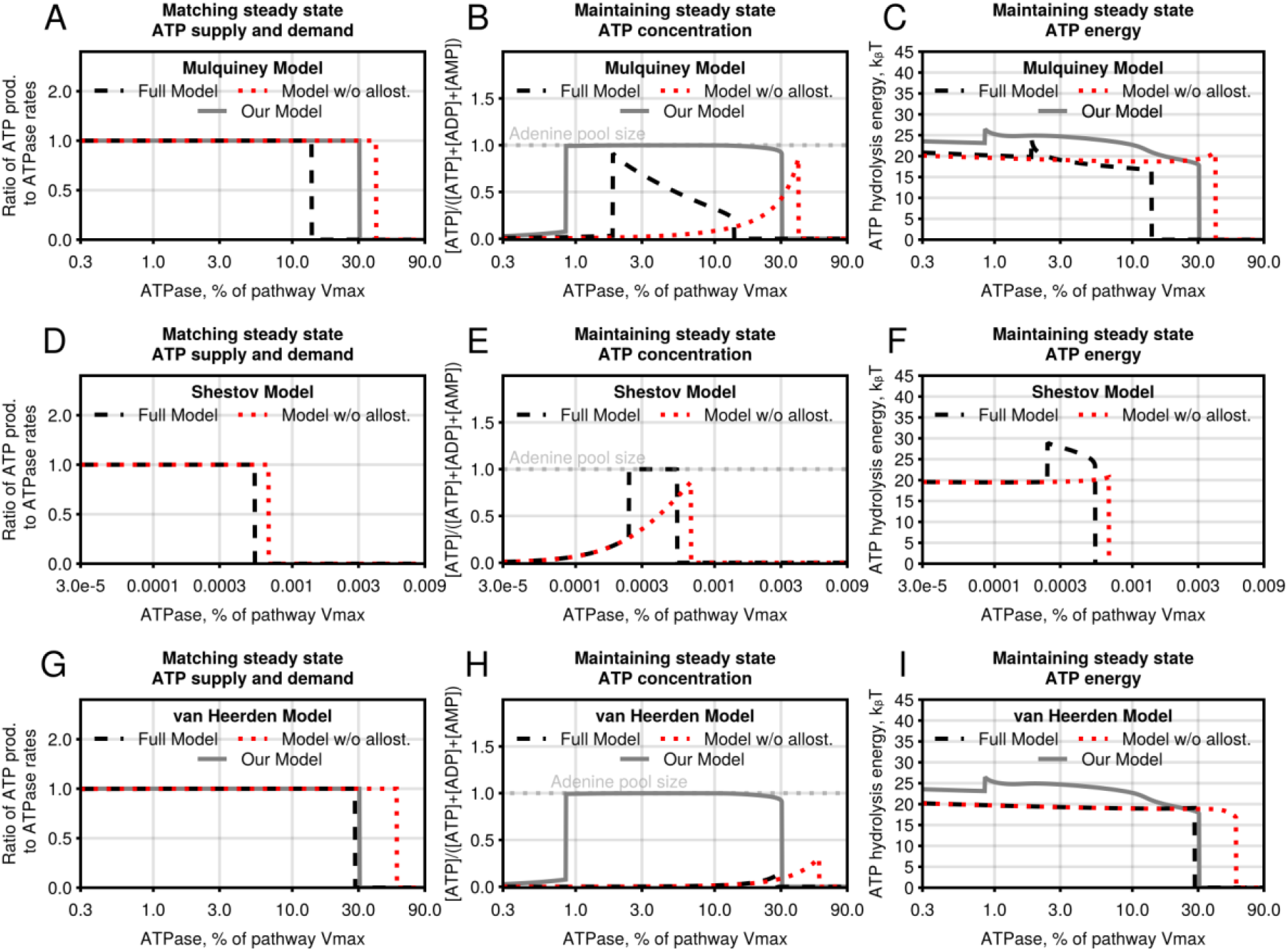
Related to Figure 3. ATP homeostasis maintenance by our glycolysis model and several previously reported models. **(A-I)** Comparison of the ability of Mulquiney model (A-C), Shestov model (D-F), and van Heerden model (G-I) to (A, D, G) match ATP supply and demand, (B, E, H) maintain ATP concentration, and (C, F, I) maintain energy released during ATP hydrolysis at a range of ATPase rates in the presence or absence of allosteric regulation. Results are steady-state values determined by simulating the models for 10^8^ seconds at each ATPase rate. Note that the Shestov model can only support ATPase rates that are 10,000 times lower than other models in relation to the respective model Vmax and, thus, plotted with different x-axis scale. Energy is calculated as a natural logarithm of the ratio of mass balance to the thermodynamic equilibrium constant Keq for the ATPase reaction. The total adenine pool size (i.e., ATP + ADP + AMP) is highlighted with a dashed grey line in (B, E, H). Julia code to reproduce this figure is available at https://github.com/DenisTitovLab/Glycolysis.jl.

**Figure 4–figure supplement 1.**
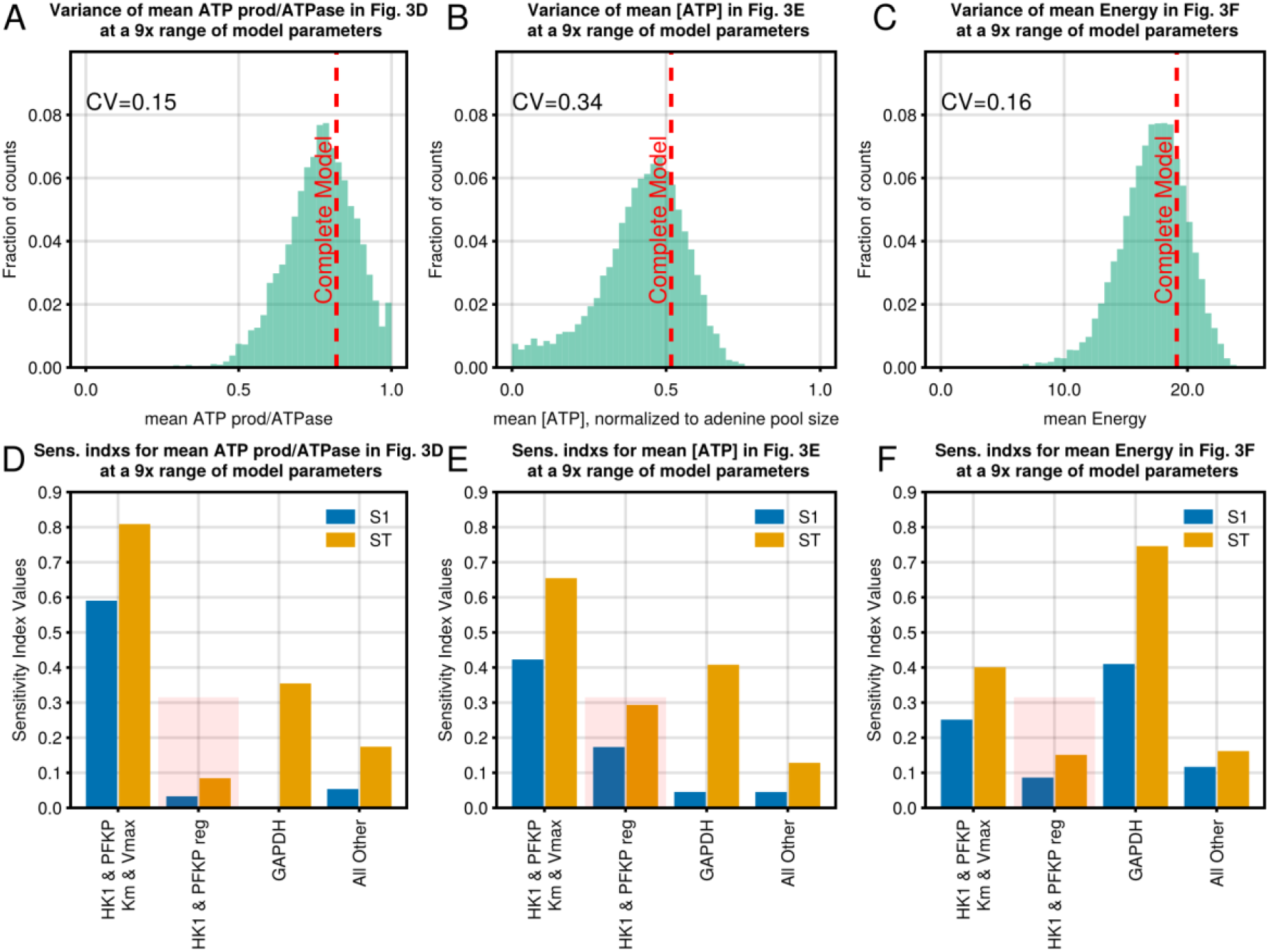
Related to Figure 4. Global sensitivity analysis of the model’s ability to maintain ATP level, ATPase energy, and ATP production. **(A-C)** Distribution of (A) mean ATP production to ATPase rate ratios, (B) mean ATP concentrations normalized by adenine pool size, (C) mean ATP hydrolysis energies from a simulation like in ***Figure 3D-F***, respectively, repeated 10,000 times using model parameters drawn from a log-uniform distribution spanning 9-fold range around their model values. **(D-F)** Sensitivity indexes of glycolysis model parameters that explain the variance of (D) mean ATP production to ATPase rate ratios, (E) mean ATP concentrations normalized by adenine pool size, (F) mean ATP hydrolysis energies, shown in Panels (A-C), respectively. Julia code to reproduce this figure is available at https://github.com/DenisTitovLab/Glycolysis.jl.

**Figure 5–figure supplement 1.**
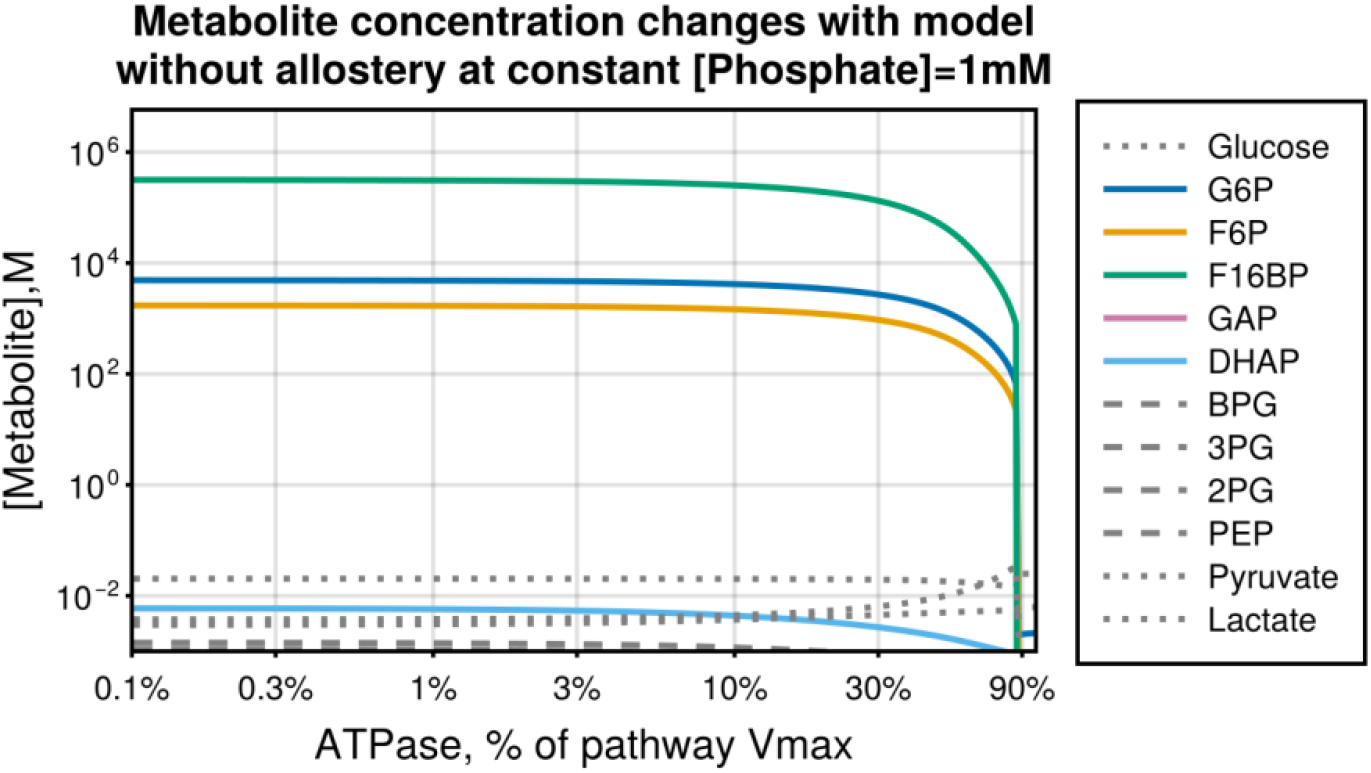
Related to Figure 5. Maintaining constant inorganic phosphate in absence of allostery leads to the accumulation of phosphorylated upper glycolysis intermediates to levels incompatible with viable cells. Steady-state metabolite concentrations at a range of ATPase rates of the glycolysis model without allosteric regulation and with constant [Phosphate] = 1mM. Same conditions as ***Figure 5C***. Note that G6P, F6P, F16BP are not in steady state and will continue to accumulate if simulation time is increased. Phosphorylated metabolites before GAPDH are color lines, phosphorylated metabolites after GAPDH are grey dashed lines, and unphosphorylated metabolites are dotted lines. Julia code to reproduce this figure is available at https://github.com/DenisTitovLab/Glycolysis.jl.

**Figure 6–figure supplement 1.**
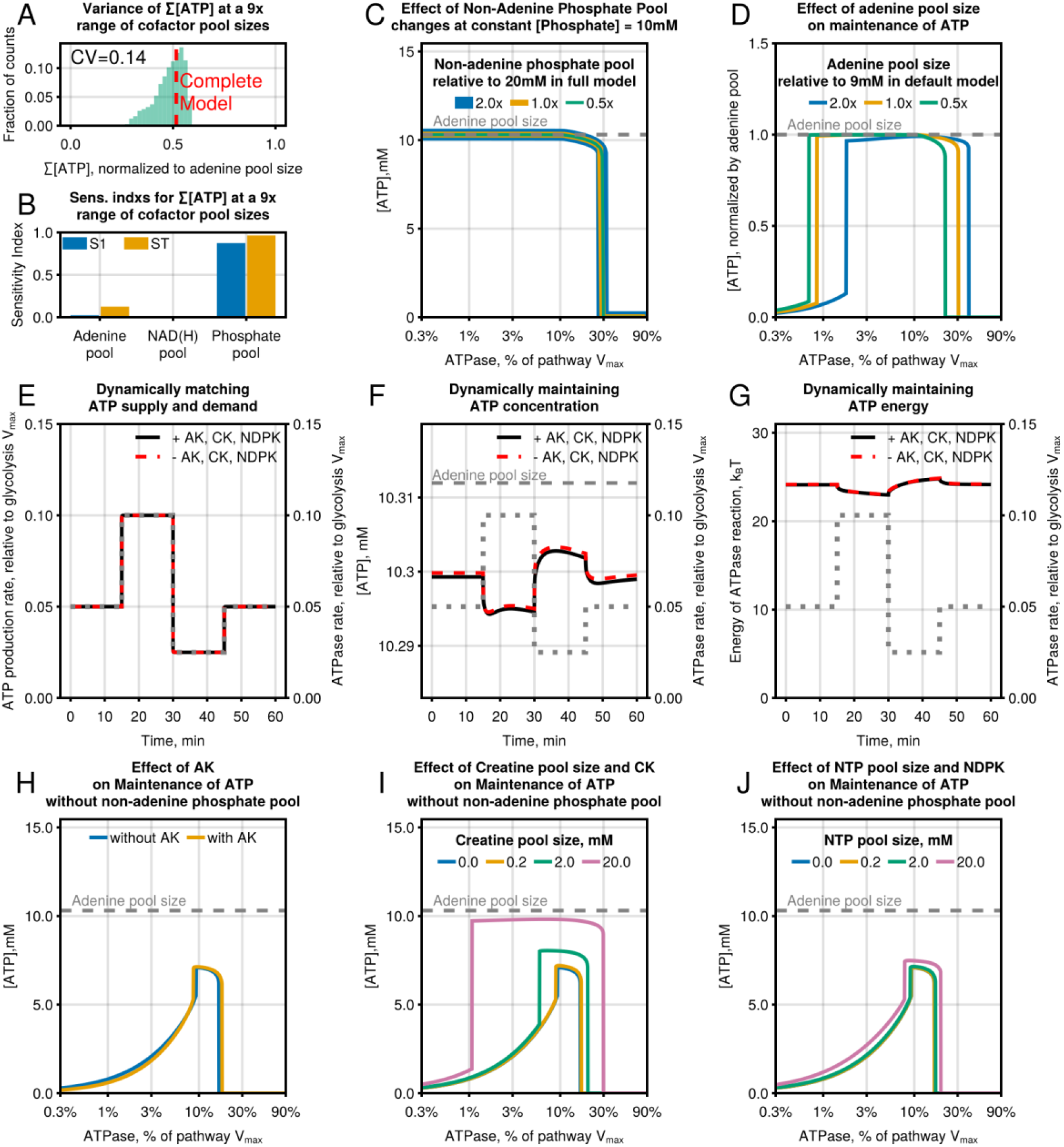
Related to Figure 6. Global sensitivity analysis of the effect of cofactor pool sizes on ATP level maintenance. **(A)** Distribution of the mean ATP concentrations normalized by adenine pool size from a simulation like in ***Figure 3E*** repeated 10,000 times using model parameters drawn from a log-uniform distribution spanning 9-fold range of adenine pool (i.e., ATP + ADP + AMP), NAD(H) pool (i.e., NAD^+^ + NADH), and non-adenine phosphate pool (i.e., sum of all phosphate groups except for ATP, ADP, and AMP) sizes around their model values. **(B)** Sensitivity indexes of two enzyme model parameters that explain the variance of ratio of ATP to ATP+ADP shown in Panel A. Error bars are 95% CI of sensitivity indexes. **(C)** Effect of changes of non-adenine phosphate pool size on model’s ability to maintain ATP levels at a fixed concentration of Phosphate = 10 mM. **(D)** Steady-state ATP concentrations at a range of ATPase rates of the glycolysis model with different levels of adenine pool size. **(E-G)** Model simulations with or without AK, CK, and NDPK showing changes in (E) ATP production rate, (F) ATP concentration (total adenine pool size is labeled with dashed grey line), and (G) energy released during ATP hydrolysis in response to a 2-fold stepwise increase or decrease of ATPase rate. Same conditions as ***Figure 2A-C*. (H-J)** Steady-state ATP concentrations at a range of ATPase rates of the glycolysis model (H) in the presence and absence of AK, (I) with different levels of creatine pool size in the presence of CK, and (J) with different levels of NTP pool size in the presence of NDPK. The total adenine pool size (i.e., ATP + ADP + AMP) is highlighted with a dashed grey line. Julia code to reproduce this figure is available at https://github.com/DenisTitovLab/Glycolysis.jl.

## ADDITIONAL FILES

Two Excel Spreadsheets with data are provided “*Supplementary File 1. Intracellular concentration of glycolytic enzymes, metabolites and isotope tracing data*.*xlsx”* and *“Supplementary File 2. Kinetic rate data for glycolytic enzymes*.*xlsx”* that contain all of the data used in this manuscript. All of the code required to reproduce all the figures is publicly available at https://github.com/DenisTitovLab/Glycolysis.jl.

## ACKNOWLEDGEMENTS

We thank the students and instructors of the 2017 Marine Biological Laboratory course on Physical Biology of the Cell for their comments on the early version of this project; participants of the 2019 Kavli Institute for Theoretical Physics workshop on Cellular Energetics for fruitful discussions; Bradley Webb for discussions about PFK regulation; James Mbata for contributions in identifying code errors; and members of Titov and Phillips labs for many helpful suggestions. This research used the Savio computational cluster resource provided by the Berkeley Research Computing program at the University of California, Berkeley (supported by the UC Berkeley Chancellor, Vice Chancellor for Research, and Chief Information Officer). Research reported in this publication was supported by the National Institute of General Medical Sciences of the National Institutes of Health under Award Numbers DP2 GM132933 and R35 GM152114 to D.V.T, and R35 GM118043 to R.P. T.E. is supported by Damon Runyon Cancer Research Foundation Fellowship DRQ 01-20.

## AUTHOR CONTRIBUTIONS

Conceptualization: RP, DVT

Data curation: MC, TE, DVT

Methodology: MC, TE, DVT

Investigation: MC, TE, DVT

Funding acquisition: RP, DVT

Supervision: RP, DVT

Writing – original draft: DVT

Writing – review & editing: MC, TE, RP, DVT

## MATERIALS AND METHODS

### Estimation of intracellular concentrations of glycolytic enzymes

We estimated the abundance of glycolytic enzymes in mammalian cell lines using publicly available proteomics data (***Geiger et al., 2012b***). We first used proteomics data to calculate the fraction a specific glycolytic enzyme occupies compared to total proteome size as a ratio of peptide intensities for the specific glycolytic enzyme to total peptide intensity for all proteins. This approach has been shown to produce good estimates of absolute protein amounts (***Wiśniewski et al., 2014***). Values of mg protein per mg total protein were then converted to concentration using the molecular weight of each enzyme and the cellular protein density of ∼200 mg/mL (***Milo, 2013***). The resulting dataset of glycolytic enzyme concentrations is reported in ***Supplementary File 1***. In the model simulations, we further converted whole-cell enzyme concentrations from ***Supplementary File 1*** to cytosolic enzyme concentrations as glycolytic enzymes are localized to the cytosol. We estimated that the enzymes of glycolysis are concentrated in a volume that is ∼50% of cellular volume, as ∼70% of the cell is cytosol (***Heinrich et al., 2021***) and ∼70% of the cytosol is water (***Ellis, 2001; Luby-Phelps, 2000***).

### Measurement of intracellular glycolytic intermediate concentrations using LC-MS

LC/MS was used to profile and quantify the polar metabolite contents of whole-cell samples. The metabolite extraction buffer was composed of 40% methanol (Fisher Scientific, Cat# A456-4) and 40% acetonitrile (Fisher Scientific, Cat# A9554) in water (Fisher Scientific, Cat# W64) containing 0.1 M formic acid (Fisher Scientific, Cat# A11750), supplemented with a mixture of 8 isotope-labeled chemicals at 1 μM (ATP (Sigma, Cat# 710695), AMP (Sigma, Cat# 900382), glucose-6-phosphate (Cambridge Isotope Laboratories, Inc., Cat# CLM-8367) and fructose-6-phosphate (Cambridge Isotope Laboratories, Inc., Cat# CLM-8616)), 10 μM (lactate (Sigma, Cat# 490040), pyruvate (Sigma, Cat# 490709) and fructose-1,6-bisphosphate (Cambridge Isotope Laboratories, Inc., Cat# CLM-3398)) or 20 μM (glucose (Sigma, Cat# 389374)), which were used as internal standards.

Metabolite analysis was performed on an Agilent 6430 Triple Quad interfaced with an Agilent 1260 Infinity II LC system. 5 μL of each metabolite sample was injected onto a SeQuant ZIC-pHILIC column 5 μm polymer 150 × 2.1 mm (Millipore/Sigma, Cat# 1504600001). Buffer A was 100% acetonitrile; buffer B was 20 mM ammonium carbonate, 0.1% ammonium hydroxide. The chromatographic gradient was run at a flow rate of 0.150 mL/min as follows: 0-0.5 min: hold at 80% A; 0.5-30.5 min: linear gradient from 80% to 20% A; 30.5-30.8 min: hold at 20% A; 30.8-31 min: linear gradient from 20% to 80% A; 31-60 min: hold at 80% A.

The MS settings were kept consistent regardless of the chromatographic separation being tested. MS parameters were as follows: gas, 350 °C at 11 L/min; Nebulizer, 35 psi at 4000 V. The MS was operated in negative ionization mode for all samples analyzed.

All experiments were performed in replicates of five (n=5) per sample group. Metabolite identification and quantification were performed with the Agilent MassHunter Qualitative Analysis software (version B.06.00). To confirm metabolite identities and to enable quantification, the pools of metabolite standards were used. To accurately quantify target metabolites, the final concentrations of standards were 100 μM, 50 μM, 25 μM, 12.5 μM, 6.25 μM, 3.125 μM, 1.56 μM, 781 nM, 390 nM, 195 nM, 97 nM, and 49 nM. In each sample, the raw peak area of each metabolite was divided by the raw peak area of the relevant isotope-labeled internal standard to calculate the absolute concentration. ATP, AMP, glucose-6-phosphate, fructose-1,6-bisphosphate, pyruvate, and lactate were normalized with their isotope-labeled counterparts. For determining the absolute concentrations of all other metabolites, the peak areas were normalized with 2 isotope-labeled internal standards which have the closest retention time. The raw peak area values were fit to a linear fitting curve equation, typically with r^2^ > 0.99, which was then used to calculate the concentration of the metabolite in each extract. The final intracellular concentrations of target metabolites were then calculated from the sample dilution-fold and the corresponding cell volume, which was estimated using the Beckman Z2 Coulter Counter.

Two sets of isomers, fructose-6-phosphate/glucose-1-phosphate (F6P/G1P) and 3-phosphoglycerate/2-phosphoglycerate were not separated under our chromatographic conditions. We took advantage of the different ratios of the signal from two transition states for each of the isomers to deconvolve the peaks. For example, F6P and G1P generate different signal ratios from two transition states with product ion values of 78.96 or 96.97. We used F6P and G1P standards to determine the ratios of signal from 78.96 product ion to that of 96.97 product ion and used these values to deconvolve the peaks containing the mixture of the two compounds.

One million C2C12 or HeLa cells were seeded on 6-well plates in 2 mL of DMEM without pyruvate (US Biological, Cat# D9802-25L), supplemented with 25 mM glucose, 3.7 g/L NaHCO_3_ and 10% FBS (Gibco, Cat# 10437028). 24 hr later, the medium was exchanged to 2 mL of DMEM medium without pyruvate, supplemented with 25 mM glucose, 3.7 g/L NaHCO_3_, 10% FBS, and 1 μM oligomycin (MP biomedicals, Cat# 151786). Metabolite extraction was performed on ice. After 2 hr incubation, the cell culture plate was placed on ice and the medium was aspirated. 150 μL of ice-chilled extraction buffer (40:40:20 = methanol:acetonitrile:water) containing 0.1 M formic acid and 8 isotope-labeled internal standards was added and cells were scraped with a cell lifter for 10 s. The lysate was transferred to a 1.5-mL tube on ice. Five minutes later, cells were centrifuged at 17000 *g* at 4 °C for 10 min. Supernatants were collected and neutralized by adding ammonium bicarbonate (final concentration of 100 mM). These samples were analyzed with LC/MS as described above to determine intracellular concentrations of metabolites.

For each experiment, a replicate set of cells was treated identically in parallel and used for measuring cell number and volume. The contents of each well were trypsinized, and cell number and volume were measured using a Beckman Z2 Coulter Counter with a size setting of 492.8-23620 fL. Intracellular concentrations were calculated using the total number of cells per sample and the volume of each cell.

### Estimates of glycolytic intermediate concentrations from the literature

In addition to our own measurements of glycolytic intermediate concentrations, we have compiled a dataset of mammalian glycolytic concentrations measured under different conditions from eight publications (***Chen et al., 2016; Edwards and Palsson, 2000; Mulquiney and Kuchel, 1999; Park et al., 2016; Røst et al., 2020; Shestov et al., 2014; Xie et al., 2016; Yurkovich et al., 2017***). The resulting dataset combined with our own measurement is reported in ***Figure 2D*** and provided in ***Supplementary File 1***. In ***Figure 2D***, we further converted whole-cell metabolite concentrations from ***Supplementary File 1*** to cytosolic metabolite concentrations as glycolysis is localized to the cytosol. We estimated that the metabolites of glycolysis are concentrated in a volume that is ∼50% of cellular volume, as ∼70% of the cell is cytosol (***Heinrich et al., 2021***) and ∼70% of the cytosol is water (***Ellis, 2001; Luby-Phelps, 2000***).

### Measurement of glycolytic fluxes using [^13^C]Glucose and [^13^C]Lactate

C2C12 or HeLa cells (1 million) were seeded on 6-well plates in 2 mL of DMEM without pyruvate, supplemented with 25 mM glucose, 3.7 g/L NaHCO_3_ and 10% FBS. 24 hr later, the medium was exchanged to 2 mL of DMEM medium without pyruvate, supplemented with 25 mM glucose, 3.7 g/L NaHCO_3_, 10% FBS, and 1 μM oligomycin. After 2 h incubation, the medium was exchanged to 2 mL of DMEM medium without pyruvate, supplemented with 25 mM ^13^C_6_-glucose(Sigma, Cat# 389274-1G), 3.7 g/L NaHCO_3_, 10% dialyzed FBS (ThermoFisher, Cat# 26400044), and 1 μM oligomycin or 2 mL of DMEM medium without pyruvate, supplemented with 25 mM glucose, 5 mM ^13^C_3_-lactate (Cambridge isotope laboratory, Cat# CLM-1579-N-0.1MG), 3.7 g/L NaHCO_3_, 10% dialyzed FBS and 1 μM oligomycin. After 1, 3, 10, and 30 min, cell culture plates were place on ice and medium was aspirated. Cells were rinsed with 2 mL of ice-cold 100 mM ammonium bicarbonate for no more than 10 s and metabolites were extracted as described above using extraction buffer (40:40:20 = methanol:acetonitrile:water) containing 0.1 M formic acid and 1 μM isotopic AMP. In each sample, the raw peak area of each metabolite was divided by the raw peak area of isotope-labeled AMP internal standard to calculate the relative abundance.

### Numerical simulations of differential equations constituting the Glycolysis Model

The glycolysis model is a system of ordinary differential equations (ODEs). The number of equations is equal to the number of metabolites, and each equation consists of a derivative of metabolite concentration with respect to time being equal to the sum of enzyme rates producing the metabolite minus the sum of enzyme rates consuming the metabolite:

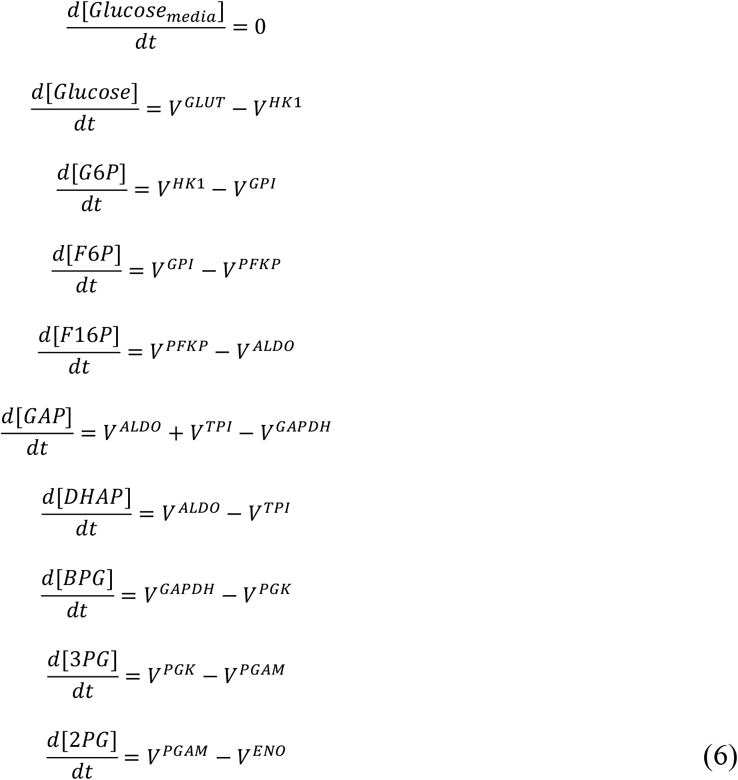

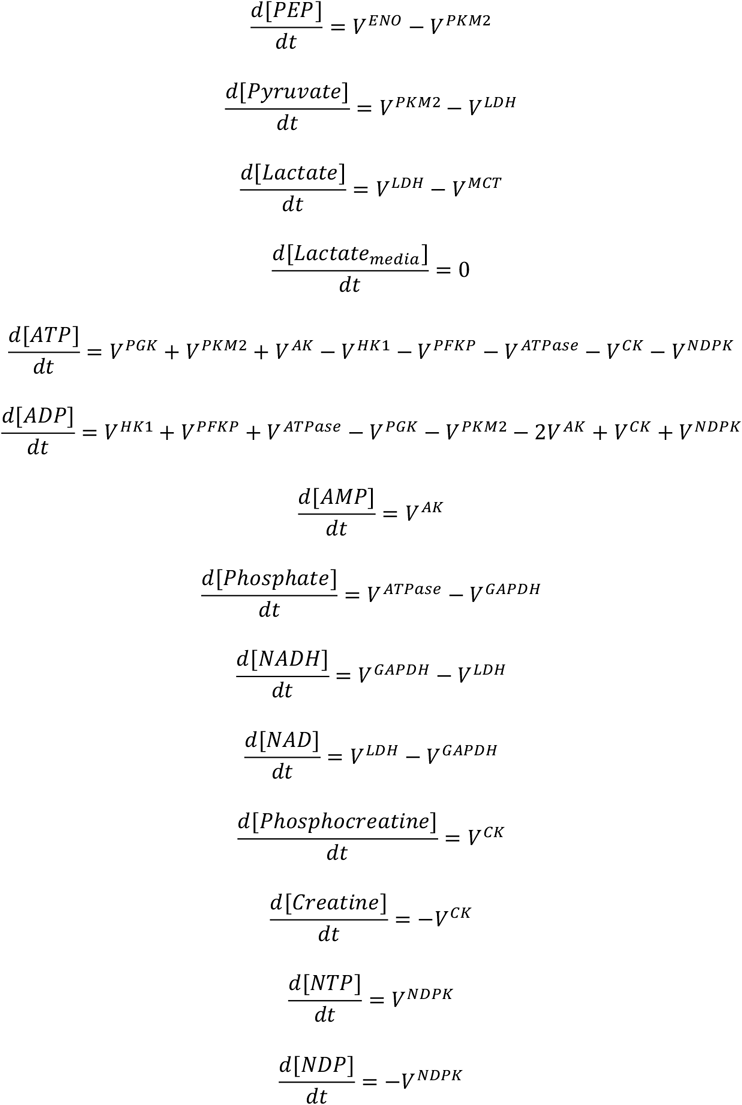

Note that NTP, NDP, Phosphocreatine, and Creatine are only included in the simulations of ***Figure 6F*** and ***Figure 6–figure supplement 1E-G***. The rate of CK and NDPK are zero in other simulations.

The ODE model can also be represented in the form of the stoichiometric matrix:

|  | GLUT | HK1 | GPI | PFKP | ALDO | TPI | GAPDH | PGK | PGM | ENO | PKM2 | LDH | MCT | ATPase | AK | CK | NDPK |
| --- | --- | --- | --- | --- | --- | --- | --- | --- | --- | --- | --- | --- | --- | --- | --- | --- | --- |
| Media Glucose | 0 | 0 | 0 | 0 | 0 | 0 | 0 | 0 | 0 | 0 | 0 | 0 | 0 | 0 | 0 | 0 | 0 |
| Glucose | 1 | -1 | 0 | 0 | 0 | 0 | 0 | 0 | 0 | 0 | 0 | 0 | 0 | 0 | 0 | 0 | 0 |
| G6P | 0 | 1 | -1 | 0 | 0 | 0 | 0 | 0 | 0 | 0 | 0 | 0 | 0 | 0 | 0 | 0 | 0 |
| F6P | 0 | 0 | 1 | -1 | 0 | 0 | 0 | 0 | 0 | 0 | 0 | 0 | 0 | 0 | 0 | 0 | 0 |
| F16BP | 0 | 0 | 0 | 1 | -1 | 0 | 0 | 0 | 0 | 0 | 0 | 0 | 0 | 0 | 0 | 0 | 0 |
| GAP | 0 | 0 | 0 | 0 | 1 | 1 | -1 | 0 | 0 | 0 | 0 | 0 | 0 | 0 | 0 | 0 | 0 |
| DHAP | 0 | 0 | 0 | 0 | 1 | -1 | 0 | 0 | 0 | 0 | 0 | 0 | 0 | 0 | 0 | 0 | 0 |
| BPG | 0 | 0 | 0 | 0 | 0 | 0 | 1 | -1 | 0 | 0 | 0 | 0 | 0 | 0 | 0 | 0 | 0 |
| 3PG | 0 | 0 | 0 | 0 | 0 | 0 | 0 | 1 | -1 | 0 | 0 | 0 | 0 | 0 | 0 | 0 | 0 |
| 2PG | 0 | 0 | 0 | 0 | 0 | 0 | 0 | 0 | 1 | -1 | 0 | 0 | 0 | 0 | 0 | 0 | 0 |
| PEP | 0 | 0 | 0 | 0 | 0 | 0 | 0 | 0 | 0 | 1 | -1 | 0 | 0 | 0 | 0 | 0 | 0 |
| Pyruvate | 0 | 0 | 0 | 0 | 0 | 0 | 0 | 0 | 0 | 0 | 1 | -1 | 0 | 0 | 0 | 0 | 0 |
| Lactate | 0 | 0 | 0 | 0 | 0 | 0 | 0 | 0 | 0 | 0 | 0 | 1 | -1 | 0 | 0 | 0 | 0 |
| Media Lactate | 0 | 0 | 0 | 0 | 0 | 0 | 0 | 0 | 0 | 0 | 0 | 0 | 0 | 0 | 0 | 0 | 0 |
| ATP | 0 | -1 | 0 | -1 | 0 | 0 | 0 | 1 | 0 | 0 | 1 | 0 | 0 | -1 | 1 | -1 | -1 |
| ADP | 0 | 1 | 0 | 1 | 0 | 0 | 0 | -1 | 0 | 0 | -1 | 0 | 0 | 1 | -2 | 1 | 1 |
| AMP | 0 | 0 | 0 | 0 | 0 | 0 | 0 | 0 | 0 | 0 | 0 | 0 | 0 | 0 | 1 | 0 | 0 |
| Phosphate | 0 | 0 | 0 | 0 | 0 | 0 | -1 | 0 | 0 | 0 | 0 | 0 | 0 | 1 | 0 | 0 | 0 |
| NTP | 0 | 0 | 0 | 0 | 0 | 0 | 0 | 0 | 0 | 0 | 0 | 0 | 0 | 0 | 0 | 0 | 1 |
| NDP | 0 | 0 | 0 | 0 | 0 | 0 | 0 | 0 | 0 | 0 | 0 | 0 | 0 | 0 | 0 | 0 | -1 |
| Phosphocreatine | 0 | 0 | 0 | 0 | 0 | 0 | 0 | 0 | 0 | 0 | 0 | 0 | 0 | 0 | 0 | 1 | 0 |
| Creatine | 0 | 0 | 0 | 0 | 0 | 0 | 0 | 0 | 0 | 0 | 0 | 0 | 0 | 0 | 0 | -1 | 0 |
| NAD | 0 | 0 | 0 | 0 | 0 | 0 | -1 | 0 | 0 | 0 | 0 | 1 | 0 | 0 | 0 | 0 | 0 |
| NADH | 0 | 0 | 0 | 0 | 0 | 0 | 1 | 0 | 0 | 0 | 0 | -1 | 0 | 0 | 0 | 0 | 0 |

The stoichiometric matrix allows us to calculate the number of conserved moieties and identify them using chemical reactions network theory. The number of conserved moieties is the number of metabolites (i.e., rows) minus the rank of the stoichiometric matrix, and the conserved moieties are the left null space of the stoichiometric matrix. The 8 conserved moieties are:

1. Adenine pool: ATP + ADP + AMP
2. NAD(H) pool: NAD + NADH
3. Media Glucose
4. Media Lactate
5. Total phosphate pool: NTP + PEP + TwoPG + DHAP + Phosphate + GAP + 2.0F16BP + Phosphocreatine + ADP + ThreePG + F6P + G6P + 2.0BPG + 2.0ATP
6. Redox state of intermediates: NADH – Pyruvate – PEP – TwoPG – ThreePG – BPG
7. Creatine pool: Creatine + Phosphocreatine
8. NTP pool: NDP+NTP.

Note that 7 and 8 are only included in the simulations of ***Figure 6F*** and ***Figure 6–figure supplement 1E-G***.

We numerically simulated ***Equations (6)*** using DifferentialEquations.jl library (***Rackauckas and Nie, 2017***). We used Rodas5P solver for stiff differential equations with 10^−15^ absolute tolerance and 10^−8^ relative tolerance for most simulations and RadauIIA5 solver for simulations with constant phosphate and simulations where rate equations for HK1 and PFKP were substituted for one-half of ATPase rate equations. Tolerances were chosen to provide minimal numerical errors estimated by Uncertainty Quantification callback AdaptiveProbIntsUncertainty from DiffEqCallbacks.jl library. Numerical simulation of ODEs requires an initial condition consisting of metabolite concentration at time 0. We used estimates of intracellular concentrations of glycolytic intermediates as initial conditions (see ***Supplementary File 1***) or a steady-state solution for dynamic simulations with step responses to changes in ATPase rate. A large range of initial conditions, including all metabolites being zero except for ATP, NAD, and extracellular glucose, produced similar steady-state solutions as shown in ***Figure 2–figure supplement 1G-I***. We simulated differential equations for 10^8^ seconds (∼3.2 years) to find the steady-state concentrations of metabolites. Typically, the solution was close to steady state after only several seconds in accordance with a rapid rate of glycolysis observed in live cells. DifferentialEquations.jl has callback capabilities, which allowed us to change any parameter of the model (e.g., ATPase rate or concentration of any metabolite) at any time during the simulation. A typical simulation took 2-20 milliseconds on a single core of Apple M1 Max processor, which allowed us to test many different conditions in a short period of time using a personal computer.

### Calculation of ATP turnover, Energy released by ATP hydrolysis, Bound and Free Metabolite Concentrations

The output of the simulation of ***Equations (6)*** is the free concentrations of metabolites at every timepoint starting from initial conditions. This output can be used to calculate many properties of the glycolysis pathway, such as rates of any reaction or concentrations of bound metabolites, by plugging metabolite concentration into enzyme rate equations described in ***Appendix 1***.

Consumption and production of ATP was calculated using the following equations:

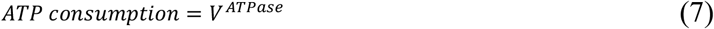

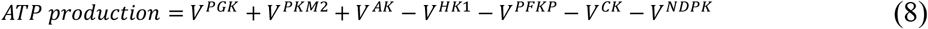

Note that the rates of CK and NDPK are zero in all simulations except those in ***Figure 6F*** and ***Figure 6–figure supplement 1E-G***.

Energy released by ATP hydrolysis in units of k_B_T was calculated using the following equation:

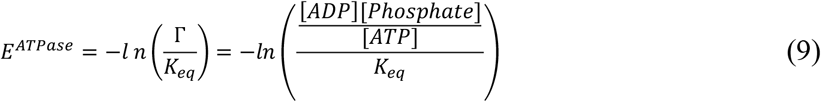

where Γ is the mass action ratio of ATP hydrolysis and *K*_*eq*_ is the equilibrium constant of ATP hydrolysis.

The concentration of enzyme-bound metabolites was calculated using binding equations described in the ***Appendix 1***. Accounting for enzyme-bound metabolites is essential to ensure an accurate comparison of model prediction to whole-cell measurements as the latter represents a sum of enzyme-bound and free metabolites. Intracellular concentrations of glycolytic enzymes are 1-10 µM, which is comparable to intracellular concentrations of some glycolytic intermediates. Therefore, some of the metabolites are mostly present in an enzyme-bound state, and their concentrations will be underestimated by the model without accounting for the enzyme-bound fraction.

### Estimation of Uncertainty of Model Predictions

Outputs of the glycolysis model have uncertainty due to uncertainty of estimates of model parameters such as kinetic constants, thermodynamic constants, and enzyme concentrations. Understanding the uncertainty of model predictions is important for quantitative comparisons of model predictions to data and for understanding the robustness of model predictions. We used bootstrapping to calculate the uncertainty of glycolysis model predictions. We rerun the model 10,000 times using different sets of parameter values, where each parameter value is generated by randomly drawing a value from a normal distribution with a mean and standard deviation corresponding to the mean and standard deviation of that parameter estimate. We bootstrapped all the parameters except we kept the kinetic parameters of ATPase and Adenylate Kinase unchanged during bootstrapping as these enzymes are not part of glycolysis and we set Vmax of ATPase to be equal to a fraction of Vmax of HK1 or Vmax PFKP whichever was smaller in a particular bootstrapped parameter set. In most analyses, we only reported the data from bootstrapped simulations that could match ATP supply and demand as indicated in relevant figure legends because including simulations that come to equilibrium due to the inability to produce ATP fast enough is not informative as they represent a completely different state of the glycolysis system akin to a dead cell. In summary, this general bootstrapping approach allows us to estimate the uncertainty of any output of our model.

### Global Sensitivity Analysis

Sensitivity analysis is a set of powerful approaches aimed to systematically explore the role of specific parameters of an ODE model in controlling specific outputs of the model, such as maintenance of high ATP level at a range of ATP turnovers, prediction of metabolite concentration, speed of the response to perturbation, etc. Sensitivity analysis can be broadly subdivided into local sensitivity analysis and global sensitivity analysis methods. Local sensitivity analysis methods using partial derivates were historically more popular, especially for simple models where analytical derivatives can be calculated. When applied to models of metabolic pathways, local sensitivity analysis is sometimes referred to as metabolic control analysis (***Heinrich and Rapoport, 1974; Kacser and Burns, 1973***). The limitations of local sensitivity analysis are that it only considers the effect of one parameter at a time (i.e., ignores interactions between parameters) and only estimates the role of the parameter at a specific value of that parameter and every other parameter (i.e., local). To overcome these limitations, global sensitivity analysis methods were developed that allow systematic evaluation of parameter importance at a large range of parameter values (i.e., global) and that can estimate the effect of interactions between parameters. The only limitation of variance decomposition methods is that they are much more computationally intensive than local sensitivity analysis methods, which limited their use before early 2000s when the computational power have caught up to the demands of these methods.

We used a global sensitivity analysis method called variance decomposition to learn about the role of specific model parameters in controlling specific outputs of our model. Here we briefly describe the idea behind this method and refer the reader to several excellent books that have been written about global sensitivity analysis for more in-depth discussion (***Saltelli, 2008; Saltelli et al., 2004***). First, we chose a scalar output *Y* of the model that we want to investigate. We focused on investigating the model parameters required to maintain high ATP concentration, high energy of ATP hydrolysis, and the ability of model to match ATP production and ATP consumption rates. Then we perform repeated simulations of the model using different starting parameters drawn from a log-uniform distribution with values 3-fold lower to 3-fold higher than the model values. We then calculated how much each parameter or combination of parameters contributes to the variance of output *Y* observed during repeated simulations. A parameter that is important for controlling *Y* will have a large contribution, and the parameter that is not important will have small or no contributions. The variance of output *Y* that depends on k input parameters *P* could be uniquely decomposed into the following terms:

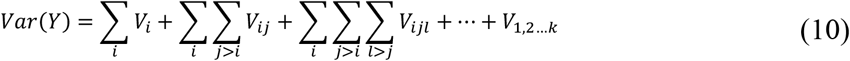

where

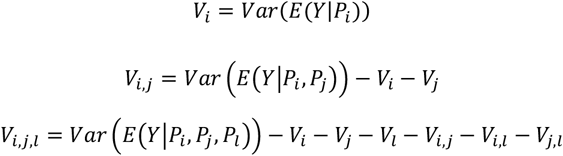

Two types of sensitivity indexes, *S*_*i*_ and 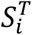, are useful in analyzing the contribution of input parameters. *S*_*i*_ is called sensitivity index, importance measure or first-order effect:

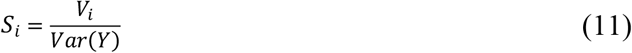

Another way to understand *S*_*i*_ is that it is a fraction of the variance of *Y* that would disappear if we fixed parameter *P*_*i*_ at some value. Parameter *P*_*i*_ would be considered important for controlling output Y if fixing its value would lead to a large reduction of the variance of Y (i.e., *S*_*i*_ is large). *S*_*i*_ has several nice properties where for an additive model ∑ *S*_*i*_ *= 1* and for any model with independent input parameters ∑ *S*_*i*_ *≤ 1*. For example, a large value of *1* − ∑ *S*_*i*_ indicates that interactions between parameters are important for controlling a specific output of the model. More generally, for any model with independent input parameters:

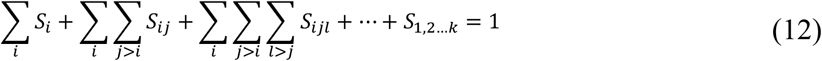

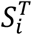 is called the total-order effect and contains all terms in ***Equation (12)*** that contain *i*:

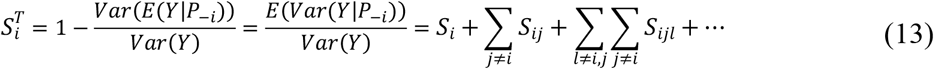

Another way to understand 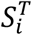 is that it is a fraction of the variance of *Y* that would be left if we fixed all parameters except parameter *P*_*i*_ at some value. Parameter *P*_*i*_ would be considered important for controlling output *Y* if fixing all the parameters except *P*_*i*_ at some value would still leave a large fraction of variance of *Y* (i.e., 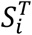 is large). Unlike *S*_*i*_, 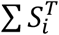 can be larger or smaller than 1 and ***Equation (12)*** does not hold for 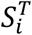 because 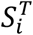 and 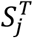 would have one or several overlapping terms (e.g., *S*_*ij*_).

We have calculated *S*_*i*_ and *S*^*T*^for various outputs of the model using the Sobol method of the GlobalSensitivity.jl package (***Dixit and Rackauckas, 2022***).

### Adoption of the Glycolysis Model for Simulation of [U-^13^C]-Glucose and [U-^13^C]-Lactate Tracing

Glycolysis model can be used to predict the results of heavy isotope tracing experiment. To predict tracing patterns, we decompose each rate equation into enzyme activity part and mass action part as described previously (***Hofmeyr, 1995; Noor et al., 2013; Reich and Selkov, 1981***):

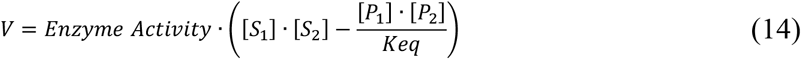

Enzyme activity depends on the total concentration of metabolites (i.e., sum of all isotopomers), while the mass action part is responsible for the propagation of the isotope label. Therefore, for modeling isotope tracing, we modify all kinetic equations to be able to use heavy and light isotope-labeled metabolites that we incorporate in the mass action part of all the equations, and we double the number of ODEs to account for time-dependent changes of both heavy and light isomers (i.e., we have one ODE for each isotope).

### Kinetic equations and numerical simulation of differential equations constituting the Two-Enzyme Model

We described the kinetics of Enzyme 1, Enzyme 2, and ATPase using simplified rate equations. We started from reversible Michaelis-Menten equations (***Noor et al., 2013***) for Enzyme 2 and ATPase. For Enzyme 1, we used the MWC model to add allosteric activation by ADP and phosphate and allosteric inhibition by ATP. We assumed that ADP and Phosphate only bind to active conformation, ATP only binds to inactive conformation, inactive MWC conformation of Enzyme 1 is catalytically inactive. The resulting complete rate equations for Enzyme 1, Enzyme 2 and ATPase are:

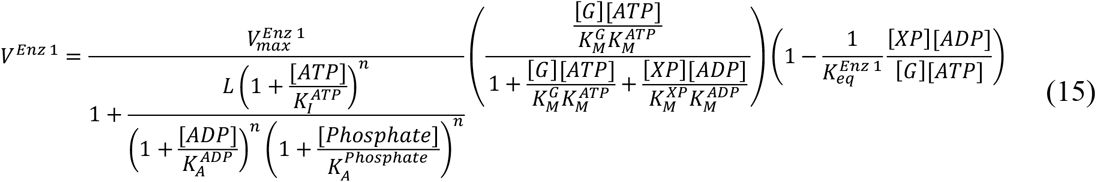

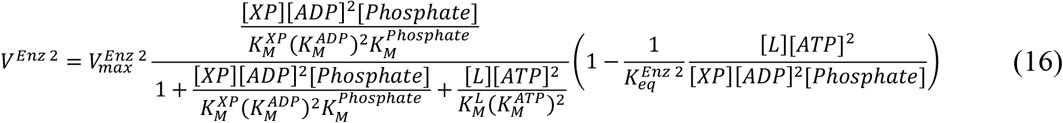

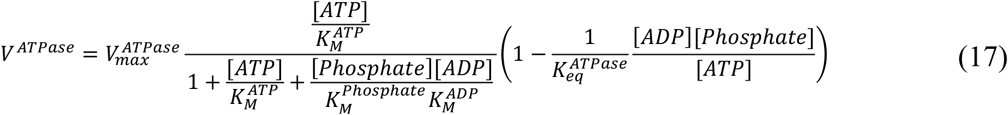

We then assumed that catalytic sites are saturated with substrates and regulatory sites are saturated with regulators simplifying the rate equations to ***Equations (2), (3), (4)*** reproduced here for convenience:

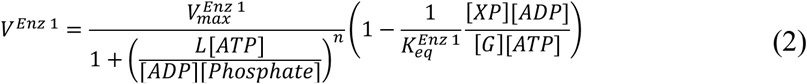

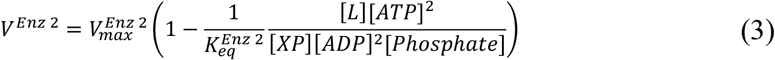

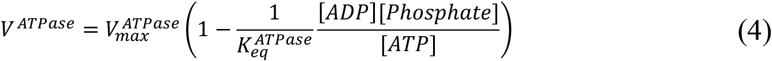

The two-enzyme model is a system of ordinary differential equations (ODEs). The number of equations is equal to the number of metabolites, and each equation consists of a derivative of metabolite concentration with respect to time being equal to the sum of enzyme rates producing the metabolite minus the sum of enzyme rates consuming the metabolite:

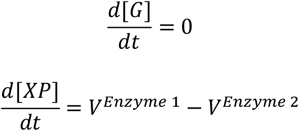

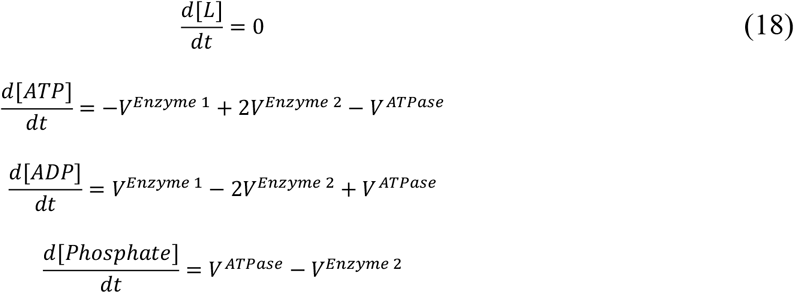

We numerically simulated ***Equations (6)*** using DifferentialEquations.jl library (***Rackauckas and Nie, 2017***) using Rodas5P solver for stiff differential equations with 10^−15^ absolute tolerance and 10^−8^ relative tolerance as for the full model described above.

## Key Resources Table

| REAGENT or RESOURCE | SOURCE | IDENTIFIER |
| --- | --- | --- |
| Experimental models: Cell lines |  |  |
| HeLa | ATCC |  |
| 293T | ATCC |  |
| C2C12 | ATCC |  |
| NIH3T3 | ATCC |  |
| Software and algorithms |  |  |
| Julia Programming Language (**Bezanson et al., 2015**) | https://julialang.org | N/A |
| DifferentialEquations.jl package for simulation of differential equations (**Rackauckas and Nie, 2017**) | https://github.com/SciML/DifferentialEquations.jl | N/A |
| GlobalSensitivity.jl package for global sensitivity analysis (**Dixit and Rackauckas, 2022**) | https://github.com/SciML/GlobalSensitivity.jl | N/A |
| Makie.jl package for plotting (**Danisch and Krumbiegel, 2021**) | https://github.com/MakieOrg/Makie.jl | N/A |

### Data and code availability

- All data reported in this paper is provided as supplementary files 1 and 2.
- All original code required to reproduce all the figures is publicly available at https://github.com/DenisTitovLab/Glycolysis.jl

## APPENDIX 1. Data-driven Identification of Kinetic Rate Equation for Glycolytic Enzymes

Appendix 1 contains descriptions of the derivation and fitting of enzyme rate equations used in the glycolysis model separated into three sections containing a description of the derivation of Monod-Wyman-Changeux kinetic rate equations (***Section 1***), an overview of the approach that we used for identifying the simplest equation for each enzyme (***Section 2***) and the largest section describing the derivation and fitting of kinetic rate equations for each glycolytic enzyme (***Section 3***).

### Section 1. Monod-Wyman-Changeux Model for Allosteric Enzymes

All the key regulatory enzymes in glycolysis (e.g., HK, PFK, GAPDH, PK) and other metabolic pathways are *allosterically regulated*, meaning that they are modulated at a region on the enzyme that is distinct from the active site, and oftentimes this modulation is mediated by metabolites that are not substrates or products of the enzyme. These regulators can change the Michaelis constant *K*_*M*_ or catalytic rate *k*_*cat*_ of the enzyme. The unusual characteristic of allosterically regulated enzymes is that they deviate from standard Michaelis-Menten-type hyperbolic kinetics by exhibiting positive and negative cooperativity. Several models have been proposed to describe the activity of such enzymes including Monod-Wyman-Changeux (MWC) (***Blangy et al., 1968; Monod et al., 1965b***) and Koshland-Nemethy-Filmer (KNF) (***Koshland et al., 1966b***). The MWC model postulates that allosteric enzymes can exist in two conformations, and whenever a metabolite binds to one subunit, it may change the global state of the enzyme, changing the state of all subunits in a concerted manner. On the other hand, the KNF model postulates that whenever substrates, products, or regulators bind to an enzyme’s subunit, they induce a conformational shift in that specific subunit that changes its affinity for binding additional metabolites. Mathematically, the MWC and KNF models are similar, but the KNF model typically requires a larger number of parameters to describe the behavior of an enzyme so we utilize the more parsimonious MWC model in this work and only use the induced fit idea from the KNF model to describe the interaction between ligands that are known to bind side by side to catalytic or regulatory sites.

In this section, we will derive the simplest form of the MWC equation for dimeric enzyme *E* with non-overlapping binding sites for substrate *S* and allosteric regulator *R* (***Appendix 1– figure 1A***). Each subunit of *E* is identical and has one site for *S* and one site for *R*. We will derive the MWC equation in stages starting with one subunit of *E* in isolation and then sequentially adding *S, R* and finally another subunit of *E*. Derivations of the MWC and KNF equations can be found in several excellent publications and books (***Marzen et al., 2013; Monod et al., 1965b; Phillips, 2020***).

The MWC model postulates that *E* can exist in two conformations. We will refer to the two conformations as active (*E*^*A*^) and inactive (*E*^*I*^). Let *L* be a constant that determines the ratio of *E*^*I*^ and *E*^*A*^. In the absence of *S* and *R*, the fraction of one subunit of *E* that is in the active and inactive conformation can be calculated using the following equations:

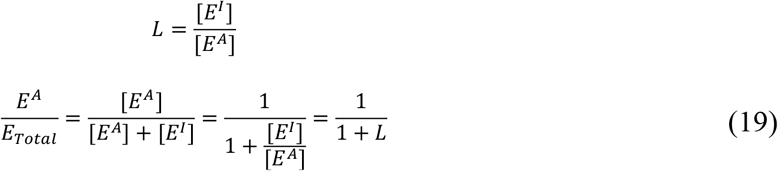

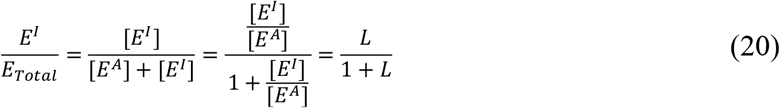

In presence of substrate *S*, we must add two more terms for *S* bound to *E*^*A*^ and *E*^*I*^. We describe the binding of *S* to *E*^*A*^ and *E*^*I*^ using the dissociation constants 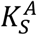 and 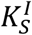. The fraction of *E* bound to *S* can be calculated using the following equations:

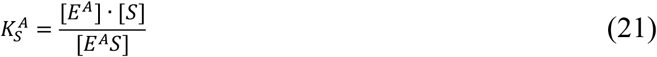

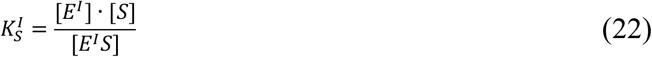

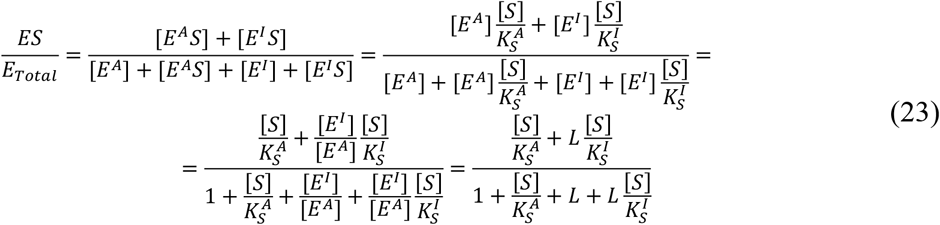

Using the equations above, the enzyme rate divided by the total concentration of enzyme can be approximated as:

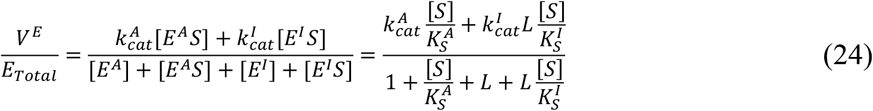

***Equation (24)*** holds in the context of rapid equilibrium or steady state. Rapid equilibrium assumes the binding and dissociation of *S* to *E* happen much faster than the catalytic reaction of *ES*. The steady state (or quasi steady state) approximation assumes that concentration of *S* changes much more slowly than concentrations of *E* and *ES*. The only difference is the physical meaning of 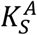 and 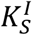 being either dissociation constants (rapid equilibrium) or Michaelis-Menten constants (steady state).

In presence of substrate *S* and allosteric regulator *R*, we add four more terms for *R* bound to *E*^*A*^, *E*^*I*^, *E*^*A*^*S*, and *E*^*I*^*S*. We describe binding of *R* to *E*^*A*^ and *E*^*I*^ using dissociation constants 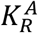 and 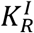, which we assume do not change if the substrate is bound or unbound. The fraction of *E* bound to *S* can be calculated using the following equations:

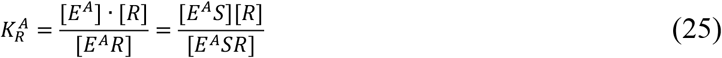

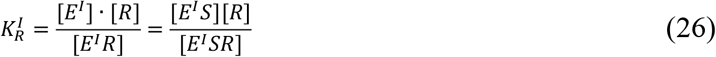

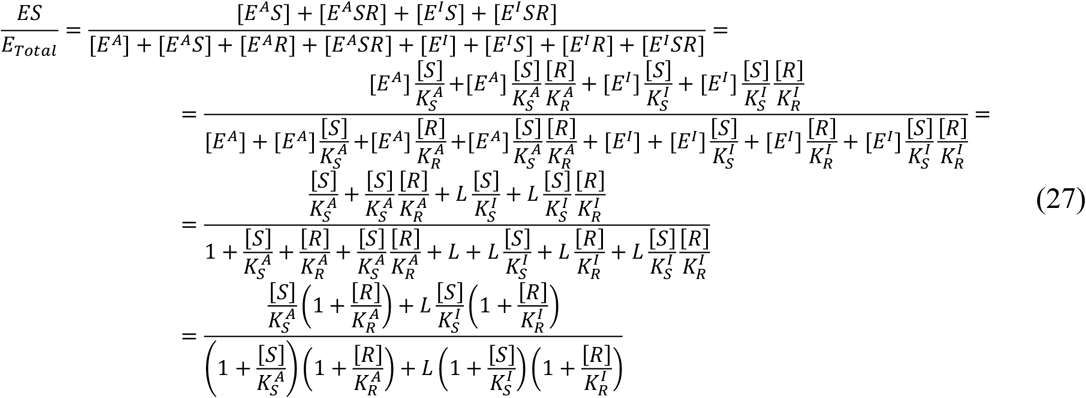

The enzyme rate naturally follows as:

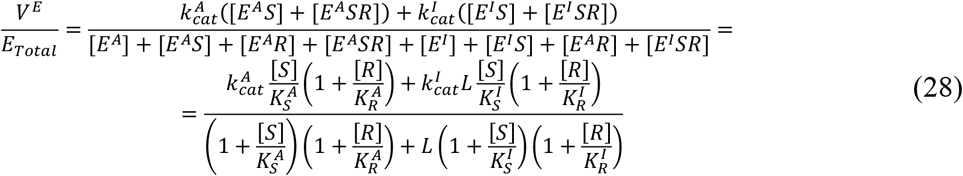

The equations so far have described a single subunit of *E*. If *E* is a dimer, we add additional terms describing binding of S and R to different subunits of *E*. The fraction of *E* bound to *S* is given by:

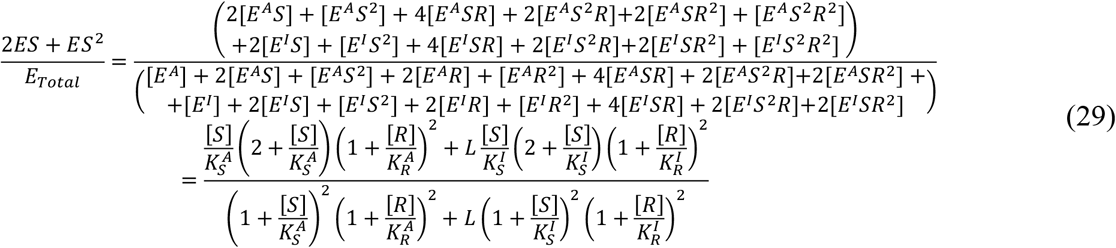

When we compute the enzyme rate, we assume that an enzyme with two bound substrates (*ES*^2^) catalyzes substrates twice as fast (*2k*_*cat*_) as an enzyme with one bound substrate, which enables us to factor out the number of subunits (here *n*=2) from every term in the numerator:

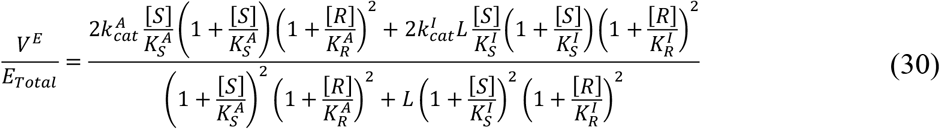

***Equation (30)*** presents a simple MWC description of allosteric regulation of a dimeric enzyme, and the [*S*]^*2*^ terms can give rise to a sigmoidal response that is found experimentally for all allosteric multimeric-enzymes in the glycolysis pathway. In ***Appendix 1–figure 1***, we show how the dependence of *V*^*E*^ from [*S*] becomes sigmoidal when S binds preferentially to *E*^*A*^ while *L* is large (i.e., *E*^*I*^ is the dominant conformation in the absence of *S*) and how the ratio of 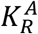 to 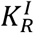 determines whether *R* is an activator or inhibitor of *E*.

**Appendix 1–figure 1.**
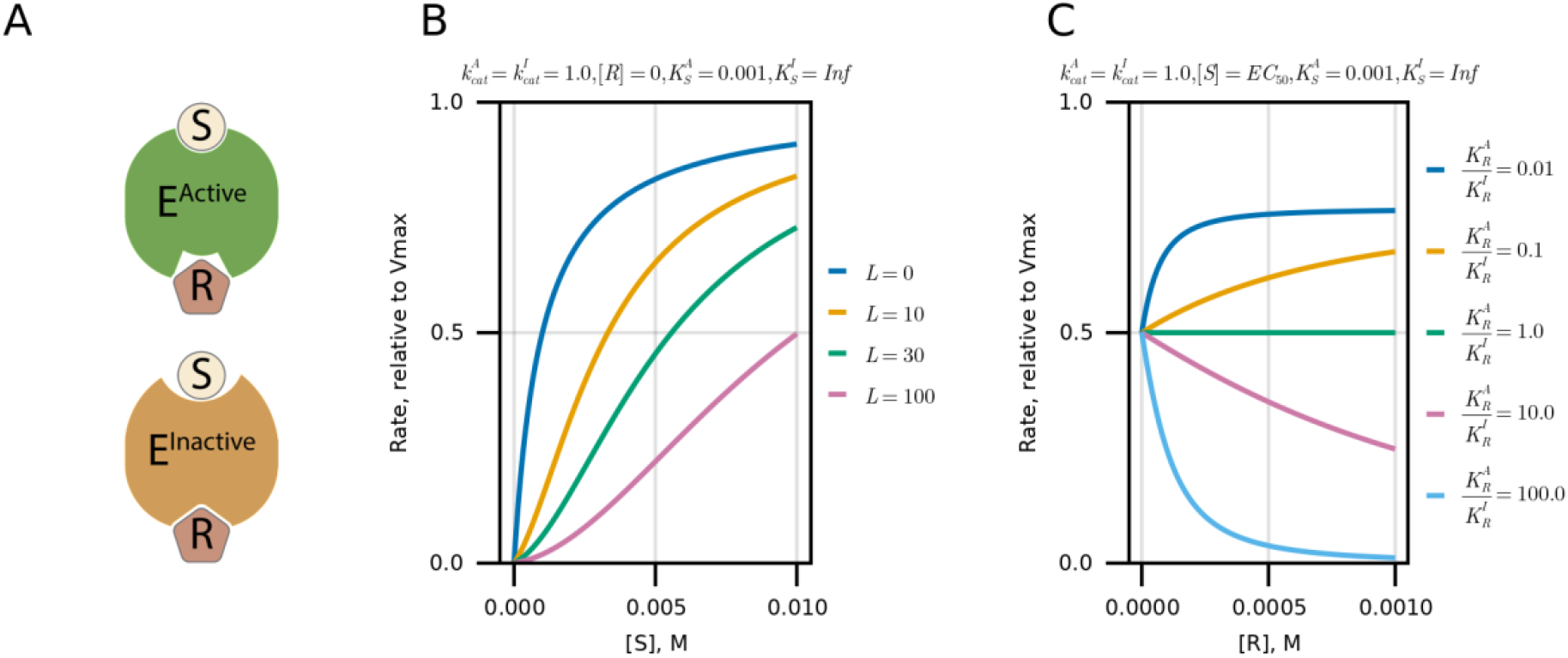
**A)** Schematic of one subunit of enzyme *E* that can exist in two conformations and can bind to substrate *S* and regulator *R*. **B)** Rate of *E* depending on the parameter *L*. **C)** Effect of regulator of rate of *E* depending on the ratio of binding constants of *R* to active and inactive conformations of *E*.

We next generalize the above equation to a reversible MWC enzyme that catalyzes a reaction between two substrates (*S*_*1*_ and *S*_*2*_) and two products (*P*_*1*_ and *P*_*2*_). MWC enzyme contains *n* identical subunits, where each subunit has one active site where substrates and products bind and *k* distinct sites for allosteric regulators (*R*_*i*_). We assume that both substrates must be bound for enzyme catalysis, that all *R*_*i*_ bind to distinct and non-overlapping sites, and that *S*_*1*_ and *P*_*1*_ as well as *S*_*2*_ and *P*_*2*_ bind in mutually exclusive manner. This equation is derived using the same approach as above and we omit the full derivation for conciseness:

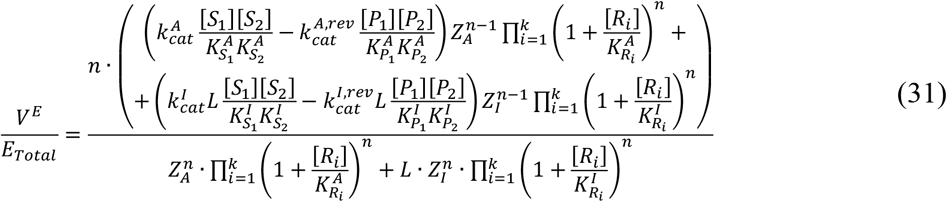

where

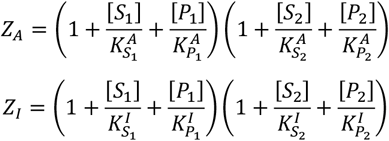

The Haldane relationship can be derived for this enzyme using an observation that the rates of all forward and reverse reactions are equal at equilibrium, and the ratio of product and substrate concentration is equal to equilibrium constant *K*_*eq*_. The principle of detailed balance states that at equilibrium, all individual steps of the process are at equilibrium. Therefore, for our MWC enzyme, the Haldane relationships will have the following form as forward and reverse reactions involving either Active or Inactive conformations must be equal at equilibrium:

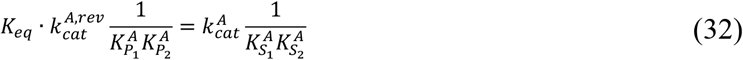

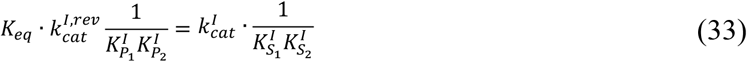

***Equation (31)*** can be rewritten using a Haldane relationship that connects kinetic and thermodynamic constants and by using specific activity 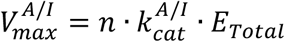:

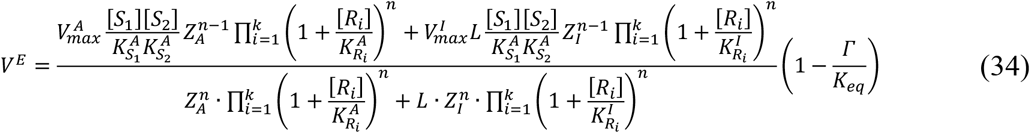

An even more general equation can include interaction terms β between substrates and products to describe enzymes where the binding of a substrate or product depends on whether another substrate or product is already bound:

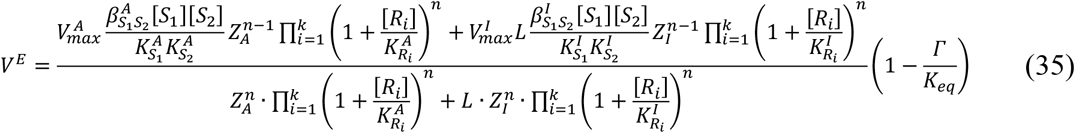

where

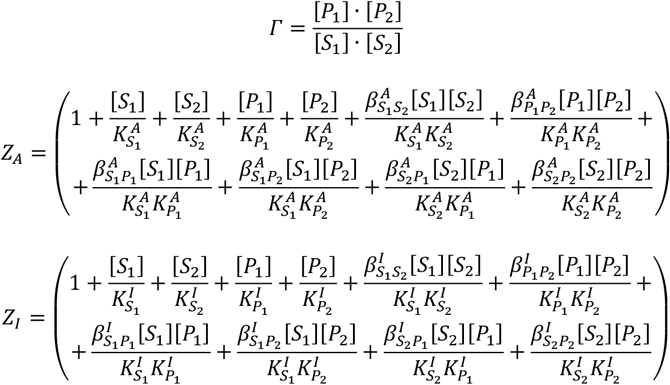

Note that ***Equation (31)*** represents the limit where 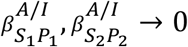and all other *β = 1*. The general form of ***Equation (35)*** could arise in biological contexts where substrates and products bind in close proximity such that their functional groups can interact to modify the binding affinity of each other. We only included interaction terms *β* for molecules that bind to overlapping or neighboring sites. It is also possible to include interaction terms for all molecules that bind to an enzyme as articulated by the classical KNF model, but it yields to a large number of parameters that are difficult to estimate with available data. We found that ***Equation (35)*** is sufficient to describe all the enzymes that we considered.

#### Section 2. Estimation of Kinetic Parameters for MWC Enzymes

In this section, we describe the general procedure to identify the kinetic rate equations and kinetic parameters that best describe the data for MWC enzymes HK1, PFKP, GAPDH, and PKM2. The detailed results of the fitting for each enzyme are described in ***Section 3***. ***Rate Equations for Enzymes in the Mammalian Glycolysis Pathway***. The fitting procedure is subdivided into two stages that are summarized in this paragraph and described in more detail throughout the rest of this section. First, we used cross-validation and regularization to identify the kinetic equation that can adequately describe the data using the smallest number of kinetic parameters. This step is important as using an equation with a large number of parameters (e.g., general MWC ***Equation (35)***) can result in overfitting, which will decrease the predictive accuracy of the model as overfit equations will not be able to accurately predict enzyme activity under conditions not encountered in the fitting data. Second, we used bootstrapping to estimate the values and confidence intervals of kinetic parameters in the resulting kinetic equation. Bootstrapping allowed us to confirm that we can estimate unique values for all kinetic parameters (i.e., there is no sloppiness or parameter non-identifiability).

We obtained data for enzyme fitting by manually digitizing figures from previous publications using the WebPlotDigitizer tool (***Rohatgi, 2022***). For each enzyme, we obtained 400-800 data points consisting of paired enzyme rates and corresponding metabolite concentrations. Some of the data were inconsistent because we combined figures from different publications generated by different groups under different conditions. To remove outlier figures, we fit a kinetic equation to the dataset missing one figure at a time and used a Bonferroni corrected t-test p values to identify figures that led to a statistically significant improvement in the goodness of fit with a conservative cutoff of p < 10^−4^. This procedure removed < 10% of the figures from the dataset for each enzyme and minimally affected the inferred kinetic parameters, but it made cross-validation and regularization used to identify the final kinetic rate equation more robust. Many of the outlier figures had points with enzyme rate values close to 0, which are difficult to accurately estimate from published plots due to limited resolution. For most outlier figures, we only had to remove such points that are close to 0 instead of removing the whole outlier figure.

We used data fitting to estimate kinetic constants for enzyme rate equation (e.g., general MWC ***Equation (35)***) from data that consisted of paired values of enzyme rate V and concentrations of substrates [S], products [P] and regulators [R]. Fitting consisted of minimizing the least squares loss function:

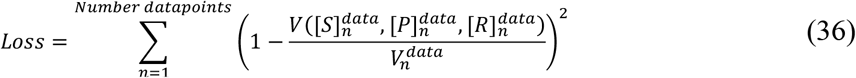

We used the global optimization algorithm Covariance-Matrix Adaptation Evolution Strategy (CMA-ES) to minimize the loss function. We encoded kinetic parameters by converting them to log scale and rescaled all parameters to lie within the interval [0, 10]. We converted the binding constants *K*, specific activity *V*_*max*_, and ratio of inactive to active conformations of MWC enzyme *L* from [10^−10^, 10^3^], [10^−3^, 1], and [10^−5^, 10^5^] in linear scale into the [0,10] in log scale using 10^−10^×10^13*x*/10^, 10^−3^×10^3*x*/10^, and 10^−5^×10^10*x*/10^ equations, respectively. Parameter encoding is essential for the success rate of the fitting procedure. We rerun the CMA-ES algorithm twenty times and took the result with the smallest Loss value to increase the chances that we find the global minimum. Rerunning our fitting procedure on the same data produced the same values of *Loss* function, which demonstrates the robustness of our data fitting procedure. Estimation of kinetic parameters using the latter method requires 30 seconds to 20 minutes to complete on a single core of Apple M1 Max processor depending on the number of parameters in the MWC equation. We allowed each figure to have its own *V*_*max*_ values to account for the fact that different publications and even different figures within the same publication use enzymes with different *V*_*max*_ due to variability in enzyme purity and many publications use arbitrary units to report enzyme rates. We normalized the rates in each figure to 1 by dividing all the rates by the maximal rate in respective figure and allowed *V*_*max*_ for each figure to be in the range of [0.25, 16] using figure specific weight parameters. We converted the weight parameters from [0.25, 16] in linear scale into the [0,10] in log scale using 2^−2^×2^6*x*/10^ equations. Despite the relatively large range of allowed *V*_*max*_ values, we often had to adjust the arbitrary *V*_*max*_ scales of several figures until the *V*_*max*_ values for each figure fell within the *V*_*max*_ range supplied to the algorithm without hitting the boundaries. The latter was usually required for figures that reported rates in the reverse direction or rates in the presence of inhibitors, where the maximal rate in the figure can be orders of magnitude away from *V*_*max*_ in the forward direction. The actual *V*_*max*_ value that was used in the model was calculated separately from data fitting by averaging specific activity for a particular enzyme from several publications.

We used a combination of cross-validation and regularization to identify the minimal equation that can accurately describe the available data. The number of parameters in general MWC ***Equation (35)*** can be decreased if a metabolite binds with the same affinity to active and inactive conformations (i.e., *K*_*A*_ *= K*_*I*_), if a metabolite only binds to active or inactive conformations (i.e., *K*_*A*_ *=* ∞ or *K*_*I*_ *=* ∞), or if a metabolite X binding to an enzyme is not affected by metabolite Y binding (i.e., 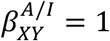). To choose between two different equations, we used 10-fold cross-validation that consisted of dividing the figures in our dataset into 10 parts at random and then fitting the equation using 9/10 of the data and testing the fit on the remaining 1/10. The procedure is repeated for each 1/10 of the data and we repeated the whole cross-validation 100 times for each equation. Cross-validation allows us to control against overfitting and ensure that we select a kinetic rate equation that is best capable of predicting enzyme activity under novel conditions not encountered during data fitting. Because it would be computationally too expensive to compare every variant of the general MWC ***Equation (35)*** with each other, we use regularization to remove many parameters at once. Regularization was performed by incorporating additional parameters into the Loss function (36). We used *min(1*, |*log(K*_*Active*_/ *K*_*Inactive*_ )|) or *min(1*, |*log(β*_*Active*_ /*β*_*Inactive*_ )|) to identify metabolite that bind with the same affinity to active and inactive conformation, and *min(1*, |*log(1000*/*K)*|) or |*β*| to identify metabolites that do not bind to a particular conformation. As we do not know a priory which metabolites bind with the same affinity to active and inactive conformation or which metabolites bind only to one conformation, we performed 4 sequential rounds of regularization. First, we found metabolites that bind with the same affinity to both conformations and set the corresponding *K*_*Active*_ *= K*_*Inactive*_ or *β*_*Active*_ *= β*_*Inactive*_. Second, for all *β*_*Active*_ *= β*_*Inactive*_ we found β for metabolite pairs where data doesn’t support their simultaneous binding and remove them by making relevant *β = 0*. Third, we find all β that are equal to 1. Finally, we found all *K* where metabolites only bind to one conformation. This procedure allowed us to drop most of the kinetics parameters in the general MWC ***Equation (35)*** while keeping the test Loss value of cross-validation unchanged or even decreased. The latter is a classic example of a bias-variance tradeoff observed in statistical learning, which suggests that the MWC ***Equation (35)*** is overfitting our data. It is likely that additional parameters could be added to our kinetic rate equations when more data becomes available than the 400-800 data points per enzyme that we had. Cross-validation and regularization is a computationally intensive procedure (1000s of CPU-hours per enzyme) as it requires hundreds of parameter estimations performed under different values of the regularization parameter. We used the Savio computational cluster at the University of California Berkeley to perform cross-validation and regularization.

After selecting the kinetic rate equation, we used bootstrapping to estimate the values of kinetic constants and their confidence intervals. Here, we repeatedly fitted our kinetic equation to data that is generated by sampling the original data with substitution so that we have the same number of data points, but some are repeated, and others are missing at random. The distribution of kinetic parameters estimated from 10,000 fits to bootstrapped data was then used to confirm that our fitting procedure estimated the unique values of kinetic parameters and to calculate the mean and standard deviation of kinetic parameters that are used in the ODE model of glycolysis.

#### Section 3. Rate Equations for Enzymes in the Mammalian Glycolysis Pathway

Here, we describe the estimation of kinetic parameters for all enzyme used in the glycolysis model including extensive discussion of known regulation of each enzyme.

#### Glucose transporter (GLUT)

*GLUT Reaction:* Glucose_*media*_ ⇄ Glucose_*cell*_

*GLUT genes:* Humans have four genes coding for GLUT isoforms SLC2A1-4 (also referred to as GLUT1-4). We focus on GLUT1 and GLUT3 isoforms as it they are the most abundant GLUT isoform in proliferating cells based on proteomics data (***Supplementary File 1***).

*GLUT kinetic rate equation:* We used the reversible Michaelis-Menten equation to describe the activity of GLUT:

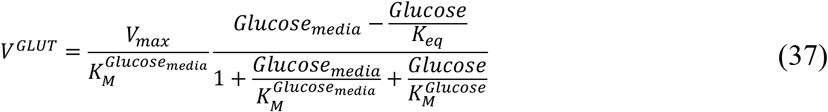

*GLUT protein bound metabolite equations:*

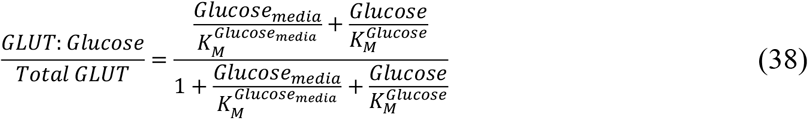

**Appendix 1–Table 1.** GLUT kinetic Parameters. Kinetic parameters were used as is from references below without fitting the equation to data. *V*_*max*_ values for some publications were adjusted from 20C to 37C by multiplying by 1.58 based on the temperature dependence (***Lowe and Walmsley, 1986***) of *V*_*max*_ and some of the *K*_*M*_ values are for 3-O-methylglucose which is believed to have similar *K*_*M*_ to glucose.

| Parameter | Mean ± SEM | Values | References |
| --- | --- | --- | --- |
| Molecular Weight, g/mol | 54,084 |  | UniProt: P11166 |
| $K_M^{Glucose_{media}} = K_M^{Glucose_{cell}}, mM$ | $20 \pm 4$ | 13.95, 17, 19.9, 26.2, 20, 20, 20 | Lowe et al. BBA 857 (1986) 146-154;<br>Gould et al. Biochemistry 1991, 30, 5139-5145;<br>Palfreyman et al. Biochem. J. (1992) 284, 275-281;<br>Nishimura et al. JBC Vol.268, No. 12, Issue of April 25, pp. 8514-8520, 1993;<br>Gould et al. Biochemistry 1989, 28, 9447-9452;<br>Keller et al. JBC Vol. 264, No. 32, Issue of November15, pp. 18884-18889, 1989;<br>Holman et al BBA, 649 (1981) 503-514; |
| $V_{max}, \frac{\mu mole}{min \cdot mg}$ | $1100 \pm 300$ | 977, 1333, 585, 293, 2444, 1258 | Lowe et al BBA 857 (1986) 146-154;<br>Palfreyman et al. Biochem. J. (1992) 284, 275-281;<br>Nishimura et al. JBC Vol.268, No. 12, Issue of April 25, pp. 8514-8520, 1993;<br>Gould et al. Biochemistry 1989, 28, 9447-9452;<br>Keller et al. JBC Vol. 264, No. 32, Issue of November15, pp. 18884-18889, 1989;<br>Holman et al BBA, 649 (1981) 503-514; |
| $K_{eq}$ | $1$ | *Postulated to be 1 since transport is diffusion driven* | |

*Reported GLUT regulation that was not included in our model:* GLUT like many other transporters can exist in two conformations with glucose binding sites facing the media or facing the cytosol. The kinetic rate equation incorporating the two conformations will include another term in the denominator containing *Glucose*_*media*_ ⋅ *Glucose/K*. We did not include this additional term as we could not find data reporting the value of *K*. Addition of this term will slow down the GLUT rate, but this will not affect our predictions appreciably as GLUT is not the rate-limiting enzyme in our model. Future extension of our models can incorporate this term to improve the accuracy of predictions.

#### Hexokinase (HK)

Human hexokinase (HK) catalyzes phosphorylation of glucose by ATP to produce glucose 6-phosphate (G6P) and ADP.

*HK Reaction:* Glucose + ATP ⇄ G6P + ADP

*HK genes:* Humans have 4 genes (HK1, HK2, HK3, HK4) coding for HK that have tissues-specific expression and distinct allosteric regulation (***Wilson, 2003***). We focus on HK1 as it is the most abundant HK isoform in proliferating cells based on proteomics data (***Supplementary File 1***).

*Regulation of HK by small molecules:* HK1-3 are inhibited by product G6P. Inhibition by G6P is competitive in relation to ATP and uncompetitive in relation to glucose (***Copley and Fromm, 1967***). Inhibition of HK1 by G6P, but not HK2-3, is relieved by inorganic phosphate (***Rose et al., 1964***). In the absence of G6P, physiological concentrations of phosphate (i.e., <10 mM) have no effect on HK1 activity (***Rijksen and Staal, 1977***). G6P induces dimerization of HK1 that is reversible by phosphate and ATP suggesting that G6P binding induces conformational changes consistent with allosteric regulation (***Chakrabarti and Kenkare, 1974; Wilson, 1973***).

*Structure of HK and number of binding sites for small molecules*: To formulate the general MWC equation for HK1, we need to know the oligomeric state of HK1, number of binding sites for each metabolite, and whether any metabolites bind to the same site. Biochemical studies have shown that HK1-3 are monomeric in solution (***Redkar and Kenkare, 1972***). The 100 kDa mammalian HK1-3 contain two putative catalytic domains that appear to be the result of a fusion of two ancestral 50 kDa hexokinases. HK1 and HK3 have catalytically inactive N-terminal and catalytically active C-terminal domains while HK2 retained catalytic activity in both domains (***Ardehali et al., 1999***).

Structural, kinetic, and mutagenesis studies have shown that HK1 has three binding sites for G6P (***Fang et al., 1998; Liu et al., 1999***). One G6P site represents a catalytic site in the C-terminal domain (overlaps with glucose), one inhibitory site for G6P overlaps with ATP catalytic site (i.e., competitive inhibition site) in the C-terminal domain (glucose and G6P can binds side by side), and another inhibitory site (i.e., allosteric site) for G6P overlaps with the phosphate binding site in the N-terminal domain (***Appendix 1–figure 2C***). Mutation of the N-terminal G6P binding site eliminates regulation of HK1 by phosphate and mutation of C-terminal site removes inhibition of truncated C-terminal catalytic domain of HK1 by AnG6P (G6P analog) (***Fang et al., 1998***). However, mutation of either N-terminal or C-terminal G6P sites alone has no effect on G6P inhibition and mutation of both sites is required to abolish regulation of HK1 by G6P (***Liu et al., 1999***). Despite the presence of multiple binding sites of G6P, binding studies showed that only one G6P is bound to enzyme at a time suggestion that binding of G6P to allosteric and competitive inhibitory sites happens in a mutually exclusive manner (***Chou and Wilson, 1974; Mehta et al., 1988***).

Structural and biochemical studies have also shown that HK1 has two sites for ATP and ADP (***Aleshin et al., 2000; Rosano et al., 1999***). One site is the catalytic site in the C-terminal domain and the second site is in the N-terminal domain. The existence of the N-terminal binding site for ATP and ADP has been confirmed with structural (***Aleshin et al., 2000; Rosano et al., 1999***) data but its role in the regulation of HK1 activity is unknown as no mutagenesis experiments have been performed to our knowledge.

Finally, HK1 has two binding sites for glucose, one in the active site in the C-terminal domain and another in the ancestral active site in the N-terminal domain. In both sites, glucose has been observed to bind side by side with G6P.

*General MWC rate equation and Haldane relationship for HK1:* Based on the structural data described above we formulated the following rate equation based on the assumption that HK1 exists in active state (*A*) with a catalytic rates 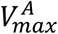 and 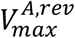 and an inactive state (*I*) with catalytic rates 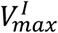 and 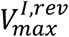. In the absence of all substrates and regulators (i.e., Glucose, ATP, ADP, *P*_*i*_, G6P), the ratio between the inactive and active states is *L*. Each substrate, product, and regulator can have a different binding constant *K* for the active and inactive enzyme state. We calculated the 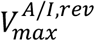 from the other kinetic constants using the Haldane relationship as the *V*_*max*_ in the reverse direction is rarely measured in the literature. Altogether, structural and regulatory information about HK1 translates into the following general MWC equation for HK1 (note that the simplified kinetic ***Equation (40)*** was used in the model as described below):

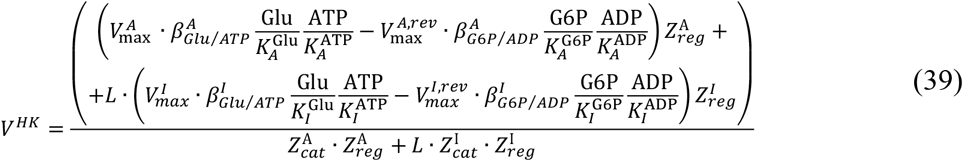

where

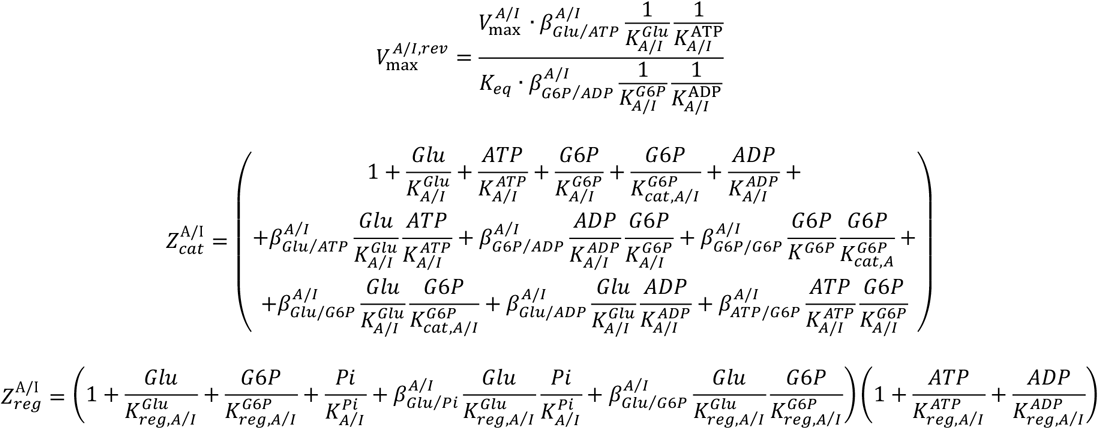

*Data used for HK1:* We manually digitized ∼500 data points from eight publications (***Bianchi et al., 1998; Copley and Fromm, 1967; Ellison et al., 1975, 1974; Kosow et al., 1973; Ning et al., 1969; Solheim and Fromm, 1983; Ureta et al., 1985***) describing the rate of purified HK1 enzyme in the presence of different concentrations of Glucose, ATP, G6P, ADP, and inorganic phosphate (*P*_*i*_).

*Identification of minimal HK1 kinetic rate equation:* ***Equation (39)*** describes all possible kinetic parameters for HK1 based on structural and biochemical data. Some of these parameters might not be required to describe HK1 activity (e.g., *K*_*A*_ *= K*_*I*_, *β*_*A*_ *= β*_*I*_, *β = 0, β = 1*) and their inclusion might lead to a poor prediction of enzyme rate under conditions not encountered in the fitting data due to overfitting. Therefore, we used regularization and cross-validation to determine if HK1 equation with a smaller number of kinetic parameters can achieve a lower *Loss* test value of cross-validation. The Loss values were calculated using ***Equations (36), (39)***. We performed four sequential rounds of regularization, as described in ***Section 2***. ***Estimation of Kinetic Parameters for MWC Enzymes***. Improvement of cross-validation *Loss* test value after the removal of parameters after each cross-validation cycle is shown in ***Appendix 1–figure 2B***. During the first regularization, we added terms 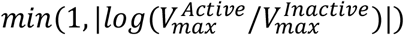, *min(1*, |*log(K*_*Active*_/*K*_*Inactive*_)|) and *min(1*, |*log(β*_*Active*_ /*β*_*Inactive*_)|) to the *Loss* function to identify metabolites that bind with the same affinity to active and inactive conformation. We were able to set *β*_*A*_ *= β*_*I*_ for all *β* and *K*_*A*_ *= K*_*I*_ for 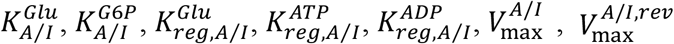 (note that this meant we could remove terms 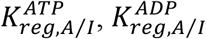 altogether as they cancel out in ***Equation (39)*** when *K*_*A*_ *= K*_*I*_), which decreased the number of parameters in ***Equation (39)*** from 41 to 21 while decreasing the cross-validation test score (***Appendix 1–figure 2B***). During the second and third rounds of regularizations, we added |*β*| and then *min(1*, |*log(β)*|) to the *Loss* function to identify *β =* 0 and *β = 1*, respectively. We were able to set *β =* 0 for 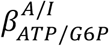 and *β = 1* for 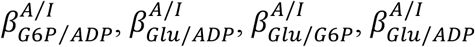, decreasing the number of parameters to 16 while further decreasing the cross-validation test score. During fourth round of cross-validation, we added terms *min(1*, |*log(1000*/*K)*|) and *min(1*, |*log(1*/*β)*|) again to identify *K =* ∞ and *β = 1*. We were able to set *K =* ∞ for 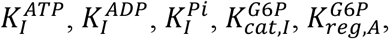,decreasing the number of parameters to 11 without an effect on the cross-validation test score. As final steps, we (i) absorbed parameter *L* into 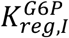 because binding of G6P to the regulatory site was the only site where a metabolite bound preferentially to inactive conformation making the inclusion of *L* and 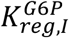 largely redundant and (ii) substituted 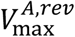 for *K*_*eq*_ by combining kinetic rate equation with Haldane relationship and (iii) set 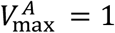 as this value is determined separately from the specific activity of the enzyme. The final simplified kinetic rate equation for HK1 had 9 parameters (including 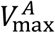) instead of 41 and about 2-fold lower *Loss* test value for cross-validation (***Appendix 1–figure 2***B) compared to ***Equation (39)***:

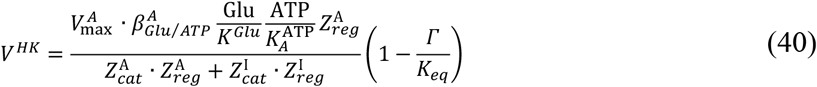

where

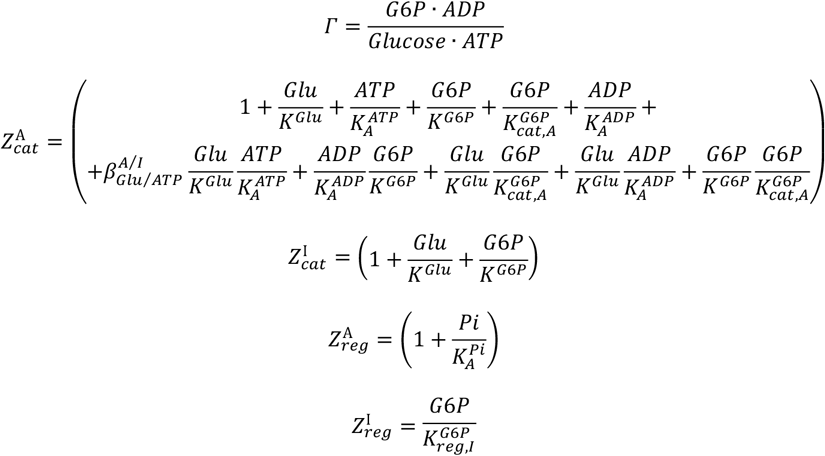

***Equation (40)*** was used to describe HK1 activity in our model. The least squares *Loss* function values that we obtained for the cross-validation train and test fits of the ***Equation (40)*** are 0.026 and 0.032, corresponding to an average error of 16% and 18% per data point, which is within the typical variability of *in vitro* kinetic experiments.

*HK1 protein bound metabolite equations:*

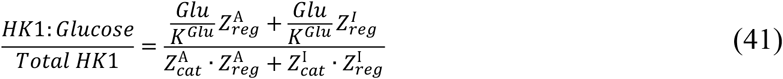

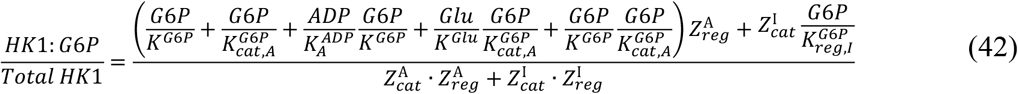

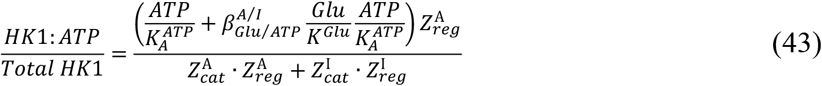

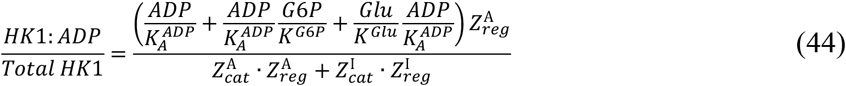

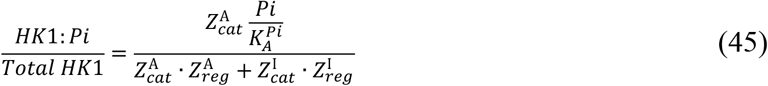

where

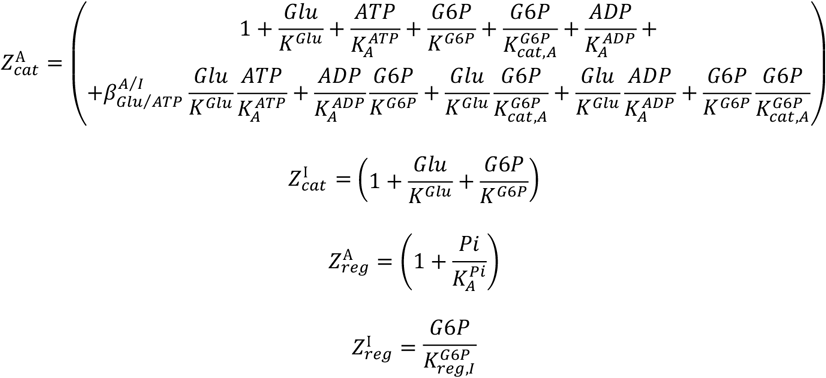

*Estimation of HK1 kinetic constants from data:* We used 10,000 fits to bootstrapped datasets to estimate kinetic parameters for ***Equation (40)*** and their confidence intervals. ***Appendix 1–figure 2C*** shows that we obtain narrow confidence interval for each of the parameters that shows that our data is sufficient to uniquely estimate all parameters of ***Equation (40)***. The fits of the kinetic equation to all data are shown in ***Appendix 1–figure 2D***.

*Discussion of the results of HK1 kinetic constants estimation:* ***Equation (40)*** describes several properties of HK1 that were not explicitly included in the data. First, ***Equation (40)*** shows that the binding of G6P to catalytic inhibitory site and regulatory site are mutually exclusive, which is supported by direct binding studies that demonstrate that HK1 has only one high-affinity site for G6P (***Chou and Wilson, 1974; Mehta et al., 1988***). Second, ***Equation (40)*** shows that mutation of either catalytic inhibitory site or regulatory site alone will not abrogate the ability of G6P to inhibit HK1 and mutation of both sites is necessary, which is supported by biochemical data (***Liu et al., 1999***). The ability of ***Equation (40)*** to predict regulation of HK1 not encountered in fitting data provides additional evidence that our fitting procedure produces accurate description of HK1 activity.

*Reported HK1 regulation that was not included in our model:* We are not aware of any regulators of HK1 that we did not consider.

**Appendix 1–figure 2.**
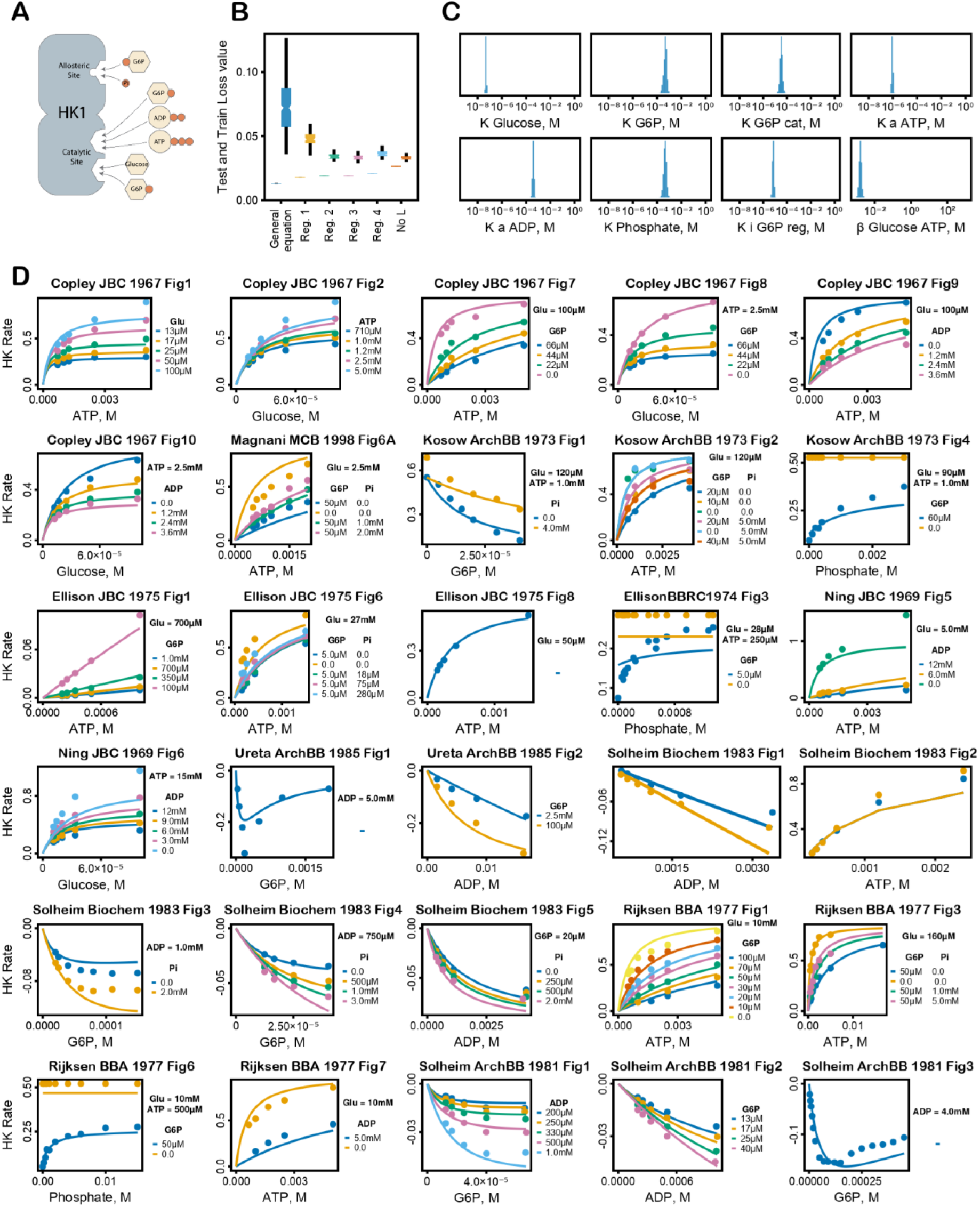
HK1 kinetic equation fitting. **(A)** Schematic of HK1 regulation, **(B)** Change of HK1 cross-validation fit (left dodge) and test (right dodge) *Loss* values after sequential rounds of regularization. ***Equation (39)*** corresponds to left most values and equation (40) corresponds to right most values, **(C)** Histogram of estimated kinetic parameters of equation (40) from fitting 10,000 bootstrapped datasets, **(D)** Equation (40) with final kinetic parameters plotted on top of all of the data used for fitting.

**Appendix 1–Table 2.**
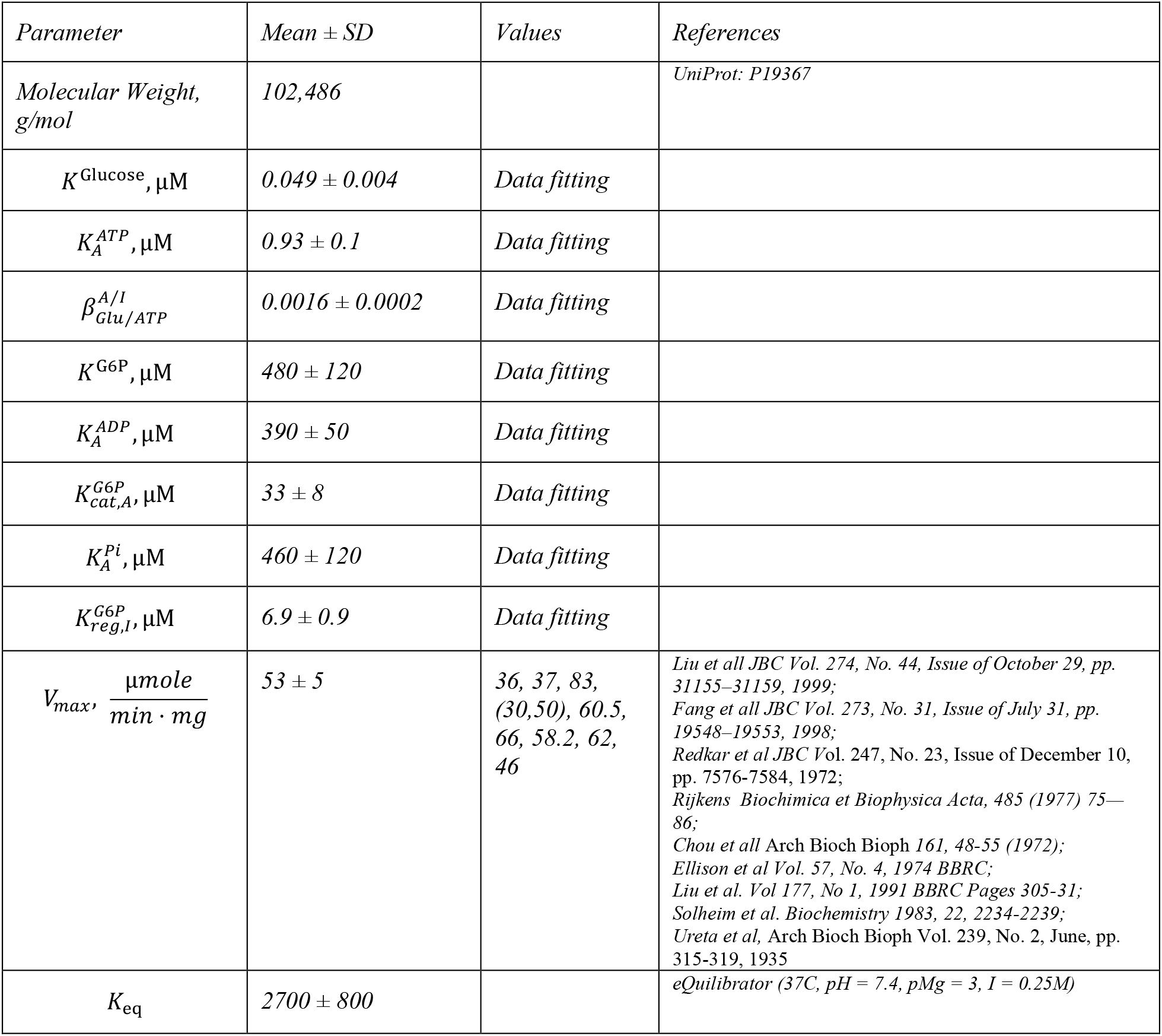
HK1 Kinetic Parameters. Mean ± standard deviation of parameter bootstrap values estimated by fitting of ***Equation (40)*** to data as described above.

| *Parameter* | *Mean ± SD* | *Values* | *References* |
| --- | --- | --- | --- |
| *Molecular Weight, g/mol* | 102,486 |  | *UniProt: P19367* |
| $K^{\text{Glucose}}, \mu\text{M}$ | $0.049 \pm 0.004$ | *Data fitting* | |
| $K_A^{\text{ATP}}, \mu\text{M}$ | $0.93 \pm 0.1$ | *Data fitting* | |
| $\beta_{\text{Glu/ATP}}^{A/I}$ | $0.0016 \pm 0.0002$ | *Data fitting* | |
| $K^{\text{G6P}}, \mu\text{M}$ | $480 \pm 120$ | *Data fitting* | |
| $K_A^{\text{ADP}}, \mu\text{M}$ | $390 \pm 50$ | *Data fitting* | |
| $K_{\text{cat},A}^{\text{G6P}}, \mu\text{M}$ | $33 \pm 8$ | *Data fitting* | |
| $K_A^{\text{Pi}}, \mu\text{M}$ | $460 \pm 120$ | *Data fitting* | |
| $K_{\text{reg},I}^{\text{G6P}}, \mu\text{M}$ | $6.9 \pm 0.9$ | *Data fitting* | |
| $V_{\text{max}}, \frac{\mu\text{mole}}{\text{min} \cdot \text{mg}}$ | $53 \pm 5$ | 36, 37, 83,<br>(30,50), 60.5,<br>66, 58.2, 62,<br>46 | *Liu et al JBC Vol. 274, No. 44, Issue of October 29, pp. 31155–31159, 1999;*<br>*Fang et al JBC Vol. 273, No. 31, Issue of July 31, pp. 19548–19553, 1998;*<br>*Redkar et al JBC Vol. 247, No. 23, Issue of December 10, pp. 7576–7584, 1972;*<br>*Rijkens Biochimica et Biophysica Acta, 485 (1977) 75–86;*<br>*Chou et al Arch Bioch Bioph 161, 48-55 (1972);*<br>*Ellison et al Vol. 57, No. 4, 1974 BBRC;*<br>*Liu et al. Vol 177, No 1, 1991 BBRC Pages 305-31;*<br>*Solheim et al. Biochemistry 1983, 22, 2234-2239;*<br>*Ureta et al, Arch Bioch Bioph Vol. 239, No. 2, June, pp. 315-319, 1935* |
| $K_{\text{eq}}$ | $2700 \pm 800$ | | *eQuilibrator (37C, pH = 7.4, pMg = 3, I = 0.25M)* |

#### Glucose-6-phosphate isomerase (GPI)

*GPI Reaction:* G6P ⇄ F6P

*GPI genes:* Humans have one gene coding for GPI

*GPI kinetic rate equation:* We used reversible Michaelis-Menten equation to describe activity of GPI:

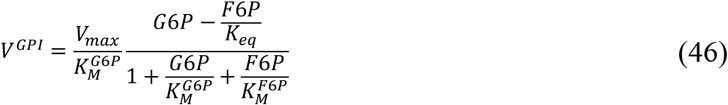

*GPI protein bound metabolite equations:*

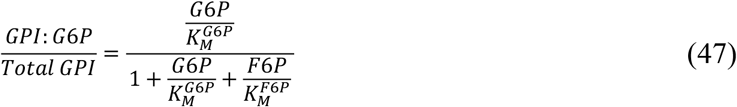

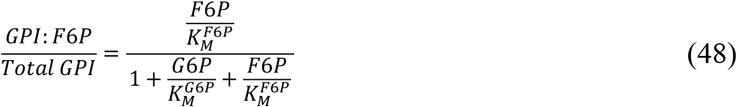

*GPI kinetic parameters:* Kinetic parameters were used as is from the references below without fitting the equation to data. More articles reported *V*_*max*_ in the reverse reaction as it is easier to measure the reverse reaction spectrophotometrically by coupling G6P production to NADPH production. We used specific activity in the reverse direction when specific activity in the forward direction was not available as measurements of both activities simultaneously show that they are close to each other (***Lin et al., 2009***).

*Reported GPI regulation that was not included in our model:* Erythrose-4-phosphate and 6-phosphogluconate are potent competitive inhibitors of GPI with *K*_*i*_ *in the range of* 1-10 µM. We have not included these regulators in our model because erythrose-4-phosphate and 6-phosphogluconate are products of the pentose phosphate pathway and are not produced by glycolytic enzymes. Future extension of the model to the pentose phosphate pathway will allow us to include these regulators.

**Appendix 1–Table 3.** Kinetic parameters for GPI.

| Parameter | Mean ± SEM | Values | References |
| --- | --- | --- | --- |
| Molecular Weight, g/mol | 63,147 |  | UniProt: P06744 |
| $K_M^{G6P}$, μM | $330 \pm 100$ | 445, 110, 660, 310, 120 | Kanno et al. *Blood Cells, Molecules, and Diseases* (1998) 24(4) Feb 28: 54–61;<br>Kahana et al. *Volume 235, Issue 8, 1 August 1960, Pages 2178-2184*;<br>Arnold et al. *Pediatric Research* volume 8, pages 26–30 (1974);<br>JED Dyson, EA Noltmann. *JBC*, 243, ISSUE 7, P1401-1414, APRIL 1968;<br>Hines, Wolfe, *Biochemistry*. Vol. 2, No. 4, July Aug., 1963 |
| $K_M^{F6P}$, μM | $70 \pm 20$ | 37, 63, 19, 32, 97, 174, 70 | H.-Y. Lin et al. / *Biochimica et Biophysica Acta* 1794 (2009) 315–323 321;<br>Kanno et al. *Blood Cells, Molecules, and Diseases* (1998) 24(4) Feb 28: 54–61;<br>Tsuboi et al. *JBC* Vol. 246, No. 24, Issue of December 25, pp. 7586- |
|  |  |  | 7594, 1971;<br>Kahana et al. Volume 235, Issue 8, 1 August 1960, Pages 2178-2184;<br>Arnold et al. *Pediatric Research* volume 8, pages 26–30 (1974);<br>JED Dyson, EA Noltmann. *JBC*, 243, ISSUE 7, P1401-1414, APRIL 1968;<br>Hines, Wolfe, *Biochemistry*. Vol. 2, No. 4, July Aug., 1963 |
| $V_{max}, \frac{\mu mole}{min \cdot mg}$ | $790 \pm 190$ | 951, 400, 1600,<br>820, 586, 402 | H.-Y. Lin et al. / *Biochimica et Biophysica Acta* 1794 (2009) 315–323 321;<br>Kanno et al. *Blood Cells, Molecules, and Diseases* (1998) 24(4) Feb 28: 54–61;<br>Tsuboi et al. *JBC* Vol. 246, No. 24, Issue of December 25, pp. 7586-7594, 1971;<br>Arnold et al. *Pediatric Research* volume 8, pages 26–30 (1974);<br>JED Dyson, EA Noltmann. *JBC*, 243, ISSUE 7, P1401-1414, APRIL 1968;<br>Hines, Wolfe, *Biochemistry*. Vol. 2, No. 4, July Aug., 1963 |
| $K_{eq}$ | $0.36 \pm 0.11$ | | *eQuilibrator* (37C, pH = 7.4, pMg = 3, I = 0.25M) |

#### Phosphofructokinase (PFK)

*PFK reaction: F6P* + *ATP* ⇄ *F16BP* +*ADP*

*PFK genes:* Humans have 3 genes (PFKP, PFKPL, and PFKM) coding for PFK that have tissues-specific expression and distinct allosteric regulation. We focus on PFKP (also referred to as PFK-C) as it is the most abundant PFK isoform in proliferating cells based on proteomics data (***Supplementary File 1***).

*Regulation of PFK by small molecules:* PFKP, PFKPL, and PFKM are regulated by an unusually large number of small molecules including inhibition by citrate, Acyl-CoA (only shown for PFKM) and activation by phosphate, AMP (except for PFKP). In addition, PFK is allosterically inhibited by its substrate ATP and activated by its substrate F6P and products ADP and F16BP (except for PFKP).

*Structure of PFKP and number of binding sites for small molecules*: To formulate the general MWC equation for PFKP, we need to know the oligomeric state of PFKP, the number of binding sites for each metabolite, and whether any metabolites bind to the same site. Biochemical studies have shown that PFKP is a tetramer in solution. Structural, kinetic, and mutagenesis studies have shown that PFKP has at least four allosteric binding sites for ADP, Citrate, F26BP, and ATP and Phosphate, which compete for the same binding site (Appendix 1– figure 3***A***).

*General MWC rate equation for PFKP:* Based on the structural data described above we formulated the following rate equation based on the assumption that PFKP exists in an active state (*A*) with a catalytic rates 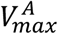 and 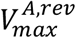 and an inactive state (*I*) with catalytic rates 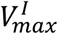 and 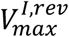. In the absence of metabolites, the ratio between the inactive and active states is *L*. Each substrate, product, and effector can have a different binding constant *K* for the active and inactive enzyme states. Note, that we calculated the 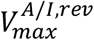 from the other kinetic constants using the Haldane relationship. Altogether, structural and regulatory knowledge of PFK translates into the following general MWC equation for PFK (note that the simplified kinetic ***Equation (50)*** was used in the model as described below):

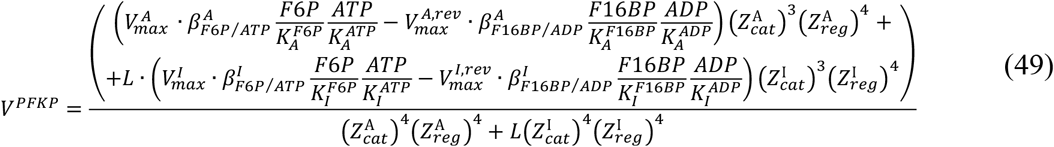

where

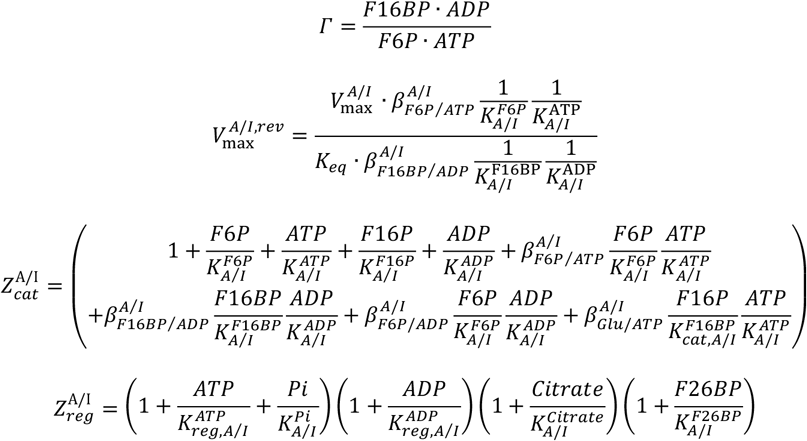

*Data used for PFKP:* We manually digitized 486 data points from 16 publications describing the rate of purified PFKP enzyme in the presence of different concentrations of F6P, ATP, F16BP, ADP, F26BP, Citrate and inorganic phosphate (*P*_*i*_). We considered data at pH 7-7.4 and in buffers without sulfate, which is a potent non-physiological activator of PFKP.

*Identification of minimal PFKP kinetic rate equation:* ***Equation (49)*** describes all possible kinetic parameters for PFKP based on structural and biochemical data. Some of these parameters might not be required to describe PFKP activity (e.g., *K*_*A*_ *= K*_*I*_, *β*_*A*_ *= β*_*I*_, *β = 0, β = 1*) and their inclusion might lead to poor prediction of enzyme rate under conditions not encountered in the fitting data due to overfitting. Therefore, we used regularization and cross-validation to determine if PFKP equation with a smaller number of kinetic parameters can achieve lower *Loss* test value of cross-validation. The Loss value was calculated using ***Equation (36)***. We set 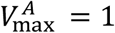 as this value is determined separately from the specific activity of the enzyme. We then performed four sequential rounds of regularization, as described in ***Section 2***. ***Estimation of Kinetic Parameters for MWC Enzymes***. Cross-validation *Loss* test value after removal of parameters after each cross-validation cycle is shown in Appendix 1–figure 3***B***. During the first regularization, we added terms 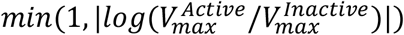, *min(1*, |*log(K* /_*Active*/_ *K*_*Inactive*_ )|) and *min(1*, |*log(β*_*Active*_/*β*_*Inactive*_ )|) to the *Loss* function to identify metabolites that bind with the same affinity to active and inactive conformation. We were able to set *β*_*A*_ *= β*_*I*_ for all *β* and *K*_*A*_ *= K*_*I*_ for 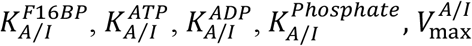, which decreased the number of parameters in ***Equation (49)*** from 30 to 21 without affecting the cross-validation test score (Appendix 1–figure 3***B***). During second and third regularizations, we added |*β*| and then *min(1*, |*log(β)*|) to the *Loss* function to identify *β =* 0 and *β = 1*, respectively. We were able to set *β =* 0 for 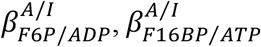 and *β = 1* for 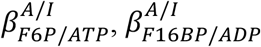, decreasing the number of parameters to 17 without affecting the cross-validation test score. During fourth round of cross-validation, we added terms *min(1*, |*log(1000*/*K)*|) to the *Loss* function to identify *K =*

∞. We were able to set *K =* ∞ for 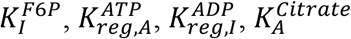 decreasing the number of parameters to 13 without an effect on the cross-validation test score. The final simplified kinetic rate equation for PFKP had 13 parameters (including 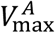 and *K*_*eq*_) instead of 30 and lower *Loss* test value for cross-validation (Appendix 1–figure 3***B***) compared to ***Equation (49)***:

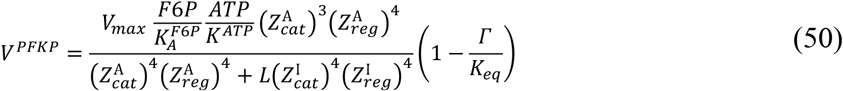

where

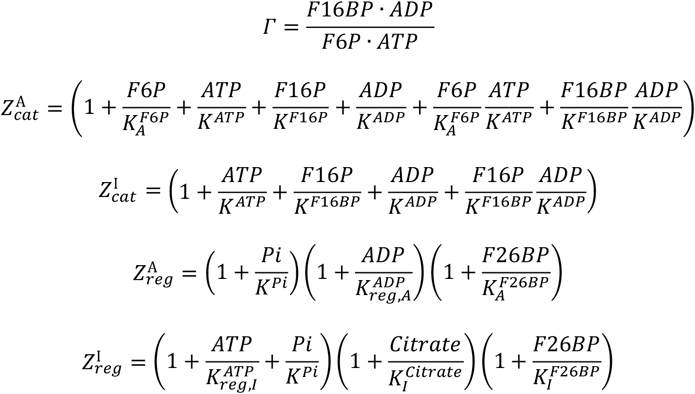

***Equation (50)*** was used to describe PFKP activity in our model. The least squares *Loss* function values that we obtained for the cross-validation train and test fits of the ***Equation (50)*** are 0.14 and 0.16, corresponding to an average error of 37% and 39% per data point, which is within the typical variability of *in vitro* kinetic experiments.

*PFKP protein bound metabolite equations:* We used the following binding equation to calculate fraction of PFKP bound to respective metabolites:

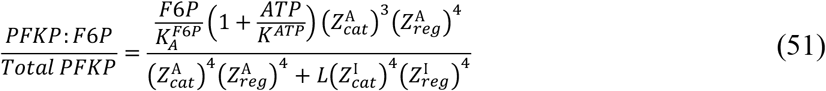

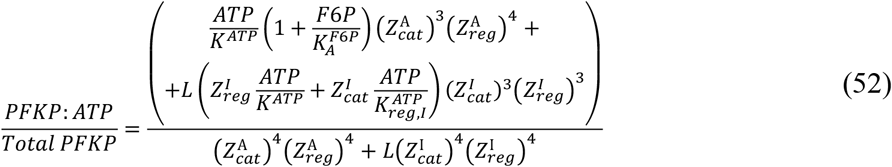

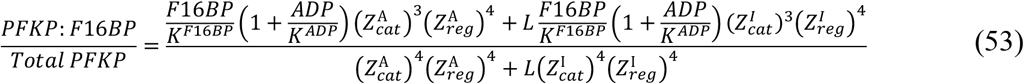

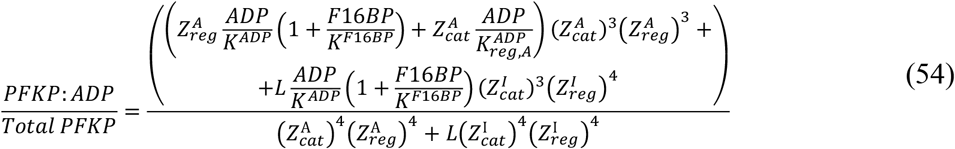

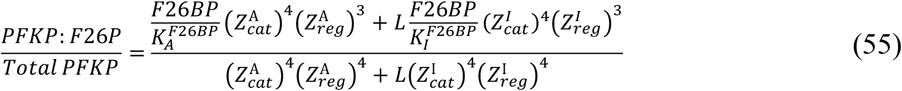

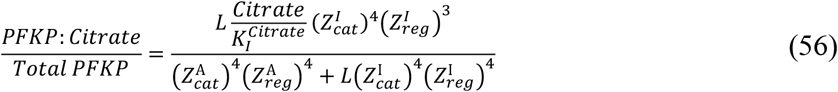

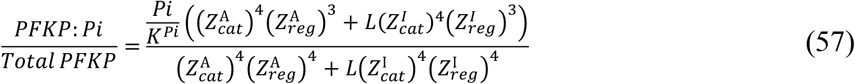

*Estimation of PFKP kinetic constants from data:* We used 10,000 fits to bootstrapped datasets to estimate kinetic parameters for ***Equation (50)*** and their confidence intervals. Appendix 1–figure 3***C*** shows that we obtain a narrow confidence interval for each of the parameters that show that our data is sufficient to uniquely estimate all the parameters of ***Equation (50)***. The fits of the kinetic equation to all data are shown in ***Appendix 1–figure 3D***.

*Reported PFKP regulation that was not included in our model:* We have not included several regulators of PFKP in our model. We hope that the following information will be useful for future efforts to improve the modeling of PFKP kinetic activity and glycolysis. Several regulators were reported to regulate PFKP at concentrations that are at least 10-fold higher than physiologically reported values including PEP with *K*_*I*_ > 1mM (***Boscá et al., 1982***), and 3PG with *K*_*I*_ > 1mM (***Tsai and Kemp, 1974***). Several regulators are known to affect other PFK isoforms but not PFKP including activation by F16BP and AMP (reported for all isoforms except PFKP (***Meienhofer et al., 1980***)), and inhibition by creatine phosphate (reported to inhibit PFKM but not PFKP specifically (***Tsai and Kemp, 1974***) but is later reported (***Fitch et al., 1979***) to be a contaminant creatine phosphate) and Acyl-CoA (reported only for muscle (***Jenkins et al., 2011***)). In addition, pH (***Ui, 1966***) and protein concentrations (***Reinhart, 1980***) are known to modulate the kinetic properties of PFK isoforms but we could not find enough kinetic data to incorporate the effect of protein concentration or pH. We did not include regulation of PFKP by citrate or 6-phosphogluconate (***Sommercorn et al., 1984***) in the model as these metabolites are produced by pathways other than glycolysis (i.e., TCA cycle and pentose phosphate pathway) and are likely involved in the coordination of glycolysis activity with TCA cycle and pentose phosphate pathway, respectively, which is beyond the scope of this report. Finally, there is evidence (***Kemp et al., 1976***) for the existence of more than two conformations of PFKP (i.e., inactive dimer with lower *V*_*max*_) but it is currently not clear if they are physiologically important, and the amount of currently available kinetic data is likely not sufficient to resolve the effect of additional conformations.

**Appendix 1–figure 3.**
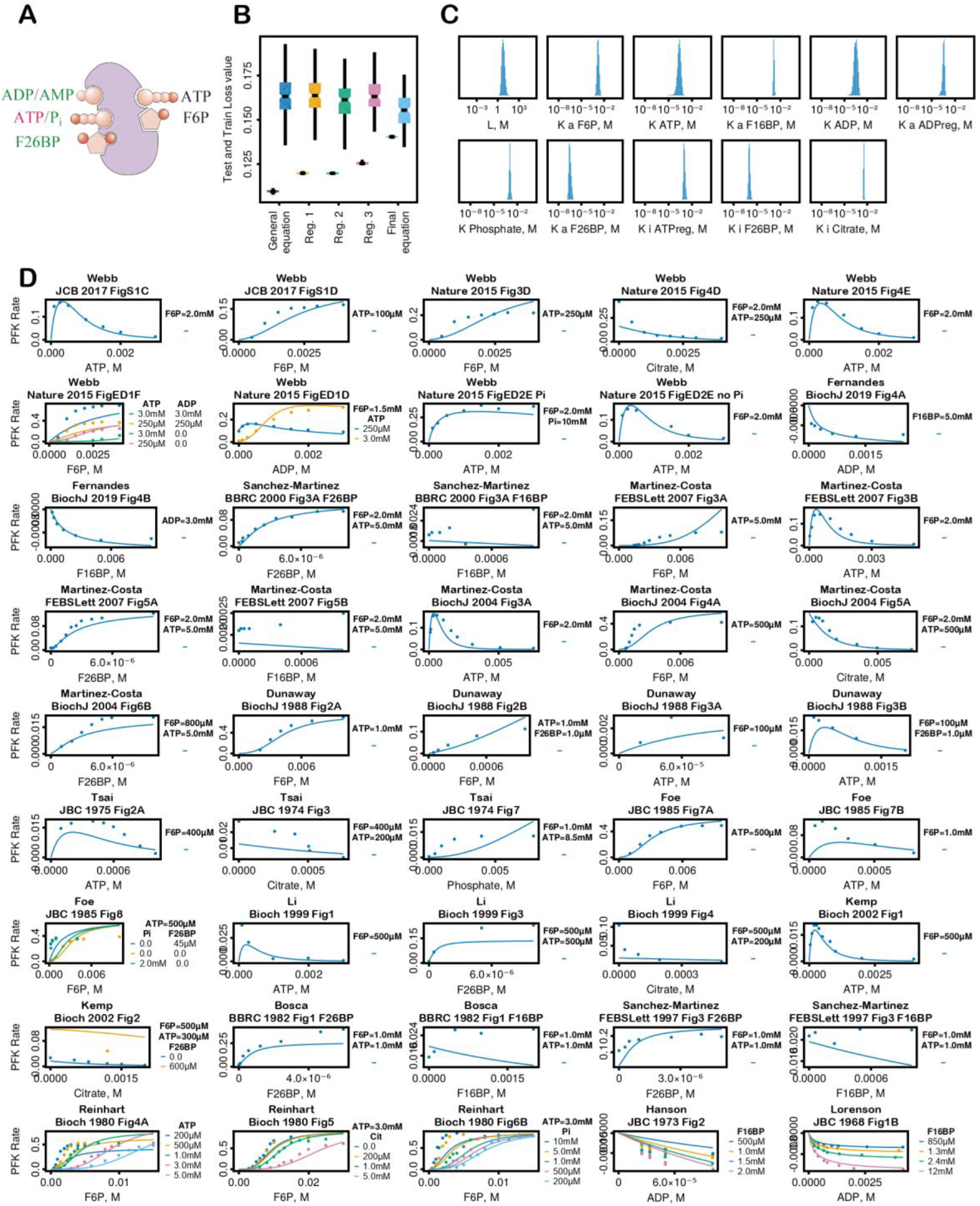
PFKP kinetic equation fitting. **(A)** Schematic of PFKP regulation, **(B)** Change of PFKP cross-validation fit (left dodge) and test (right dodge) *Loss* values after sequential rounds of regularization. ***Equation (49)*** corresponds to left most values and ***Equation (50)*** corresponds to right most values, **(C)** Histogram of estimated kinetic parameters of ***Equation (50)*** from fitting 10,000 bootstrapped datasets, **(D) *Equation (50)*** with final kinetic parameters plotted on top of all of the data used for fitting.

**Appendix 1–Table 4.** PFKP Kinetic Parameters. Mean ± standard deviation of parameter bootstrap values estimated by fitting of ***Equation (50)*** to data as described above.

| *Parameter* | *Mean ± SEM* | *Values* | *References* |
| --- | --- | --- | --- |
| *Molecular Weight, g/mol* | 85,596 |  | *UniProt: Q01813* |
| *L* | $6.9 \pm 4.0$ | *Data fitting* | |
| $K_A^{F6P}, \mu M$ | $1700 \pm 500$ | *Data fitting* | |
| $K^{ATP}, \mu M$ | $250 \pm 130$ | *Data fitting* | |
| $K^{F16BP}, mM$ | $1500 \pm 200$ | *Data fitting* | |
| $K^{ADP}, \mu M$ | $360 \pm 190$ | *Data fitting* | |
| $K^{Phosphate}, \mu M$ | $1100 \pm 300$ | *Data fitting* | |
| $K_{reg,I}^{ATP}, \mu M$ | $1100 \pm 300$ | *Data fitting* | |
| $K_{reg,A}^{ADP}, \mu M$ | $340 \pm 100$ | *Data fitting* | |
| $K_A^{F26BP}, \mu M$ | $0.34 \pm 0.12$ | *Data fitting* | |
| $K_I^{F26BP}, \mu M$ | $1.1 \pm 0.3$ | *Data fitting* | |
| $K_I^{Citrate}, \mu M$ | $3600 \pm 300$ | *Data fitting* | |
| $V_{max}, \frac{\mu mole}{min \cdot mg}$ | $80 \pm 22$ | 62.1, 59.27, 34, 41.2, 102, 180 | *Webb et al. J. Cell Biol. 2018, Vol. 216 No. 8 2305–2313;*<br>*Webb et al. NATURE, VOL 523, 2 JULY 2015, 111-114;*<br>*Martinez-Costa et al. FEBS Letters 581 (2007) 3033–3038;*<br>*Martinez-Costa et al. Biochem. J. (2004) 377, 77–84;*<br>*Foe, Kemp. JBC Vol.260, No.2, Issue of January 25, pp 726-730, 1985;*<br>*Li et al. Biochemistry 1999, 38, 16407-16412;* |
| $K_{eq}$ | $760 \pm 380$ | | *eQuilibrator (37C, pH = 7.4, pMg = 3, I = 0.25M)* |

#### Reversible non-enzymatic hydration/dehydration of DHAP and GAP

*Reactions:* DHAP(ketone) + H_2_O ⇄ DHAP(diol), GAP(ketone) + H_2_O ⇄ GAP(diol)

**Appendix 1–Table 5.**
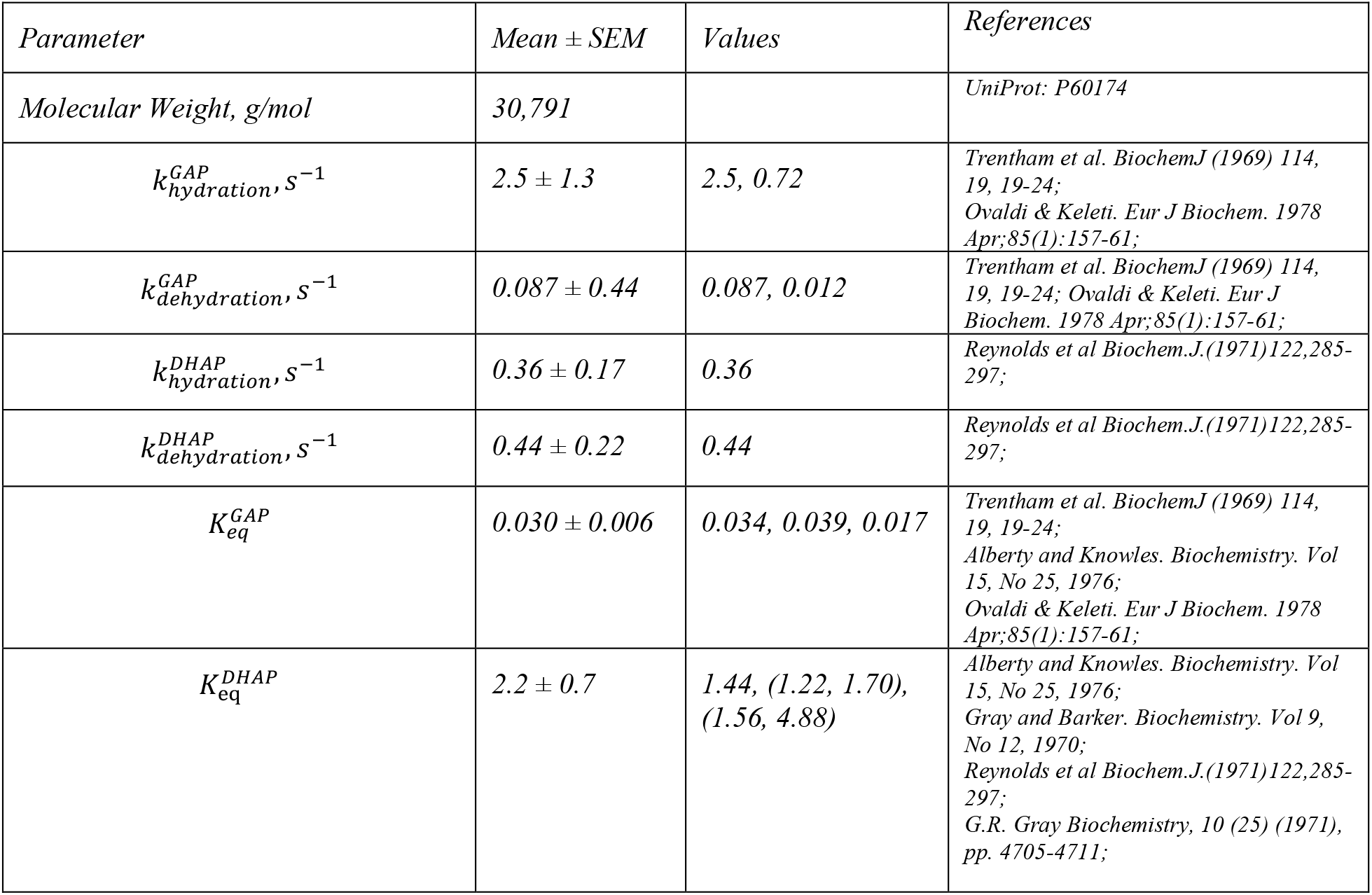
Kinetic Parameters for non-enzymatic hydration/dehydration of DHAP and GAP. Kinetic parameters were taken from the literature as is.

| *Parameter* | *Mean ± SEM* | *Values* | *References* |
| --- | --- | --- | --- |
| *Molecular Weight, g/mol* | *30,791* |  | *UniProt: P60174* |
| $k_{hydration}^{GAP}, S^{-1}$ | $2.5 \pm 1.3$ | 2.5, 0.72 | *Trentham et al. BiochemJ (1969) 114, 19, 19-24; Ovaldi & Keleti. Eur J Biochem. 1978 Apr;85(1):157-61;* |
| $k_{dehydration}^{GAP}, S^{-1}$ | $0.087 \pm 0.44$ | 0.087, 0.012 | *Trentham et al. BiochemJ (1969) 114, 19, 19-24; Ovaldi & Keleti. Eur J Biochem. 1978 Apr;85(1):157-61;* |
| $k_{hydration}^{DHAP}, S^{-1}$ | $0.36 \pm 0.17$ | 0.36 | *Reynolds et al Biochem.J.(1971)122,285-297;* |
| $k_{dehydration}^{DHAP}, S^{-1}$ | $0.44 \pm 0.22$ | 0.44 | *Reynolds et al Biochem.J.(1971)122,285-297;* |
| $K_{eq}^{GAP}$ | $0.030 \pm 0.006$ | 0.034, 0.039, 0.017 | *Trentham et al. BiochemJ (1969) 114, 19, 19-24; Alberty and Knowles. Biochemistry. Vol 15, No 25, 1976; Ovaldi & Keleti. Eur J Biochem. 1978 Apr;85(1):157-61;* |
| $K_{eq}^{DHAP}$ | $2.2 \pm 0.7$ | 1.44, (1.22, 1.70), (1.56, 4.88) | *Alberty and Knowles. Biochemistry. Vol 15, No 25, 1976; Gray and Barker. Biochemistry. Vol 9, No 12, 1970; Reynolds et al Biochem.J.(1971)122,285-297; G.R. Gray Biochemistry, 10 (25) (1971), pp. 4705-4711;* |

#### Aldolase (ALDO)

*ALDO reaction:* F16BP ⇄ GAP + DHAP

*ALDO genes:* Humans have three genes coding for ALDO – ALDOA, ALDOB, ALDOC. We focus on ALDOA as it is the most highly expressed ALDO isoform in proliferating mammalian cell lines (***Supplementary File 1***).

*ALDOA kinetic rate equation:* ALDO reaction mechanism involves covalent reaction intermediate between eneamine of DHAP and enzyme (referred to as E-DHAP). The reaction is sequential where F16BP reacts to yield GAP and E-DHAP, GAP is released, and then E-DHAP is hydrolyzed to produce DHAP (***Rose et al., 1987***). We use the following reaction scheme for ALDO:

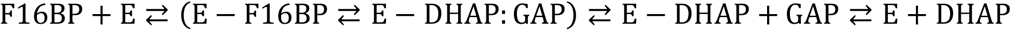

For completeness, we provide a derivation of the Ordered Uni Bi kinetic equation for ALDO using steady-state approximation (similar derivations are widely reported in the literature but we hope that readers will find it useful to have derivations of many kinetic rate equations reported in the same text). Steady-state approximation assumes that enzyme species do not change over time (i.e. their derivative are equal to zero) or change much faster than metabolite concentrations leading to the following set of equations for ALDO:

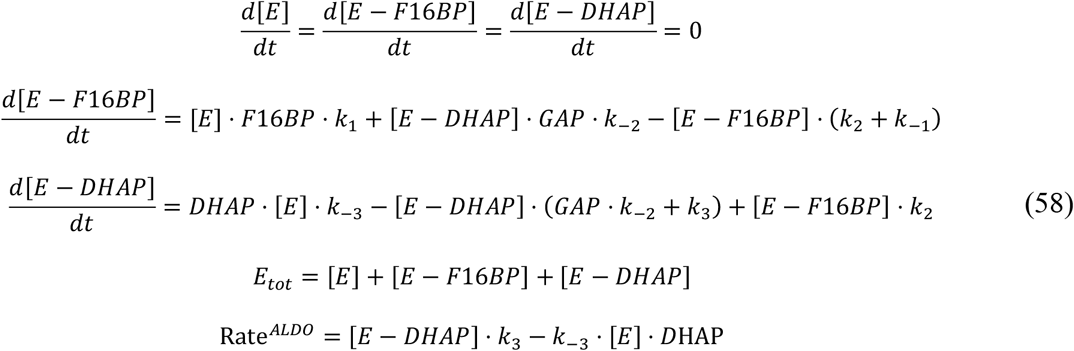

Solution of equations (58) for *V* ^*ALDO*^ yields:

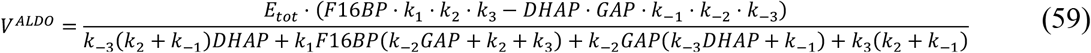

We can convert kinetic rate constants into parameters that are more easily measured experimentally like *K*_*M*_, *V*_*max*_, *K*_*eq*_, etc. The latter is achieved by taking the limits in the absence or presence of saturating concentrations of various substrates:

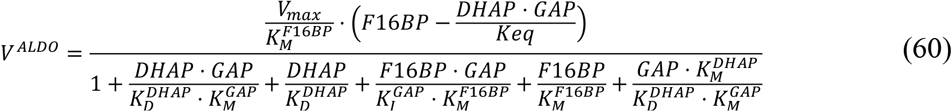

where 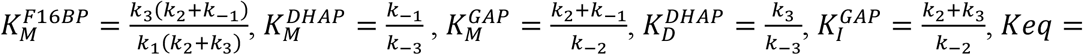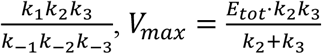, and *V* 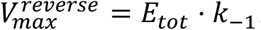. (Note that 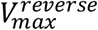 is not used in equation (60) but we report it for completeness). As a check, we confirm that all the constants are in units of concentrations.

*ALDOA protein bound metabolite equations:* We used the following equations, derived from equations, to calculate the amount of F16BP, GAP and DHAP bound to ALDO. Note that in our kinetic scheme complex of ALDO:F16BP represents a combined concentration of complex ALDO:F16BP or ALDO:GAP:DHAP. We made a simplifying assumption that equal fraction of ALDO:F16BP are ALDO:F16BP and ALDO:GAP:DHAP:

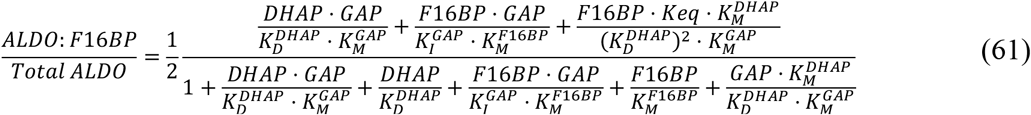

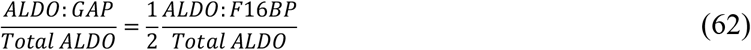

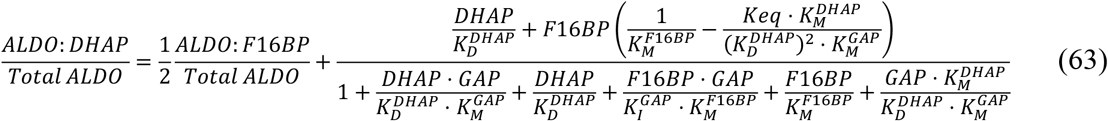

*Reported ALDOA regulation that was not included in our model:* GAP was reported to bind to free enzyme with ∼10µM affinity and exhibits substrate inhibition indicating that ALDO-GAP complex does not react with DHAP (***Rose and O’Connell, 1969***). We did not include such an interaction as data describing this is limited and in cells there is much more DHAP than GAP due to *K*_*eq*_ of TPI reaction so this interaction might not be physiologically relevant under most conditions. In addition, erythrose-4-phosphate and phosphate were reported to be a non-competitive and competitive inhibitor of ALDO with K_i_ *in the range of* 100 µM and 1mM, respectively. We have not included erythrose-4-phosphate because it is a product of pentose phosphate pathway and is not produced by glycolytic enzymes. Future extension of the model to pentose phosphate pathway will allow us to include this regulator. There were reports of ATP and ADP being inhibitors of ALDO but it was later shown that this was due to chelation of Mg^2+^ as ATP and ADP instead Mg-ATP and Mg-ADP were used in those assays (***Kasprzak and Kochman, 1980***).

**Appendix 1–Table 6.**
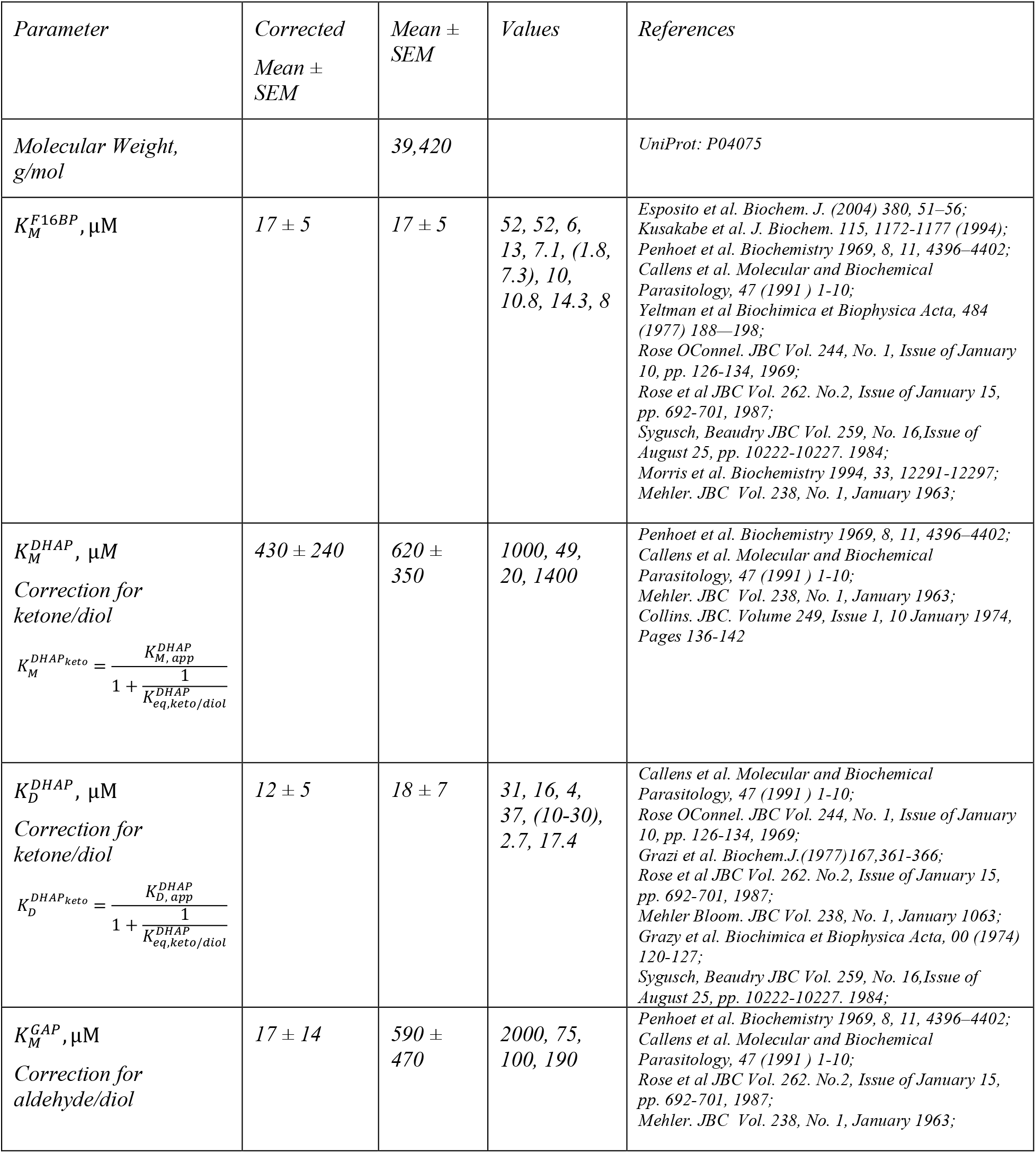
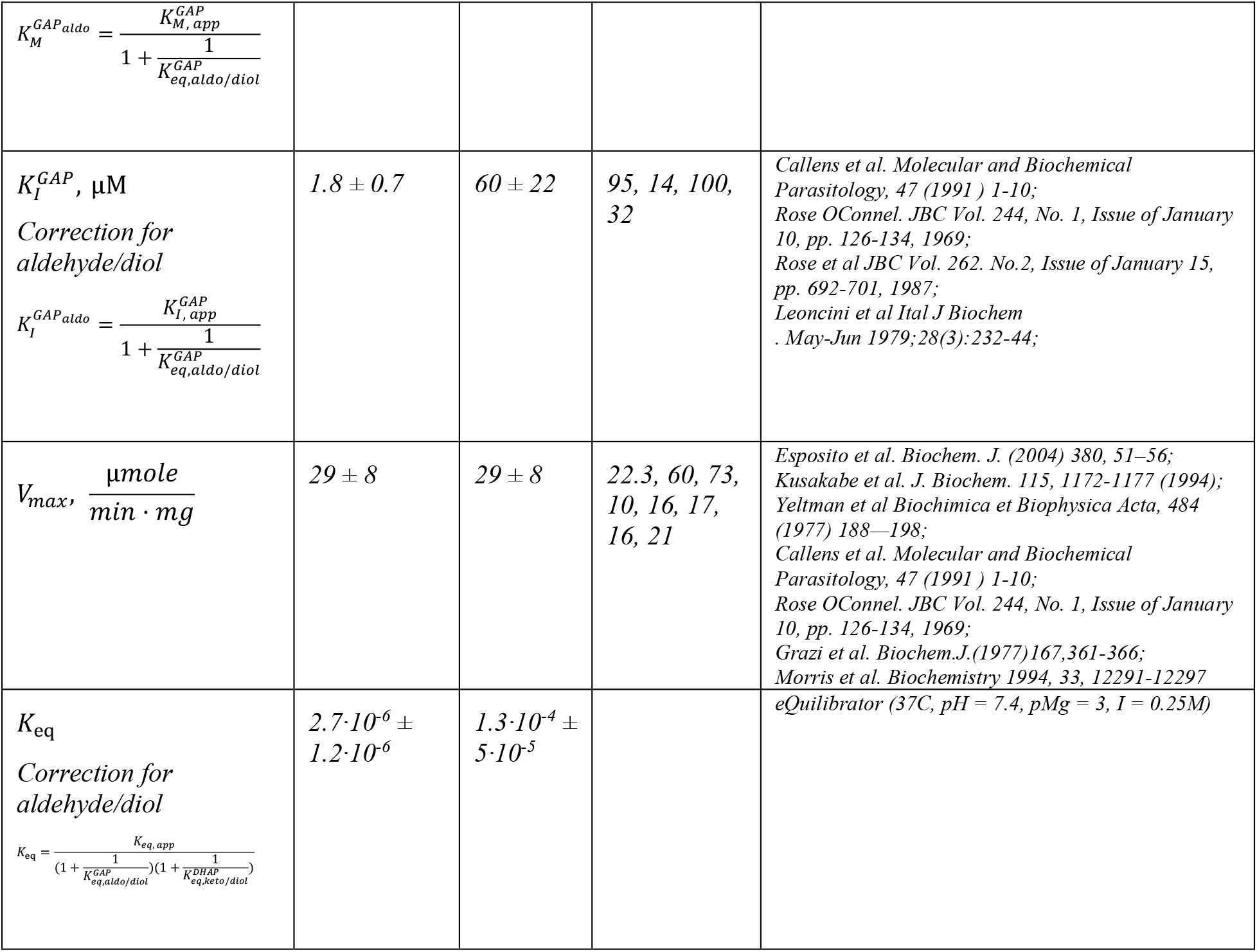
ALDOA kinetic parameters. Kinetic parameters were used as is from references below without fitting the equation to data. *K*_*M*_, *K*_*I*_, *K*_*D*_, *K*_*eq*_ values for GAP and DHAP need were adjusted for aldehyde/ketone vs diol forms as previously described (*Reynolds et al*., *1971*; *Trentham et al*., *1969*). GAP exists in solution as either aldehyde form or diol form, and DHAP as ketone or diol. Diols form after hydration of aldehydes or ketones. Aldehyde/ketone forms of GAP and DHAP are the substrates for TPI, ALDO and GAPDH. However, a significant fraction of DHAP (∼30%) and the majority of GAP (>95%) is present in diol form at equilibrium. If this is not considered, the apparent *K*_*M*_, *K*_*I*_, *K*_*D*_, *K*_*eq*_ values will be inaccurate.

#### Triose phosphate isomerase (TPI)

*TPI Reaction:* DHAP ⇄ GAP

*TPI genes:* Humans have one gene coding for TPI called TPI1.

*TPI kinetic equation:* We used the reversible Michaelis-Menten equation to describe the activity of TPI.

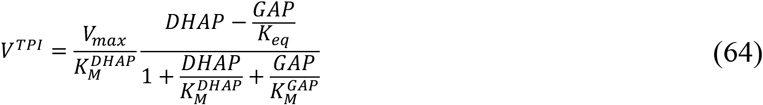

*TPI protein-bound metabolite equations:*

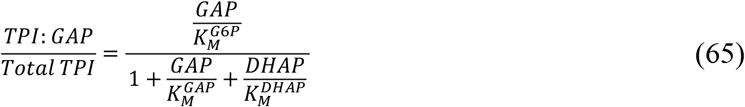

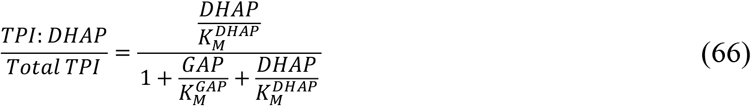

*TPI Kinetic Parameters:* Kinetic parameters were used as-is from references without fitting the equation to data but with correction for the prevalence of diol forms of GAP and DHAP described below. *K*_*M*_, *K*_*I*_, *K*_*D*_, *K*_*eq*_ values for GAP and DHAP need were adjusted for aldehyde/ketone vs diol forms as previously described (***Reynolds et al., 1971; Trentham et al., 1969***). GAP exists in solution as either aldehyde form or diol form, and DHAP as ketone or diol. Diols form after hydration of aldehydes or ketones. Aldehyde/ketone forms of GAP and DHAP are the substrates for TPI, ALDO and GAPDH. However, a significant fraction of DHAP (∼30%) and the majority of GAP (>95%) is present in diol form at equilibrium. If this is not considered, the apparent *K*_*M*_, *K*_*I*_, *K*_*D*_, *K*_*eq*_ values will be inaccurate.

*Reported TPI regulation that was not included in our model:* In addition, PEP has been described as an inhibitor of TPI (***Grüning et al., n*.*d***.) with *K*_*I*_ ≈ 0.2 mM. We did not include this interaction as the *K*_*I*_ is >10-fold higher than the intracellular concentration of PEP and TPI is already by far the most active enzyme in glycolysis due its high expression and high activity so its inhibition should have a minor if any effect on the pathway.

**Appendix 1–Table 7.** Kinetic parameters for TPI. Kinetic parameters were used as is from references below without fitting the equation to data.

| *Parameter* | *Corrected*<br>*Mean ±*<br>*SEM* | *Mean ±*<br>*SEM* | *Values* | *References* |
| --- | --- | --- | --- | --- |
| *Molecular Weight, g/mol* | *N/A* | *N/A* | *30,791* | *UniProt: P60174* |
| $K_M^{DHAP}, \mu M$<br>*Correction for ketone/diol*<br>$K_M^{DHAP_{keto}} = \frac{K_M^{DHAP_{app}}}{1 + \frac{1}{K_{eq,aldo/diol}^{DHAP}}}$ | *470 ± 80* | *680 ± 100* | *(250, 330),*<br>*(1500, 820,*<br>*690), 870,*<br>*(600,620),*<br>*(590,590,580)* | *Yuan et al Arch Bioch Bioph Vol 198, November, pp42-52, (1979); Gracy, R.W.; Methods Enzymol. 41B, 442-447 (1975); Burton & Waley, Biochim Biophys Acta. 1968 Mar 25;151(3):714-5; Kriestch et al. Eur J Biochem. 1970 Jun;14(2):289-300; Lee et al BBA, 1971 Jul 21;242(1):261-7;* |
| $K_M^{GAP}, \mu M$<br>*Correction for aldehyde/diol*<br>$K_M^{GAP_{aldo}} = \frac{K_M^{GAP_{app}}}{1 + \frac{1}{K_{eq,aldo/diol}^{GAP}}}$ | *11 ± 2* | *390 ± 20* | *(370, 260),*<br>*(340, 430, 550),*<br>*(460, 390, 350,*<br>*500), (320,320),*<br>*(420,400,400)* | *Yuan et al Arch Bioch Bioph Vol 198, November, pp42-52, (1979); Gracy, R.W.; Methods Enzymol. 41B, 442-447 (1975); Burton & Waley, Biochim Biophys Acta. 1968 Mar 25;151(3):714-5; Kriestch et al. Eur J Biochem. 1970 Jun;14(2):289-300; Lee et al BBA, 1971 Jul 21;242(1):261-7;* |
| $V_{max}, \frac{\mu mole}{min \cdot mg}$ | *6600 ± 800* | *6600 ± 800* | *(5360, 6900),*<br>*10236, 6700,*<br>*6400,*<br>*(7935,2302,723*<br>*1)* | *Yuan et al Arch Bioch Bioph Vol 198, November, pp42-52, (1979); Gracy, R.W.; Methods Enzymol. 41B, 442-447 (1975); Burton & Waley, Biochim Biophys Acta. 1968 Mar 25;151(3):714-5; Kriestch et al. Eur J Biochem. 1970 Jun;14(2):289-300; Lee et al BBA, 1971 Jul 21;242(1):261-7;* |
| $K_{eq}$<br>*Correction for aldehyde/diol*<br>$K_{eq} = K_{eq}^{app} \frac{1 + \frac{1}{K_{eq,aldo/diol}^{DHAP}}}{1 + \frac{1}{K_{eq,aldo/diol}^{GAP}}}$ | *0.0045 ± 0.0024* | *0.11 ± 0.06* | | *eQuilibrator (37C, pH = 7.4, pMg = 3, I = 0.25M)* |

#### Glyceraldehyde-3-phosphate dehydrogenase (GAPDH)

*GAPDH Reaction: GAP* + *NAD*^+^ + *Phosphate* ⇄ *BPG* + *NADH*

*GAPDH genes:* Humans have two genes coding for GAPDH called GAPDH and GAPDHS. We focused on GAPDH as it is the most abundant isoform in most tissues except testis that express GAPDHS.

*Regulation of GAPDH by small molecules:* GAPDH exhibits unusual cooperative regulation only by its substrates and products and not by other metabolites.

*Structure of GAPDH and number of binding sites for small molecules*: To formulate the general MWC equation for GAPDH, we need to know the oligomeric state of GAPDH, the number of binding sites for each metabolite, and whether any metabolites bind to the same site. Biochemical studies have shown that GAPDH is a tetramer in solution. Structural studies have shown that the following pairs of metabolites NAD^+^ and NADH, GAP and BPG, and BPG and Phosphate (Pi) bind to one subunit of GAPDH in a mutually exclusive manner (***Appendix 1– figure 4A***). For example, NAD^+^ and NADH cannot bind to one subunit simultaneously but can bind to different subunits of the same tetramer.

*General MWC rate equation for GAPDH:* Based on the structural data described above we formulated the following rate equation based on the assumption that GAPDH exists in an active state (*A*) with a catalytic rates 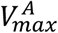 and 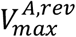 and an inactive state (*I*) with catalytic rates 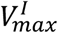 and 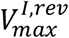. In the absence of metabolites, the ratio between the inactive and active states is *L*. Each substrate and product can have a different binding constant *K* for the active and inactive enzyme states. Note, that we calculated the 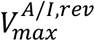 from the other kinetic constants using the Haldane relationship. Altogether, structural and regulatory knowledge of GAPDH translates into the following general MWC equation for GAPDH (note that the simplified kinetic ***Equation (68)*** was used in the model as described below):

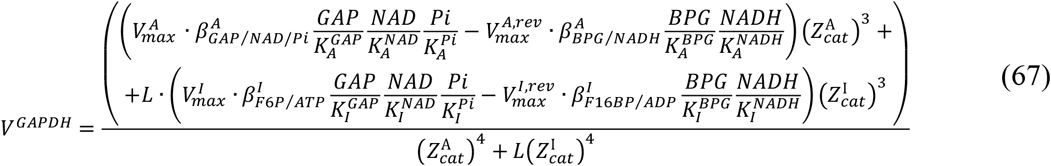

where

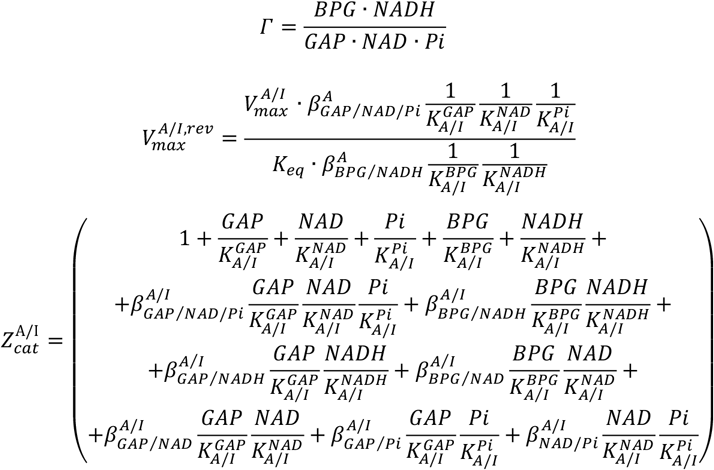

*Data used for GAPDH:* We estimated GAPDH kinetic parameters by curve fitting 522 data points from one publication (***Smith and Velick, 1972***).

*Identification of minimal GAPDH kinetic rate equation:* To identify the minimal MWC equation for GAPDH, parameter fitting was performed iteratively using different versions of MWC equation describing GAPDH activity with larger or smaller number of kinetic constants. We found that the best fit to data can be achieved by making one conformation catalytically inactive (i.e.,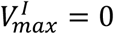) and by adding negative cooperativity between NADH and BPG when they are both bound on the same subunit when GAPDH is in the inactive conformation (i.e.,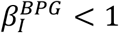).

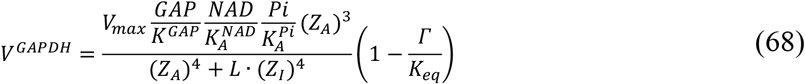

where

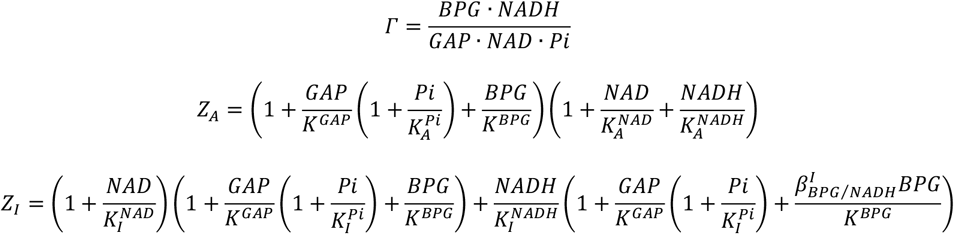

*GAPDH protein bound metabolite equation:* Fraction of GAPDH bound to a specific metabolite can be calculated by taking all the terms in *(Z*_*A*_)^*4*^ + *L* · *(Z*_*I*_)^*4*^ that contain the metabolite of interest and dividing it by *(Z*_*A*_)^*4*^ + *L* · *(Z*_*I*_)^*4*^. We use the following equations to calculate the levels of protein-bound metabolites:

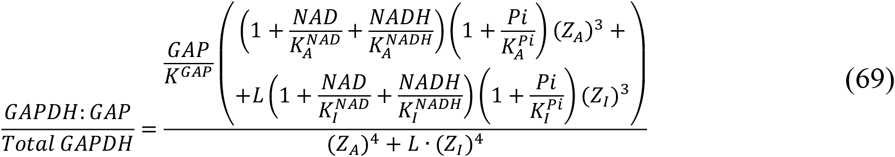

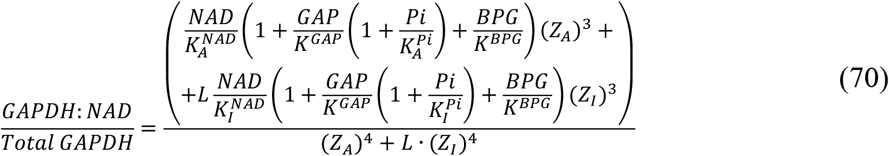

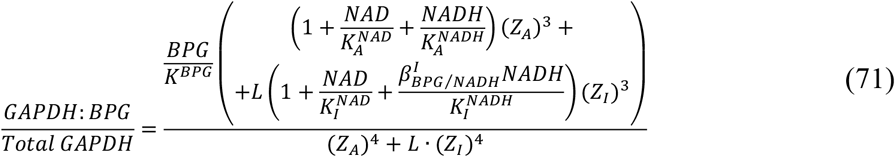

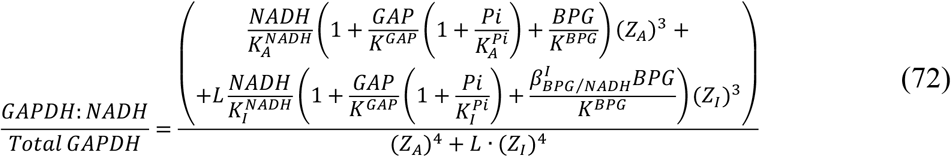

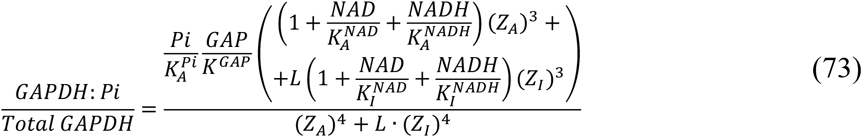

*Estimation of GAPDH kinetic constants from data:* Parameters estimated using fitting are displayed below and the fits are shown in ***Appendix 1–figure 4***. We estimated the uncertainty associated with parameter estimation using bootstrapping by rerunning the fitting 10,000 times on bootstrapped dataset with substitution. We adjusted kinetic and thermodynamic constants for the presence of aldehyde and diol forms of GAP as described above for ALDO and TPI. Finally, we calculated the specific activity of GAPDH by averaging values from several publications.

*Reported GAPDH regulation that was not included in our model:* We are not aware of any widely accepted regulators of GAPDH that we did not include in our description of GAPDH kinetics.

**Appendix 1–figure 4.**
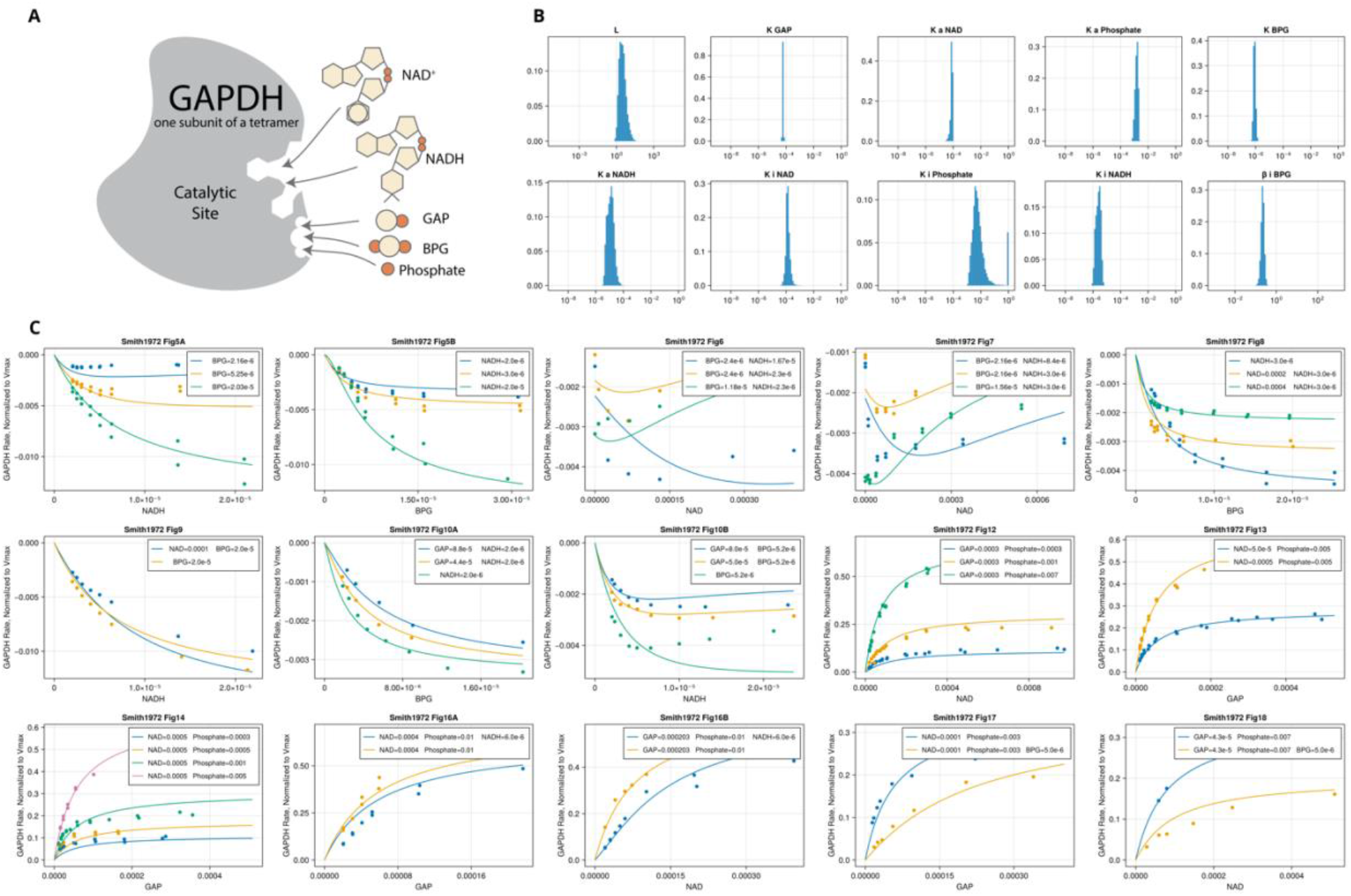
GAPDH fitting. **(A)** Binding of substrates and products to one GAPDH subunit that was used to derive MWC equation, **(B)** Distribution of bootstrap values of GAPDH kinetic constants, **(C)** GAPDH rate equation (lines) plotted on top of the data (points) used for fitting. All concentrations are M.

**Appendix 1–Table 8.** GAPDH Kinetic Parameters. Median ± median absolute deviation of parameter bootstrap values estimated by fitting of ***Equation (68)*** to data as described above.

| *Parameter* | *Median ± MAD* | *Values* | *References* |
| --- | --- | --- | --- |
| *Molecular Weight, g/mol* | 44,501 |  | UniProt: O14556 |
| *L* | 2.9 ± 1.7 | Data Fitting | |
| $K_M^{GAP}, \mu M$<br>*Correction for aldehyde/diol*<br>$K_M^{GAP_{aldo}} = \frac{K_{M,app}^{GAP}}{1 + \frac{1}{K_{eq,aldo/diol}^{GAP}}}$ *(Uncorrected values in brackets)* | 1.7 ± 0.1<br>(59 ± 3) | Data Fitting | |
| $K_A^{NAD}, \mu M$ | 78 ± 9 | Data Fitting | |
| $K_A^{Pi}, \mu M$ | 1,500 ± 400 | Data Fitting | |
| $K^{BPG}, \mu M$ | 0.82 ± 0.14 | Data Fitting | |
| $K_A^{NADH}, \mu M$ | 12 ± 7 | Data Fitting | |
| $K_I^{NAD}, \mu M$ | 130 ± 34 | Data Fitting | |
| $K_I^{Pi}, \mu M$ | 5,500 ± 3,800 | Data Fitting | |
| $K_I^{NADH}, \mu M$ | 2.5 ± 1.1 | Data Fitting | |
| $\beta_I^{BPG}$ | 0.22 ± 0.04 | Data Fitting | |
| $V_{max}, \frac{\mu mole}{min \cdot mg}$ | 130 ± 20 | 154, 164, 162, 98, 55, 117, 150-180 | Smith and Velick Vol. 247, No. 1. Issue of JBC January 10, pp. 273-284, 1972;<br>Ferdinant Biochem. J. (1964), 92, 578;<br>Kvassman et al Eur. J. Biochem. 172,427-431 (1988);<br>Wang, Alaupovic. ARCHIVES OF BIOCHEMISTRY AND BIOPHYSICS Vol. 205, No. 1, November, pp. 136-145, 1980;<br>Orsi, Cleland. BIOCHEMISTRY, VOL. 11, no. 1, 1 972;<br>Meunier, Dalziel Eur. J. Biochem. 82,483-492 (1978);<br>Wilder Biochimica et Biophysica Acta. 997 (1989) 65-77 |
| $K_{eq}$<br>*Correction for aldehyde/diol*<br>$K_{eq} = K_{eq}^{app} \left( 1 + \frac{1}{K_{eq,aldo/diol}^{GAP}} \right)$ *(Uncorrected values in brackets)* | 68 ± 20<br>(2.0 ± 0.4) | 3.5, 1.28, 1.36, 1.36, 4.77, 1.88, 1.26, 2.01 | Zhang et al Int J Mol Sci. 2017 Apr; 18(4): 876;<br>Cornell et al J Biol Chem. 1979 Jul 25;254(14):6522-7;<br>Greinert et al Fluid Phase Equilibria Volume 517, 1 August 2020, 112597 |

### Phosphoglycerate kinase (PGK)

*Reaction*: *BPG* +*ADP* ⇄ 3*PG* + *ATP*

*PGK genes:* Humans have two genes encoding PGK called PGK1 and PGK2. We focus on PGK1 as it is the most abundant isoform based in proliferating cells based on proteomics data (***Supplementary File 1***) while PGK2 is only expressed in the testis.

*PGK kinetic equation:* We used Rapid Equilibrium Random Binding approximation to describe PGK activity. The assumption of Rapid Equilibrium approximation is that the kinetics of binding and dissociation of substrates and products with enzyme are much faster than enzyme reaction rate so that we can assume that complexes between substrates, products and enzyme are in equilibrium. The assumption of Random Binding approximation is that substrates and product can bind to enzyme in random order. We used the following Rapid Equilibrium Random Binding Bi Bi kinetic equation for PGK (***Segel, Irwin, 1993***):

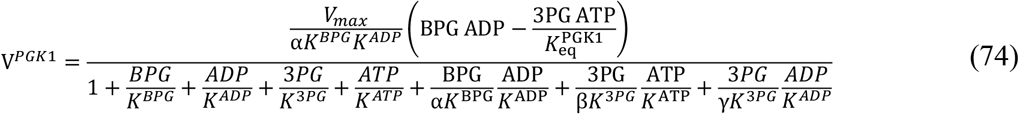

where *K*^*BPG*^, *K*^*ADP*^, *K*^*3PG*^, *K*^*ATP*^ are dissociation constants of enzyme and corresponding metabolites and α, β, γ are constants that determine whether binding of one ligand to enzyme increases (0 < α, β, γ < 1) or decreases (α, β, γ > 1) the affinity for the second ligand with α referring to a complex of enzyme with BPG and ADP, β – 3PG and ATP, and γ – 3PG and ADP. Full equation should also include δterm to describe complex of enzyme with BPG and ATP but we have omitted this term because as described below fitting shown that δ>> 1000 suggesting that BPG and ATP bind to PGK in mutually exclusive manner.

*PGK protein-bound metabolite equations:* We used the following equation to calculate the amount of BPG, ADP, 3PG and ATP bound to PGK1:

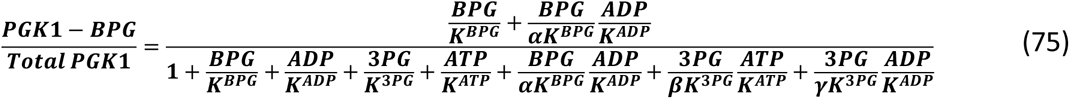

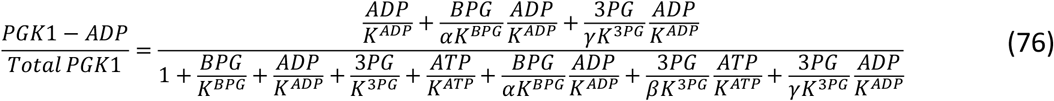

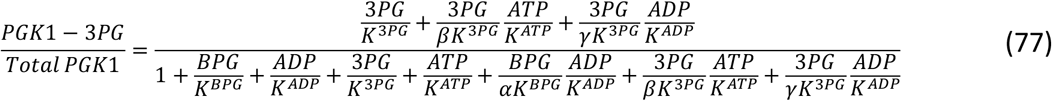

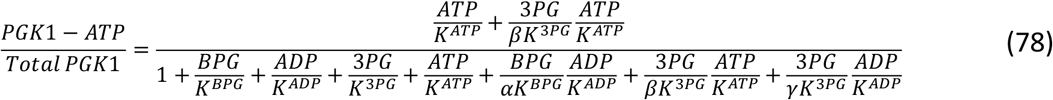

*Data used for PGK1:* We manually assembled 386 data points from four publications (***Ali and Brownstone, 1976; Lavoinne et al., 1983, 1979; Lee and O’Sullivan, 1975***) describing the rate of purified PGK1 enzyme in the presence of different concentrations of BPG, ADP, 3PG, and ATP.

*Estimation of PGK1 kinetic constants from data:* Parameters of ***Equation (74)*** estimated using fitting are displayed below and the fits are shown in ***Appendix 1–figure 5***. We estimated the uncertainty associated with parameter estimation using bootstrapping by rerunning the fitting 10,000 times on bootstrapped dataset with substitution.

*Reported PGK regulation that was not included in our model:* We are not aware of any known regulators of PGK1 that we did not include in our rate equation. We note that the kinetics of PGK1 deviate from Rapid Equilibrium Random Binding approximation at low concentrations of ATP where Lineweaver-Burke plots of *1/V* vs *1/S* deviate from linearity, which cannot be accounted for using ***Equation (74)***. Non-linear reciprocal plots require a kinetic equation that contains power terms of substrate or product concentrations in numerator and denominator. Power terms can result from several enzyme mechanisms including multiple substrates and product binding sites for the enzyme (this is not supported by PGK1 structural data), cooperative interaction between subunits (not applicable to PGK1 as it is a monomer), or a specific combination of kinetic constants of the reaction (***Ferdinand, 1966; King, 1956***). We believe that the latter is the most likely scenario for PGK1. We have not accounted for this behavior of PGK1 as it occurs at low unphysiological concentrations of ATP < 1mM but future models can include this behavior if warranted by the data.

**Appendix 1–figure 5.**
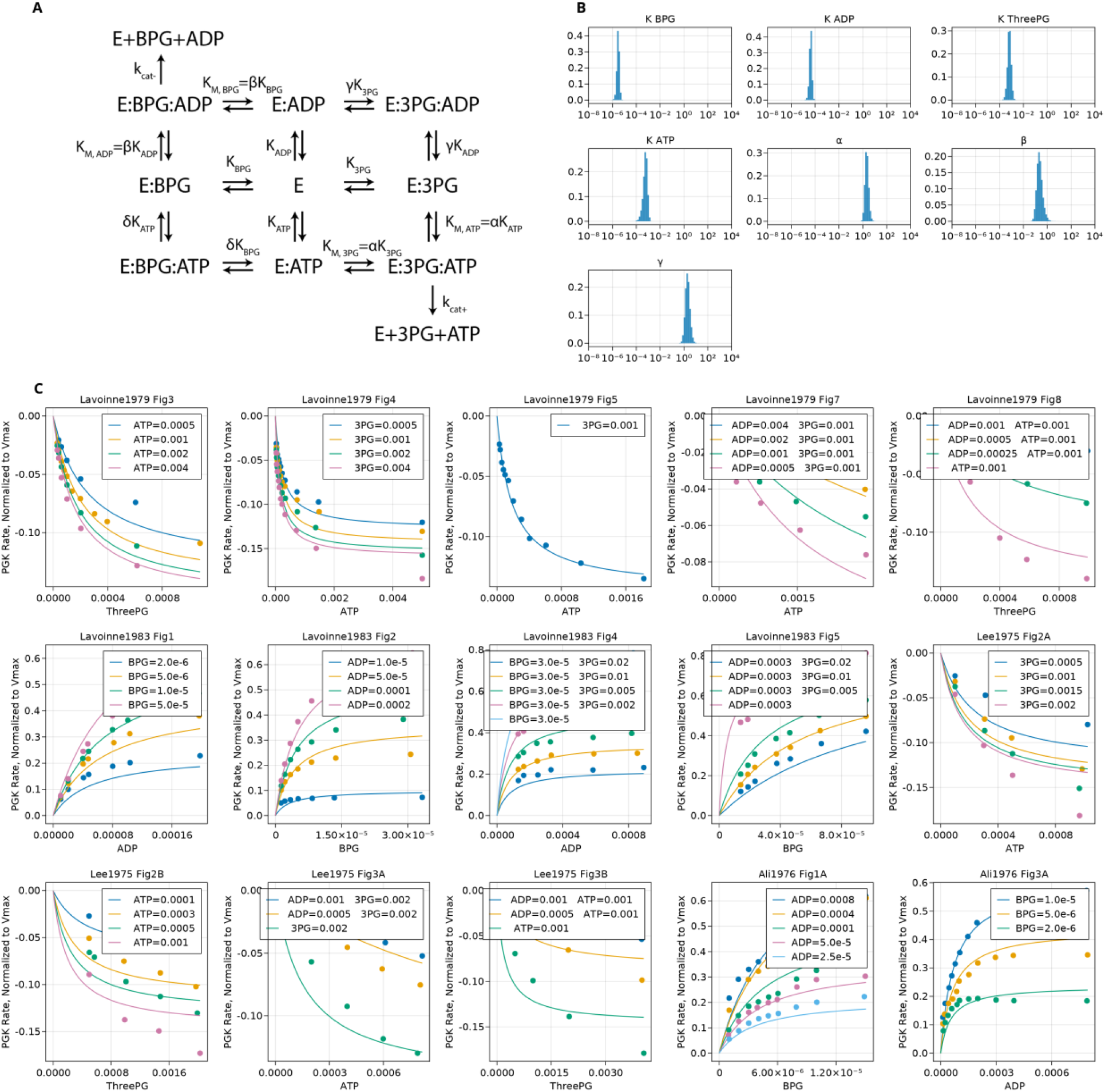
PGK1 fitting. **(A)** PGK1 kinetic scheme, **(B)** Distribution of bootstrap values of PGK1 kinetic constants, **(C)** PGK1 rate equation (lines) plotted on top of the data (points) used for fitting. All concentrations are M.

**Appendix 1–Table 9.** PGK1 Kinetic Parameters. Mean ± standard deviation of parameter bootstrap values estimated by fitting of ***Equation (74)*** to data as described above.

| *Parameter* | *Mean ± SEM* | *Values* | *References* |
| --- | --- | --- | --- |
| *Molecular Weight, g/mol* | *N/A* | *44,796* | *UniProt: P07205* |
| $K_{BPG}, \mu M$ | $3.0 \pm 0.7$ | *Data Fitting* | |
| $K_{ADP}, \mu M$ | $42 \pm 10$ | *Data Fitting* | |
| $K_{3PG}, \mu M$ | $660 \pm 220$ | *Data Fitting* | |
| $K_{ATP}, \mu M$ | $580 \pm 230$ | *Data Fitting* | |
| $\alpha$ | $2.1 \pm 0.7$ | *Data Fitting* | |
| $\beta$ | $0.22 \pm 0.11$ | *Data Fitting* | |
| $\gamma$ | $2.0 \pm 0.8$ | *Data Fitting* | |
| $V_{max}, \frac{\mu mole}{min \cdot mg}$ | $3500 \pm 900$ | 6780, 2900, 1900, 1838 (converted from reverse rate 300), 4290 (converted from reverse rate 700) | *Lavoinne et al BIOCHIMIE, 1983, 65, 211-220;*<br>*Lavoinne et al BIOCHIMIE, 1982, 64, 159-161;*<br>*Krietsch, Bucher. Eur. J. Biochem. 17 (1970)568-580;*<br>*Lavoinne et al BIOCHIMIE, 1979, 61, 1043-1053;*<br>*OSullivan, JBC, Vol. 250, No. 4, Issue of February 25, pp. 1275-1281, 1975;* |
| $K_{eq}$ | $2000 \pm 700$ | | *eQuilibrator (37C, pH = 7.4, pMg = 3, I = 0.2M)* |

#### Phosphoglycerate mutase (PGAM)

*Reaction:* 3PG ⇄ 2PG

*PGAM genes:* Humans have three gene coding for PGAM called PGAM1, PGAM2, and PGAM4. We focus on PGAM1 as it is the most highly expressed isoform in proliferating cell lines based on proteomics data (***Supplementary File 1***). In the literature, PGAM1 is also referred to as the brain isoform and PGAM2 as a muscle isoform.

*PGAM kinetic equation:* We used the reversible Michaelis-Menten equation to describe the activity of PGAM. This is a simplification of PGAM kinetic mechanisms as described below.

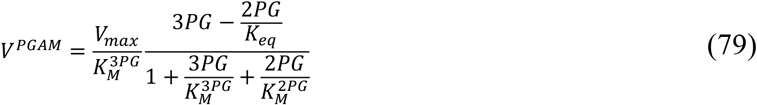

*PGAM protein-bound metabolite equations:* We used the following equation to calculate the amount of 3PG and 2PG bound to PGAM:

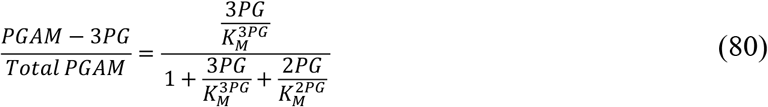

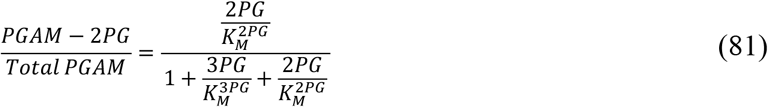

*Reported PGAM regulation that was not included in our model:* Kinetic mechanism of PGAM is known to be more complex than a reversible Michaelis-Menten equation. PGAM reaction proceeds through a Ping-Pong mechanism. A covalent intermediate of PGAM phosphorylated on His residue can bind either 2-PG or 3-PG and convert it to 3-PG or 2-PG, respectively, through a 2,3-bisphosphoglycerate (2,3-BPG) intermediate. For the reaction to proceed, PGAM must first be phosphorylated by 2,3-BPG or less efficiently by 1,3-BPG. In other words, 2,3-BPG or 1,3-BPG are essential cofactors of PGAM that must be present in catalytic amounts. The full description of PGAM kinetics will require a description of PGAM phosphorylation kinetics by 2,3-BPG and 1,3-BPG. In addition, enzymes that regulate 2,3-BPG levels in cells will have to be added to our model, including 2,3-BPG synthase (BPGM), 2,3-BPG phosphatase (TIGAR), and possibly other enzymes. The presence of 2,3-BPG is not essential for glycolysis activity as BPGM knockout cells have increased 3-PG levels but otherwise normal glycolysis activity while having no 2,3-BPG (***Oslund et al., 2017***). Therefore, we made a simplifying assumption that phosphorylated PGAM is the only form of the enzyme in our model. Future iterations of the model will be able to account for PGAM activity more accurately by incorporating PGAM kinetic rate equation in the presence of 2,3-BPG and 1,3-BPG and by adding enzymes required for 2,3-BPG metabolism.

**Appendix 1–Table 10.** PGAM Kinetic Parameters. We used kinetic parameters as is from the literature without fitting. Kinetic measurements (***Bartrons and Carreras, 1982***) suggest similar activity for PGAM1 and PGAM2. Therefore, we used kinetic constants from data describing either PGAM1 or PGAM2 activity.

| *Parameter* | *Mean ± SEM* | *Values* | *References* |
| --- | --- | --- | --- |
| *Molecular Weight, g/mol* | *N/A* | 28,804 | *UniProt: P18669* |
| $K_M^{3PG}, \mu\text{M}$ | $270 \pm 60$ | (168, 250 (chicken muscle)), 600 (chicken muscle), (200, 100 (pig)), 255 (rabbit muscle), 400 (human erythrocytes), 200 | *Rose, Dube. J Biol Chem, 1978 Dec 10;253(23):8583-92; Grisolia, Cleland. Biochemistry. 1968 Mar;7(3):1115-21; Bartrons Carreras. Biochimica et Biophysica Acta, 708 (1982) 167-177; Stankiewicz, Hass. Vol. 261, No. 27, Issue of September 25, pp. 12715-12721 1986; Atauri et al. Biochimica et Biophysica Acta 2005; Miller Haas. Biochem Biophys Res Commun. 1980 Jul 16;95(1):132-9;* |
| $K_M^{2PG}, \mu\text{M}$ | $19 \pm 7$ | 14 (chicken muscle), 10 (chicken muscle), 32 | *Rose, Dube. J Biol Chem, 1978 Dec 10;253(23):8583-92; Rodwell et al. J Biol Chem. 1957 Oct;228(2):875-90; Britton, Clark. Biochem. J.(1972)130,397-410;* |
| $V_{max}, \frac{\mu\text{mole}}{\text{min} \cdot \text{mg}}$ | $3000 \pm 900$ | 2493 (1197 s<sup>-1</sup>, chicken muscle), 2060 (chicken muscle), 8000, (chicken muscle), (4760, 3680 (converted from 137,000, 106,000, $\mu\text{mole/min per mole}$), 500, 1940, 420 | *Rose, Dube. J Biol Chem, 1978 Dec 10;253(23):8583-92; Rodwell et al. J Biol Chem. 1957 Oct;228(2):875-90; Grisolia, Cleland. Biochemistry. 1968 Mar;7(3):1115-21; Bartrons Carreras. Biochimica et Biophysica Acta, 708 (1982) 167-177; Stankiewicz, Hass. Vol. 261, No. 27, Issue of September 25, pp. 12715-12721 1986; Britton, Clark. Biochem. J.(1972)130,397-410; Miller Haas. Biochem Biophys Res Commun. 1980 Jul 16;95(1):132-9;* |
| $K_{eq}$ | $0.18 \pm 0.05$ | | *eQuilibrator (37C, pH = 7.4, pMg = 3, I = 0.2M)* |

#### Enolase (ENO)

*Reaction:* 2PG ⇄ PEP

*ENO genes:* Humans have three gene coding for enolase ENO1, ENO2, and ENO3. We focus on ENO1 as it is the most highly expressed isoform in proliferating mammalian cells based on proteomics data (***Supplementary File 1***).

*ENO kinetic equation:* We used the reversible Michaelis-Menten equation to describe the activity of ENO.

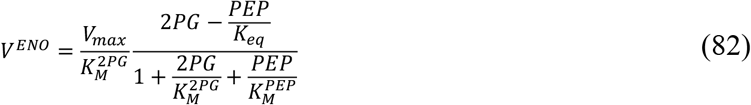

*ENO protein-bound metabolite equations:* We used the following equation to calculate the amount of 2PG and PEP bound to ENO:

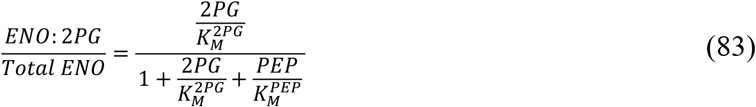

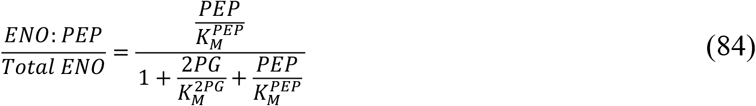

**Appendix 1–Table 11.** ENO Kinetic Parameters. We used kinetic parameters as is from the literature without fitting. Kinetic constants for mammalian ENO1, ENO2 and ENO3 are similar so we combined kinetic parameters measured in any of these enzymes (***Suzuki et al., 1980***).

| *Parameter* | *Mean ± SEM* | *Values* | *References* |
| --- | --- | --- | --- |
| *Molecular Weight, g/mol* | *N/A* | *47,169* | *UniProt: P06733* |
| $K_M^{2PG}, \mu\text{M}$ | $40 \pm 9$ | *17, (5.7, 5.8), 16, 16.3, (57, 45, 36), (90, 90), 50* | *Thompson et al J. Am. Chem. Soc. 2021, 143, 2, 785–797;*<br>*Kornblatt et al. Biochem Cell Biol Feb-Mar 1989;67(2-3):103-7;*<br>*Wang, Himoe. JBC Vol. 249, No. 12, Issue of June 25, pp. 3895-3902;*<br>*V. Hannaert et al. Eur. J. Biochem. 270, 3205–3213 (2003);*<br>*Suzuki et al. J Biochem. 1980 Jun;87(6):1587-94;*<br>*Farrar, Deal. Journal of Protein Chemistry, Vol. 14, No. 6, 1995;*<br>*Gorsich et al. Journal of Protein Chemistry, Vol. 18, No. 1, 1999;* |
| $K_M^{PEP}, \mu\text{M}$ | $160 \pm 30$ | *(51, 45), 238, (150, 190, 140), (250, 170), 250* | *Kornblatt et al. Biochem Cell Biol Feb-Mar 1989;67(2-3):103-7;*<br>*V. Hannaert et al. Eur. J. Biochem. 270, 3205–3213 (2003);*<br>*Suzuki et al. J Biochem. 1980 Jun;87(6):1587-94;*<br>*Farrar, Deal. Journal of Protein Chemistry, Vol. 14, No. 6, 1995;*<br>*Gorsich et al. Journal of Protein Chemistry, Vol. 18, No. 1, 1999;* |
| $V_{\max}, \frac{\mu\text{mole}}{\text{min} \cdot \text{mg}}$ | $71 \pm 9$ | *82, 107, 27, 30, (87, 68, 51), (65, 91), 82.2* | *Thompson et al J. Am. Chem. Soc. 2021, 143, 2, 785–797;*<br>*Kornblatt et al. Biochem Cell Biol Feb-Mar 1989;67(2-3):103-7;*<br>*Wang, Himoe. JBC Vol. 249, No. 12, Issue of June 25, pp. 3895-3902;*<br>*V. Hannaert et al. Eur. J. Biochem. 270, 3205–3213 (2003);*<br>*Suzuki et al. J Biochem. 1980 Jun;87(6):1587-94;*<br>*Farrar, Deal. Journal of Protein Chemistry, Vol. 14, No. 6, 1995;*<br>*Gorsich et al. Journal of Protein Chemistry, Vol. 18, No. 1, 1999;* |
| $K_{\text{eq}}$ | $4.4 \pm 1.0$ | | *eQuilibrator (37C, pH = 7.4, pMg = 3, I = 0.2M)* |

#### Pyruvate kinase (PK)

*Reaction: PEP* +*ADP* ⇄ *Pyruvate* + *ATP*

*PK genes:* Humans have 2 genes (PKM, PKLR) coding for PK that have tissue-specific expression and distinct allosteric regulation. In addition, both genes can express isoforms with distinct allosteric regulation like PKM1 and PKM2. We focus on PKM2 as it is the most abundant PK isoform in proliferating cells based on proteomics data (***Supplementary File 1***).

*Regulation of PKM2 by small molecules:* PKM2 is regulated by allosteric activators F16BP, histidine and serine, and allosteric inhibitors phenylalanine, alanine, tryptophan, methionine, valine, and proline. In addition, PKM2 is allosterically activated by its substrate PEP (***Yuan et al., 2018***).

*Structure of PKM2 and number of binding sites for small molecules*: To formulate the general MWC equation for PKM2, we need to know the oligomeric state of PKM2, the number of binding sites for each metabolite, and whether any metabolites bind to the same site. Biochemical studies have shown that PKM2 is tetrameric in solution but can dissociate into dimers and monomers in dilute solutions. Structural, kinetic, and mutagenesis studies (***Chaneton et al., 2012; Yuan et al., 2018***) have shown that PKM2 has two allosteric binding sites that are distinct from catalytic site: one for F16BP and another for amino acids (***Appendix 1–figure 6A***).

*General MWC rate equation for PKM2:* Based on the structural data described above we formulated the following rate equation based on the assumption that PKM2 exists in active state (*A*) with catalytic rates 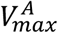 and 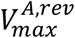 and an inactive state (*I*) with catalytic rates 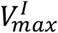 and 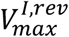. In the absence of metabolites, the ratio between the inactive and active states is *L*. Each substrate, product and effector can have a different binding constant *K* for the active and inactive enzyme state. Note, that we calculated the 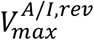 from the other kinetic constants using the Haldane relationship. We have only considered regulation by F16BP since it’s a product of glycolysis and also by phenylalanine, which is the most potent amino acid regulator and we reasoned that including this regulator will allow us to better constraint the parameters of MWC equation for PKM2 even though phenylalanine is not included in glycolysis model. Altogether, structural and regulatory knowledge of PKM2 translates into the following general MWC equation for PKM2 (note that the simplified kinetic ***Equation (86)*** was used in the model as described below):

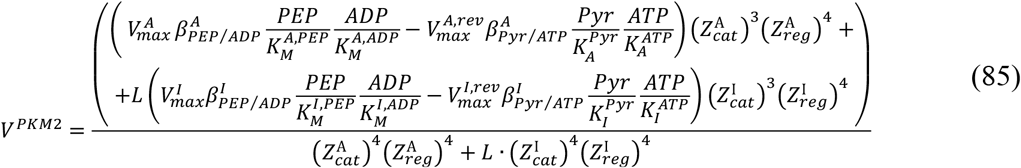

where

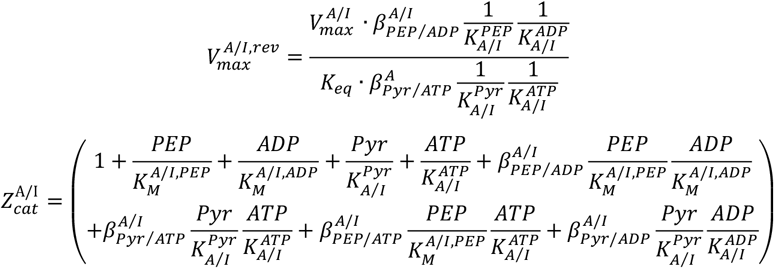

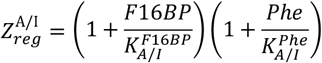

*Data used for PKM2:* We manually digitized 591 data points from 14 publications describing the rate of purified PKM2 enzyme in the presence of different concentrations of PEP, ADP, Pyruvate, ATP, F16BP.

*Identification of minimal PKM2 kinetic rate equation:* ***Equation (85)*** describes all possible kinetic parameters for PKM2 based on structural and biochemical data. Some of these parameters might not be required to describe PKM2 activity (e.g., *K*_*A*_ *= K*_*I*_, *β*_*A*_ *= β*_*I*_, *β = 0, β = 1*) and their inclusion might lead to poor prediction of enzyme rate under conditions not encountered in the fitting data due to overfitting. Therefore, we used regularization and cross-validation to determine if PKM2 equation with a smaller number of kinetic parameters can achieve lower *Loss* test value of cross-validation. The Loss value was calculated using equation (36). We set 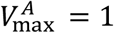 as this value is determined separately from the specific activity of the enzyme. We then performed four sequential rounds of regularization, as described in ***Section 2***. ***Estimation of Kinetic Parameters for MWC Enzymes***. Cross-validation *Loss* test value after removal of parameters after each cross-validation cycle is shown in ***Appendix 1–figure 6B***. During the first regularization, we added terms 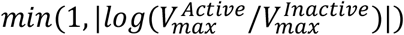, *min(1*, |*log(K*_*Active*_/*K*_*Inactive*_)|) and *min(1*, |*log(β*_*Active*_ /*β*_*Inactive*_)|) to the *Loss* function to identify metabolites that bind with the same affinity to active and inactive conformation. We were able to set *β*_*A*_ *= β*_*I*_ for 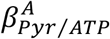 and 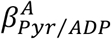 and *K*_*A*_ *= K*_*I*_ for 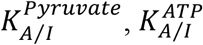, and 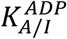, which decreased the number of parameters in ***Equation (85)*** from 24 to 19 without affecting the cross-validation test score (***Appendix 1–figure 6B***). During second and third regularizations, we added |*β*| and then *min(1*, |*log(β)*|) to the *Loss* function to identify *β =* 0 and *β = 1*, respectively. We were able to set *β = 1* for all *β* except 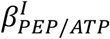 and we also removed the term 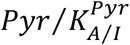 as high values of *β* terms involving Pyr indicated that it preferentially binds to enzyme after ATP and ADP are bound to active site. Second and third regularizations further decreased the number of parameters to 14 without affecting the cross-validation test score. During fourth round of cross-validation, we added terms *min(1*, |*log(1000*/*K)*|) to the *Loss* function to identify *K =* ∞. We were able to set *K =* ∞ for 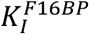 decreasing the number of parameters to 13 without an effect on the cross-validation test score. The final simplified kinetic rate equation for PKM2 had 13 parameters (including 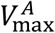, 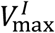 and *K*_*eq*_) instead of 24 and similar *Loss* test value for cross-validation (***Appendix 1–figure 6B***) compared to ***Equation (85)***:

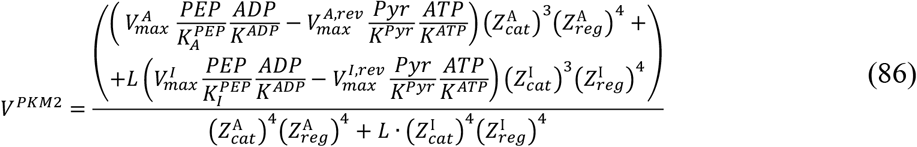

where

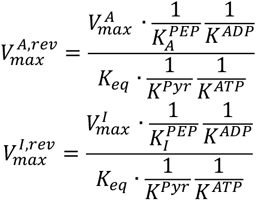

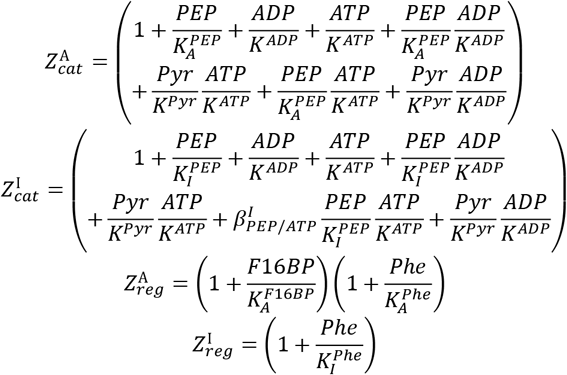

***Equation (86)*** was used to describe PKM2 activity in our model. The least squares *Loss* function values that we obtained for the cross-validation train and test fits of the ***Equation (86)*** are 7.1% and 9.3%, corresponding to an average error of 27% and 30% per data point, which is within the typical variability of the *in vitro* kinetic experiments.

*PKM2 protein bound metabolite equations:* We used the following binding equation to calculate fraction of PFKP bound to respective metabolites:

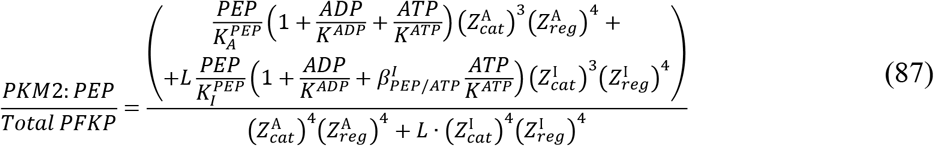

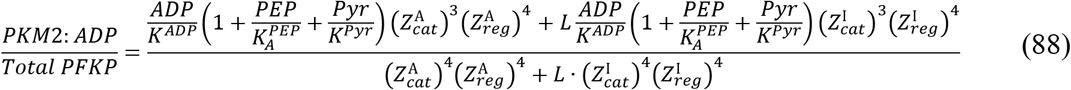

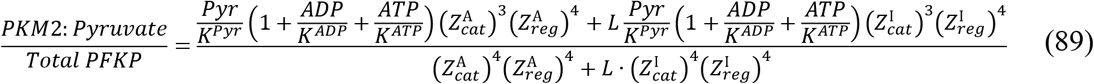

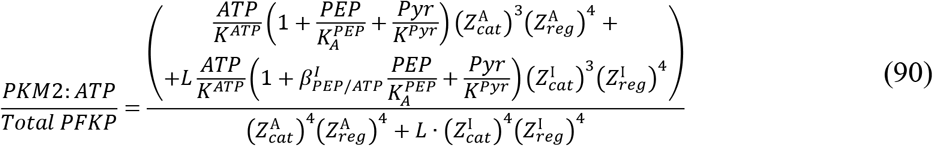

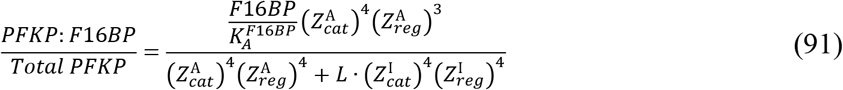

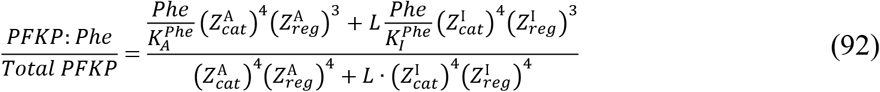

*Estimation of PKM2 kinetic constants from data:* We used 10,000 fits to bootstrapped datasets to estimate kinetic parameters for ***Equation (86)*** and their confidence intervals. ***Appendix 1–figure 6*** shows that we obtain narrow confidence interval for each of the parameters that shows that our data is sufficient to uniquely estimate all 12 parameters of ***Equation (86)***. The fits of the kinetic equation to all data are shown in ***Appendix 1–figure 6***.

*Reported PKM2 regulation that was not included in our model:* Several amino acids have been reported to be allosteric regulators of PKM2 that we did not consider as they are not substrates or products of glycolysis. Effect of F16BP on PK activity is slow and requires preincubation to achieve steady-state so we only considered publication where preincubation was reported to get the most accurate description of steady-state PK kinetic properties (***Pogson, 1968***). In some publications, F16BP and phenylalanine were reported to affect the Vmax of the enzymes (***Liu et al., 2020; Nandi et al., 2020; Yuan et al., 2018***). The reason for the discrepancy is not fully understood. We have included data from studies where Vmax varied and studies where it did not so our results represent an average of these likely inconsistent descriptions of PKM2 regulation. Some reports (***Nagao et al., 1977***) suggest that this discrepancy might be due to measurements of PKM2 activity in dilute conditions (e.g., 0.002 mg/ml) where PKM2 is known to dissociate into monomers and dimers and this might cause the changes in Vmax as the monomer are believed to be less active. Intracellular conditions with PKM2 level >1mg/ml favor tetrameric form whose *V*_*max*_ is not affected by effectors so the effect of regulators on Vmax might not be physiologically relevant in live cells.

**Appendix 1–figure 6.**
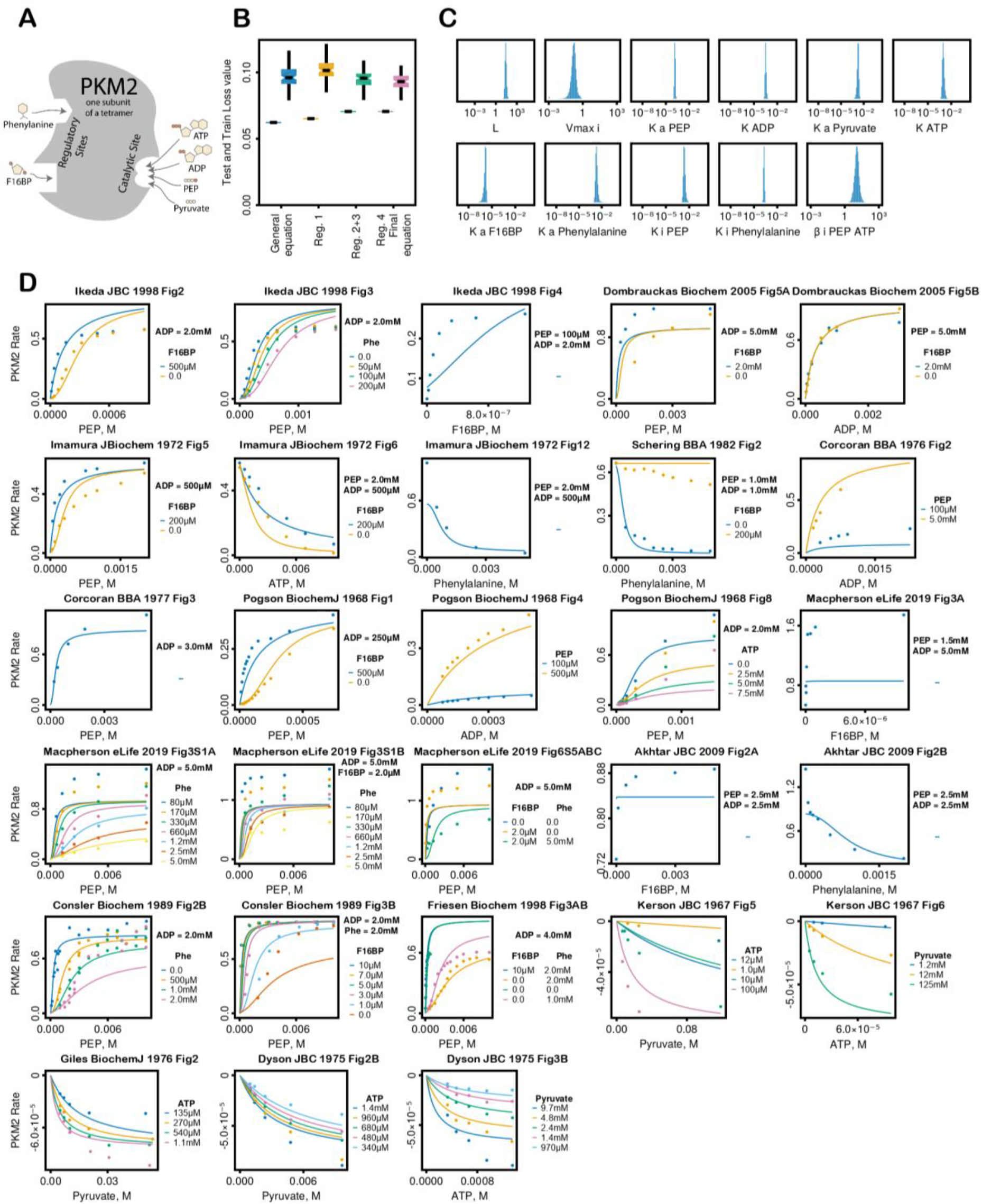
PKM2 kinetic equation fitting. **(A)** Schematic of PKM2 regulation, **(B)** Change of PKM2 cross-validation fit (left dodge) and test (right dodge) *Loss* values after sequential rounds of regularization. ***Equation (85)*** corresponds to left most values and ***Equation (86)*** corresponds to right most values, **(C)** Histogram of estimated kinetic parameters of ***Equation (86)*** from fitting 10,000 bootstrapped datasets, **(D) *Equation (86)*** with final kinetic parameters plotted on top of all of the data used for fitting.

**Appendix 1–Table 12.** PKM2 Kinetic Parameters. Median ± median absolute deviation of parameter bootstrap values estimated by fitting of ***Equation (86)*** to data as described above.

| *Parameter* | *Median ± MAD* | *Values* | *References* |
| --- | --- | --- | --- |
| *Molecular Weight, g/mol* | 57,937 |  | UniProt: P14618-1 |
| *L* | 26 ± 7 | Data fitting | |
| $K_A^{PEP}, \mu M$ | 150 ± 20 | Data fitting | |
| $K^{ADP}, \mu M$ | 320 ± 50 | Data fitting | |
| $K^{Pyruvate}, \mu M$ | 1800 ± 400 | Data fitting | |
| $K^{ATP}, \mu M$ | 940 ± 250 | Data fitting | |
| $K_A^{F16BP}, \mu M$ | 1.4 ± 0.4 | Data fitting | |
| $K_A^{Phenylalanine}, \mu M$ | 2100 ± 600 | Data fitting | |
| $K_I^{PEP}, \mu M$ | 2400 ± 600 | Data fitting | |
| $K_I^{Phenylalanine}, \mu M$ | 200 ± 30 | Data fitting | |
| $\beta_{PEP/ATP}^I$ | 12 ± 5 | Data fitting | |
| $V_{max}^A, \frac{\mu mole}{min \cdot mg}$ | 540 ± 100 | 520, 530, 405, 248, 770, 1341, 416, 780, 361, 397, 225 | Imamura, K., and Tanaka, T. (1982) *Methods Enzymol.* 90, 150–165;<br>Ikeda, Naguchi. *JBC.* Vol. 273, No. 20, Issue of May 15, pp. 12227–12233, 1998;<br>Nagao et al. *J Biochem.* 1977 Nov;82(5):1331-45;<br>Dombrauckas et al *Biochemistry* 2005, 44, 9417-9429;<br>Imamura et al. *J. Biochem.*, 72, 1001-1015 (1972);<br>Schering et al. *Biochimica et Biophysica Acta*, 717 (1982) 337-347;<br>Corcoran et al. *Biochimica et Biophysica Acta*, 446 (1976) 96–104;<br>Tanaka et al. *The Journal of Biochemistry*, Vol. 62, No. 1, 196;<br>Macpherson et al. *eLife* 2019;8:e45068;<br>Ibsen et al. *Biochemistry* 1981, 20, 1497-1506;<br>Freisen et al. *Biochemistry*, Vol. 37, No. 9, 1998; |
| $V_{max}^I, \frac{\mu mole}{min \cdot mg}$ | 86 ± 41 =<br>$=V_{max}^A \times$<br>$\times 0.16 \pm 0.07$ | Data fitting | |
| $K_{eq}$ | 20,000 ± 6,000 | | *eQuilibrator* (37C, pH = 7.4, pMg = 3, I = 0.25M) |

#### Lactate dehydrogenase (LDH)

*Reaction*: *Pyruvate* + *NADH* ⇄ *Lactate* + *NAD*^+^

*LDH genes:* Humans have three gene coding for LDH called LDHA, LDHB, LDHC. LDHA, also known as M or muscle isoform of LDH, and LDHB, also known as H or heart isoform of LDH, are both highly expressed in proliferating cells (***Supplementary File 1***). LDHC is expressed in testis. LDHA and LDHB have similar kinetic properties.

*LDH kinetic equation:* LDH reaction mechanism proceeds through ordered binding and release of substrates and products through the kinetic reaction scheme called Theorell-Chance mechanism (***Zewe and Fromm, 1962***) shown in ***Appendix 1–figure 7A***. Theorell-Chance is a simplification of ordered Bi Bi mechanism where rate limiting steps are binding and dissociation of NAD or NADH with the enzyme and ternary complexes of enzyme and substrates or products are not kinetically significant.

For completeness, we provide a derivation of Theorell-Chance equation for LDH using steady state approximation (this equation is widely reported in the literature, but we hope that readers will find it useful to have derivations of kinetic rate equations reported in the same text that make all the assumptions explicit). Steady state approximation assume that enzyme species do not change over time (i.e., their derivative are equal zero) or change much faster than metabolite concentrations leading to the following set of equations for LDH:

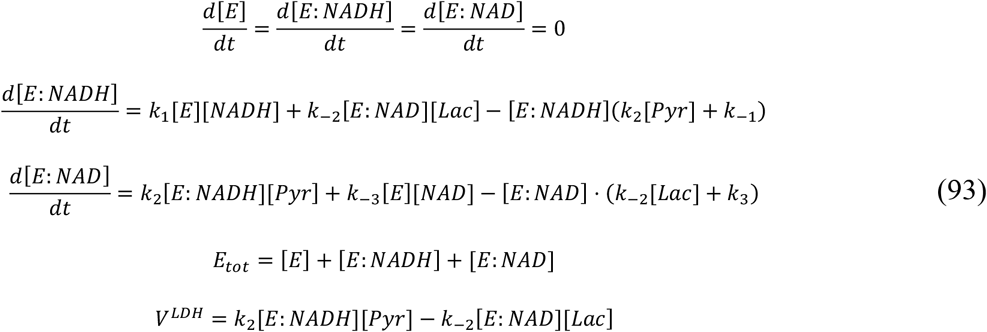

Solution of equations above yields the following rate equation for LDH:

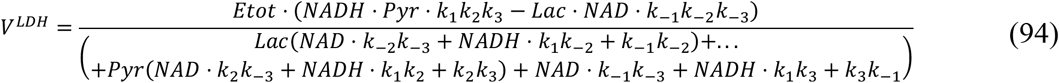

We can rewrite this equation using apparent kinetic constants, which can be calculated by taking limits of ***Equation (94)*** in presence of 0 or ∞ concentrations of NADH, Pyruvate, NAD^+^ and Lactate:

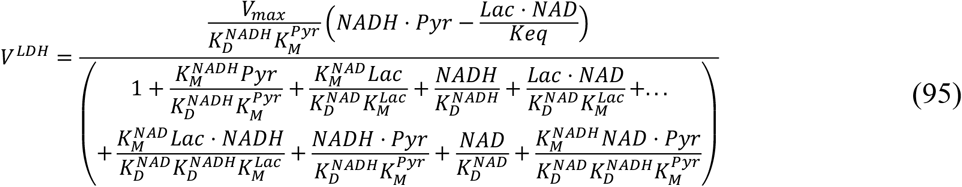

where 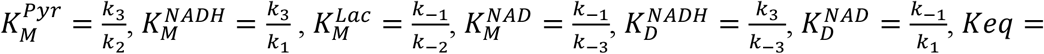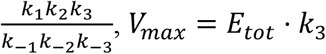, and 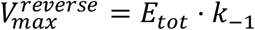. (Note that 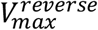 is not used in equation (95) but we report it for completeness).

*LDH protein-bound metabolite equations:* We used the following equations, derived from ***Equations (93)***, to calculate the amount of NAD^+^, NADH bound to LDH (note that Theorell-Chance mechanism assumes no significant level of complexes of enzyme with Lactate or Pyruvate, so we do not include these):

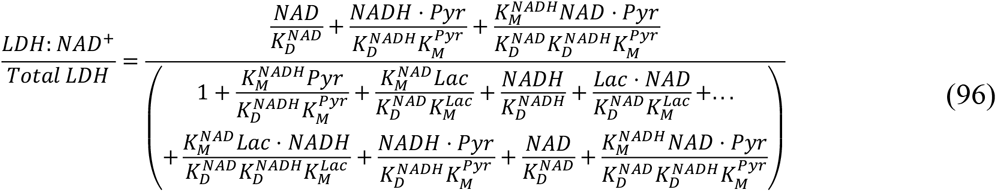

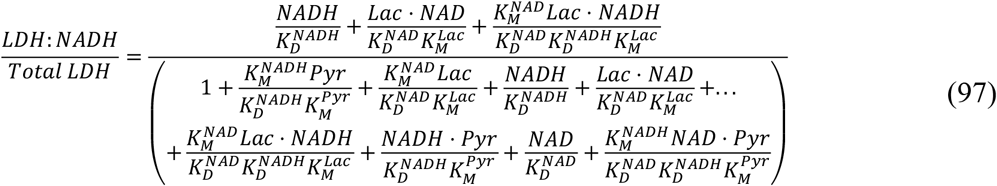

*Data used for LDH:* We manually assembled 406 data points from 3 publications (***Wang, 1977; Zewe and Fromm, 1965, 1962***) describing the rate of purified LDH enzyme in the presence of different concentrations of Pyruvate, NADH, Lactate, and NAD.

*Estimation of LDH kinetic constants from data:* Parameters of ***Equation (95)*** estimated using fitting are displayed below and the fits are shown in ***Appendix 1–figure 7***. We estimated the uncertainty associated with parameter estimation using bootstrapping by rerunning the fitting 10,000 times on bootstrapped dataset with substitution.

*Reported LDH regulation that was not included in our model:* LDH exhibits substrate inhibition by pyruvate and lactate that is believed to be due to the formation of inactive ternary complexes LDH-NAD-Pyr and LDH-NADH-Lac (***Zewe and Fromm, 1962***). We have not included substrate inhibition as *K*_*I*_ for substrate inhibition is an order of magnitude higher than *K*_*M*_ for corresponding substrates, and it was difficult to find kinetic data that uses pyruvate or lactate concentrations that are above *K*_*I*_.

**Appendix 1–figure 7.**
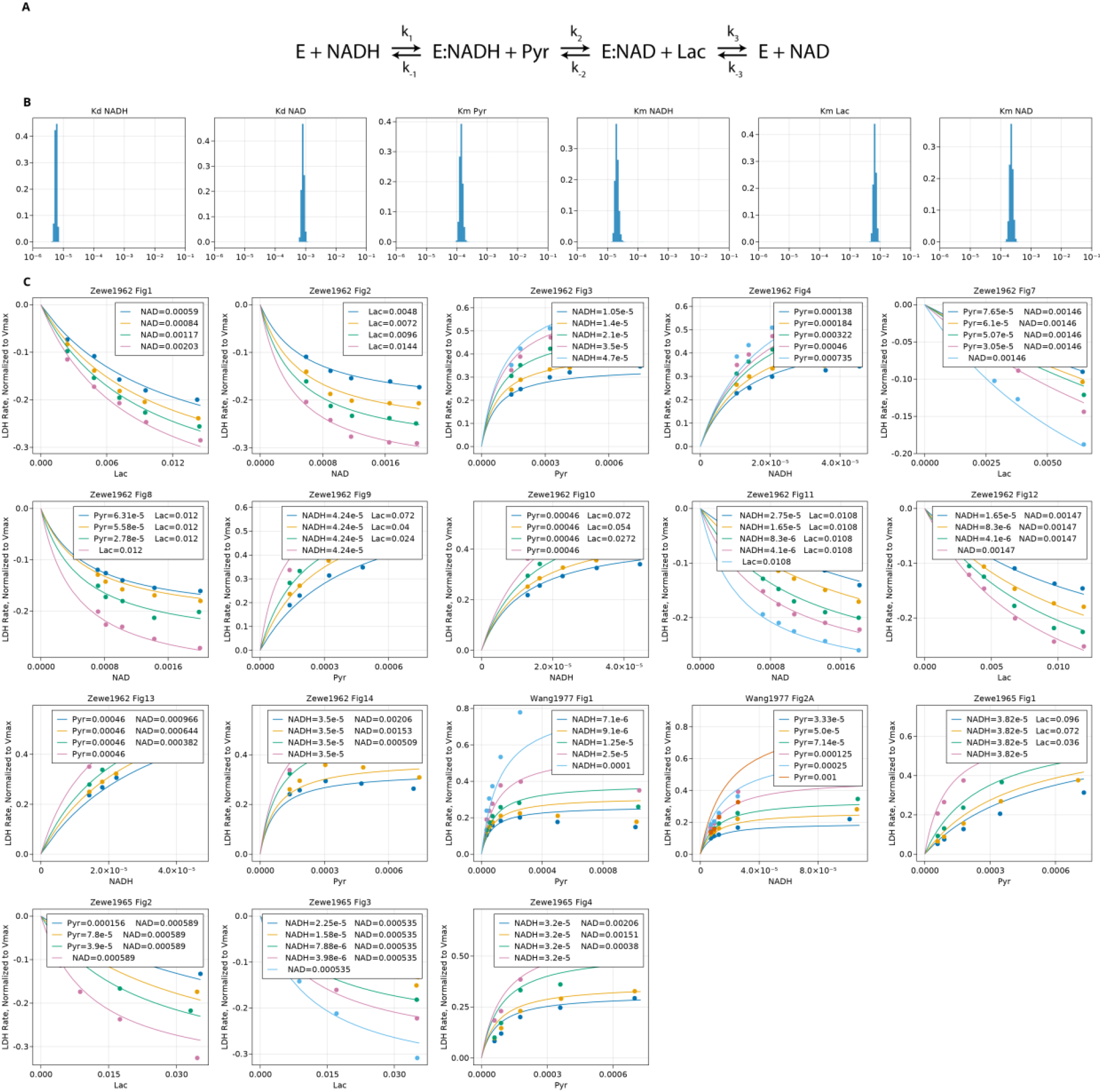
LDH fitting. **(A)** LDH kinetic scheme, **(B)** Distribution of bootstrap values of LDH kinetic constants, **(C)** LDH rate equation (lines) plotted on top of the data (points) used for fitting. All concentrations are M.

**Appendix 1–Table 13.** LDH Kinetic Parameters. Mean ± standard deviation of parameter bootstrap values estimated by fitting of ***Equation (95)*** to data as described above.

| *Parameter* | *Mean ± SEM* | *Values* | *References* |
| --- | --- | --- | --- |
| *Molecular Weight, g/mol* | *N/A* | *36,638* | *UniProt: P07195* |
| $K_M^{Pyr}, \mu M$ | $140 \pm 20$ | *Data Fitting* | |
| $K_M^{NADH}, \mu M$ | $20 \pm 2$ | *Data Fitting* | |
| $K_M^{Lac}, \mu M$ | $6600 \pm 700$ | *Data Fitting* | |
| $K_M^{NAD}, \mu M$ | $220 \pm 30$ | *Data Fitting* | |
| $K_D^{NADH}, \mu M$ | $5.7 \pm 0.4$ | *Data Fitting* | |
| $K_D^{NAD}, \mu M$ | $810 \pm 80$ | *Data Fitting* | |
| $V_{max}, \frac{\mu mole}{min \cdot mg}$ | $320 \pm 40$ | (209, 199, 193), 452 (converted from 276 s-1), 401 (converted from 245s-1), 366, 426 (converted from 260s-1) | *Pridgar et al. CLIN. CHEM. 30/8, 1353-1357 (1984); Jackson et al. Process Biochemistry 51 (2016) 1248–1255; Peng et al Biochemistry 2014, 53, 11, 1849–1857; Wang Bolen. Biophysical Journal Volume 71, Issue 4, October 1996, Pages 2117-2122; Eszes et al. FEBS Letters, Volume399, Issue3 December 16, 1996, Pages 193-197* |
| $K_{eq}$ | $13000 \pm 5000$ | | *eQuilibrator (37C, pH = 7.4, pMg = 3, I = 0.2M)* |

#### Monocarboxylate transporter (MCT)

*MCT Reaction:* Lactate_*cell*_ ⇄ Lactate_*media*_

*MCT genes:* Humans have four genes coding for MCT isoforms called SLC16A1-4. We focused on SLC16A1 (also MCT1) as it is the most abundant isoform in proliferating mammalian cells based on proteomics data (***Supplementary File 1***).

*MCT kinetic rate equation:* We used reversible Michaelis-Menten equation to describe activity of MCT.

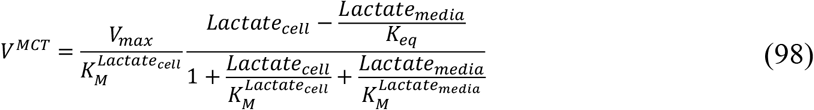

*MCT protein bound metabolite equation:*

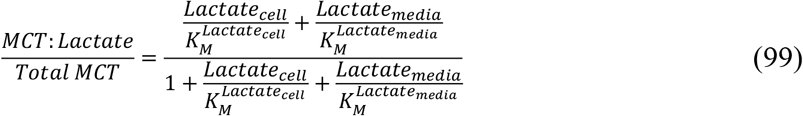

*Reported MCT regulation that was not included in our model:* MCT like all other transporters can exist in two conformations with lactate binding sites facing the media or facing the cytosol. The kinetic rate equation incorporating the two conformations will include another term in the denominator containing 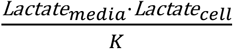. We did not include this additional term as we could not find data reporting the value of *K*. Addition of this term will slow down the MCT rate in the presence of high concentration of intracellular and media lactate. In addition, MCT can transport both lactate and pyruvate across the cell membrane. We have not included pyruvate transport by MCT, and lactate transport is dominant due to high concentration of lactate and inclusion of pyruvate transport will require addition of mitochondrial electron transport chain to maintain redox balance. Finally, MCT is a proton-coupled transporter of lactate so pH difference across plasma membrane will have an effect of MCT rate. Future models can incorporate these activities of MCT to improve the accuracy of predictions.

**Appendix 1–Table 14.** MCT kinetic Parameters. Kinetic parameters were used as is from references below without fitting the equation to data. *V*_*max*_ values for some publications were converted from *V*_*max*_ of cellular lactate uptake in cancer cells and erythrocytes. MCT level in cancer cells was taken to be 1.44•10^−4^ mg MCT1 per mg cellular protein as in ***Supplementary File 1***. To convert *V*_*max*_ values to 37C we used the temperature dependence of MCT *V*_*max*_ (***Carpenter and Halestrap, 1994***).

| Parameter | Mean ± SEM | Values | References |
| --- | --- | --- | --- |
| Molecular Weight, g/mol | 53,944 |  | UniProt: P53985 |
| $K_M^{Lactate_{media}} = K_M^{Lactate_{cell}}, \mu M$ | $5.5 \pm 0.9$ | 5.7, 5.34, 7.7, 4.68, 9.14, 2.99 | Ovens et al Biochem. J. (2010) 425, 523–530;<br>Carpenter et al Biochem J. (1994) 304,751-760;<br>Broer et al JBC. Vol. 272, No. 48, Issue of November 28, pp. 30096–30102, 1997;<br>Spenser Lehninger Biochem.J. (1976) 154,405-414;<br>Halestrap. Biochem.J.(1976)156,193-207;<br>Edlund Biochem.J.(1988)249,117-126; |
| $V_{max}, \frac{\mu mole}{min \cdot mg}$ | $470 \pm 230$<br>(Median ± MAD) | 469 (converted from $12.2 s^{-1}$ at 6C), 893 (converted from 0.065 $\mu mole/min$ per $\mu L$ cancer cell volume using 1.98 $\mu L/mg$ intracell volume to mg protein from same paper), 4722 (converted from 0.68 $\mu mole/min$ per mg cancer cell protein), 149, 124 (converted from 1.8, 1.5 nmole/min per mg 37C) | Ovens et al Biochem. J. (2010) 425, 523–530;<br>Carpenter et al Biochem.J.(1994) 304,751-760;<br>Spenser Lehninger Biochem.J. (1976) 154,405-414;<br>Guan et al. The AAPS Journal (2019) 21: 13; |
| $K_{eq}$ | 1 | Postulated to be 1 since transport is diffusion driven | |

#### Adenylate kinase (AK)

Adenylate kinase (AK) is not a glycolytic enzyme but it is abundant in most cells where it catalyzes interconversion of ATP, ADP and AMP. The function of AK is to ensure that AMP, which is a product of some energy consuming reaction but is not a substrate or product of glycolysis or respiration, can be converted back to ADP and ATP. We have added AK to the model to allow for more of the inorganic phosphate to be extracted from ATP. Addition of AK also allows for the formation AMP, which is an important regulator of some PFK isoforms, but it does not affect our model as it is not a regulator of PFKP at physiologically relevant concentrations.

*Reaction: 2ADP* ⇄ *ATP* + *AMP*

*AK kinetic rate equation:* We used a Rapid Equilibrium Bi Bi kinetic rate equation to describe AK activity:

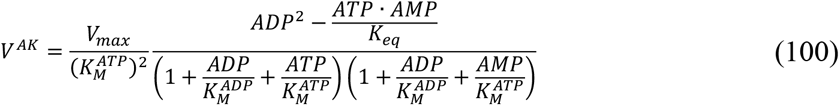

*AK kinetic parameters:* we choose the following *K*_*M*_ values based on the BRENDA database 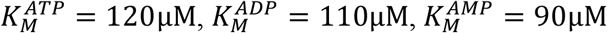 and *K*_*eq*_ = 0.48 based on eQuilibrator (***Beber et al., 2022***) value for AK reaction at *pH=7*.*4, pMg = 3* and *Ionic Strength = 0*.*2M. V*_*max*_ was set to be equal to the highest *V*_*max*_ · *Conc* of the model, which was for TPI.

#### Creatine kinase (CK)

Creatine kinase (CK) is not a glycolytic enzyme, but it is believed to be important for ATP homeostasis, especially in muscle cells where it is most abundant. We have added CK to the model simulations reported in ***Figure 6F*** and ***Figure 6–figure supplement 1E-G*** to investigate the role of CK in ATP homeostasis.

*Reaction: Creatine* + *ATP* ⇄ *Phosphocreatine* +*ADP*

*CK kinetic rate equation:* We used a Rapid Equilibrium Bi Bi kinetic rate equation to describe CK activity:

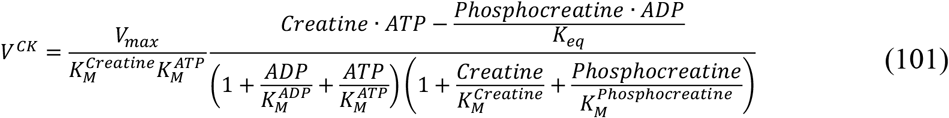

*CK kinetic parameters:* we used the following *K*_*M*_ values based on the BRENDA database 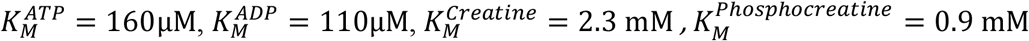 *and K*_eq_ *= 0*.*006* based on eQuilibrator (***Beber et al., 2022***) value for CK reaction at *pH=7*.*4, pMg = 3* and *Ionic Strength = 0*.*2M. V*_*max*_ was set to be equal to the highest *V*_*max*_ · *Conc* of the model, which was for TPI.

#### Dinucleotide Phosphate Kinase (NDPK)

Dinucleotide Phosphate Kinase (NDPK) is not a glycolytic enzyme, but it is believed to be important for ATP homeostasis. The function of NDPK is to phosphorylate NDP to NTP to allow the latter to be used in various biological processes, such as RNA transcription, glycogen synthesis, etc. We have added NDPK to the model simulations reported in ***Figure 6F*** and ***Figure 6–figure supplement 1E-G*** to investigate the role of NDPK in ATP homeostasis.

*Reaction: NDP* + *ATP* ⇄ *NTP* +*ADP*

*NDPK kinetic rate equation:* We used a Rapid Equilibrium Bi Bi kinetic rate equation to describe NDPK activity:

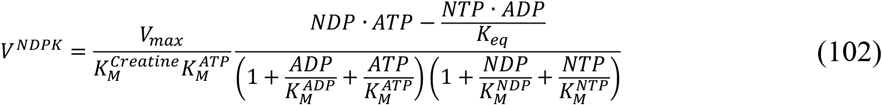

*NDPK kinetic parameters:* we used the following *K*_*M*_ values based on the BRENDA database 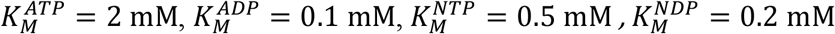 and *K*_*eq*_ *= 2*.*16* (average of 2.57 (UDP), 2.71 (GDP) and 1.21 (CTP)) based on eQuilibrator (***Beber et al., 2022***) value for NDPK reaction at *pH=7*.*4, pMg = 3* and *Ionic Strength = 0*.*2M. V*_*max*_ was set to be equal to the highest *V*_*max*_ · *Conc* of the model, which was for TPI.

#### ATPase

ATPase is included in our model as a mimic of cellular ATP consumption and is not a real enzyme. Activity (*V*_*max*_) of ATPase is varied in the model to study the effect of increased or decreased ATP consumption rate on model behavior. Typically, we set *V*_*max*_ of ATPase to be a fraction of the *V*_*max*_ · *Conc* of the slowest enzyme in the model (i.e., HK1), which corresponds to the pathway *V*_*max*_ as the whole pathway cannot proceed at a faster rate than its slowest enzyme.

*Reaction: ATP* + *H*_*2*_*O* ⇄*ADP* + *Phosphate*

*ATPase kinetic rate equation:* We used a Rapid Equilibrium Bi Bi kinetic rate equation to describe the combined effect of all ATP-consuming enzymes inside the cells:

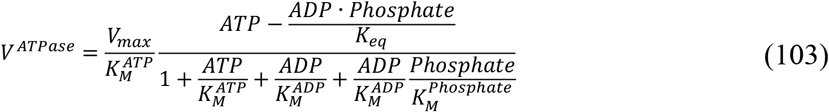

*ATPase kinetic parameters:* Since, the specific enzyme doesn’t exist we choose arbitrary values of 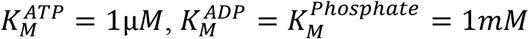 and *K*_*eq*_ *= 8*3,*00*0 was set based on eQuilibrator (***Beber et al., 2022***) value for ATP hydrolysis at *pH=7*.*4, pMg = 3* and *Ionic Strength = 0*.*2M. V*_*max*_ is set to different values in the model to study the effect of increased or decreased ATP consumption rate on model behavior.

